# Molecular phylogeny of the SELMA translocation machinery recounts the evolution of complex photosynthetic eukaryotes

**DOI:** 10.1101/2025.03.31.646294

**Authors:** Rafael I. Ponce-Toledo, David Moreira, Purificación López-García, Philippe Deschamps

## Abstract

Photosynthetic eukaryotes and their relatives are the result of an intricate evolutionary history involving a series of plastid acquisitions through endosymbiosis, multiple reversions to heterotrophy, and sometimes total plastid losses. Among these events, one of the most debated is the emergence and diversification of the CASH lineages (Cryptophyta, Alveolata, Stramenopiles and Haptophyta). Although they all include species bearing a complex plastid that derived from the endosymbiosis of a red alga, their phylogenetic relationships remain controversial, and the timing and number of plastid acquisitions are still undetermined. The inner metabolism of all plastids is mostly supported by nuclear-encoded proteins, and consequently, mechanisms allowing the relocation of those proteins have evolved or were recycled at each endosymbiotic event. Thus, the study of the composition and origins of those translocation machineries provides important clues for understanding how photosynthetic lineages have emerged and might be related. In CASH species, the SELMA complex, composed of about 20 proteins, is dedicated to the transport of pre-proteins across the periplastidial membrane, the second outermost membrane of secondary red plastids. In this work, we present the comprehensive genomic survey and phylogenetic analysis of the proteins composing the SELMA complex. We confirm the presence, homology and monophyletic origin of SELMA in the four CASH lineages and use these observations to infer a scenario for the serial transmission of secondary red plastids that differs from previous hypotheses and alters how the evolution of photosynthetic eukaryotes is envisioned.

## INTRODUCTION

Eukaryotes acquired the ability to carry out oxygenic photosynthesis in plastids more than 1.2 billion years ago thanks to primary endosymbiosis, namely the engulfment and assimilation of a cyanobacterium symbiont by a heterotrophic host (Archibald 2009; Parfrey et al. 2011). Three extant lineages have diversified from this founder event: Glaucophyta, Viridiplantae and Rhodophyta; composing the monophyletic supergroup Archaeplastida (Adl et al. 2012). Although the nature of the last common ancestor of the host lineage of Archaeplastida is still undetermined, phylogenetic analyses of the reduced genomes of primary plastids have allowed to trace their common ancestry to an extinct cyanobacterium related to Gloeomargaritales (Ponce-Toledo et al. 2017). Following primary endosymbiosis, photosynthesis was repeatedly transmitted to distant phyla of eukaryotes thanks to secondary and tertiary endosymbioses, during which various hosts associated with Archaeplastida symbionts, either red or green algae. For instance, Chlorarachniophyta (Rhizaria) and Euglenida (Excavata) both carry green alga derived plastids that originated from two independent secondary endosymbiosis events, involving symbionts related to Ulvophyceae and Prasinophyceae respectively (Rogers et al. 2007; Hrdá et al. 2012). Likewise, Cryptophyta, Alveolata, Stramenopiles and Haptophyta (termed thereafter CASH (Petersen et al. 2014)) include species with complex plastids related to red algae. Contrary to the green ones, the exact origins of red algae-derived complex plastids, but also the phylogeny and evolutionary histories of CASH lineages, have been controversial and are still a matter of strong debates. The long prevailing “Chromalveolate” hypothesis argued in favor of single endosymbiosis from which all CASH lineages would have derived. This was mostly supported by the similarity of intracellular membrane organizations between those phyla, considered an ancestral trait, but also by the idea that endosymbioses being complex evolutionary procedures, they should be considered as rare events (Cavalier-Smith 1999; Keeling 2009). Interestingly, the same kind of phylogenetic analyses that demonstrated the independent origins of green plastids have repeatedly showed that all Rhodophyta-derived secondary plastids are monophyletic (Yoon et al. 2002; Rodríguez-Ezpeleta et al. 2005; Baurain et al. 2010), supporting the Chromalveolate hypothesis. However, the accumulation of molecular data for protists’ nuclear genomes has offered new phylogenetic insights that systematically rejected the monophyly of the chromalveolates (Baurain et al. 2010; Burki et al. 2020). Thus, there is an incongruence opposing a seemingly vertical inheritance of plastids in CASH phyla and the absence of direct descent relationships between the nuclear genomes of the same groups. One way to reconcile these opposite patterns is to speculate that complex plastids were acquired via intricate serial endosymbioses (Bodył 2005; Sanchez-Puerta et al. 2007; Baurain et al. 2010; Petersen et al. 2014; Stiller et al. 2014; Strassert et al. 2020). In this scenario, a primordial secondary endosymbiosis gave rise to one of the four CASH phyla, which was later involved in one or several higher order endosymbioses, transmitting the complex plastid to other CASH lineages. This hypothetical evolutionary timeline is, however, still mainly built upon indirect evidence.

An emblematic feature of endosymbioses resides in the reduction of the symbiont genome and in the massive transfer of genes toward the host nucleus. These are called Endosymbiotic Gene Transfers (EGT) and have greatly modeled the genomes of photosynthetic eukaryotes (Martin et al. 1998). The genomes of Archaeplastida, for instance, contain a vast proportion of genes of cyanobacterial origin (Timmis et al. 2004). Similarly, genomes of species with complex plastids also enclose genes related to cyanobacteria as well as many eukaryotic genes that they obtained from their algal symbiont (Paper et al. 2000; S.B. Gould et al. 2006; Li et al. 2006). A consequence of EGT is that most of the proteins engaged in plastid metabolisms are encoded in the nuclear genome and must be relocated into the plastid to perform their function. Primary plastids are surrounded by two membranes which contain a protein import complex called “Tic/Toc” - for Translocators of the Inner and Outer Chloroplast membranes (Sjuts et al. 2017). Nuclear encoded plastidial pre-proteins present a particular N-terminal sequence called the transit peptide that triggers their recognition and addressing to the Tic/Toc complex for translocation. This transit peptide is subsequently cleaved and mature proteins are released in the stroma or sent to the thylakoids. Complex plastids derive from photosynthetic green or red algae and are consequently surrounded by more membrane layers. Plastids of CASH species (except Dinophyta) possess four membranes, being, from the outside to the inside: the outermost membrane (OM); the periplastidial membrane (PPM), and finally the original plastid double membrane (PM) (fig. 1a). Dinophyta have lost the PPM secondarily. In Cryptophyta, Haptophyta and Stramenopiles, the OM is continuous with the outer nuclear membrane and populated with ribosomes, hence the plastid resides in the endoplasmic reticulum lumen (or Chloroplast Endoplasmic Reticulum; CER; fig. 1a (Gibbs 1979)). In Alveolata, the plastid is located in the cytoplasm. Between the PPM and the PM lies the periplastidial space (PPS), corresponding to the former cytoplasm of the algal symbiont (fig. 1a). In Cryptophyta, the PPS encloses a nucleomorph (fig. 1a), which is a relic of the symbiont’s nucleus (Greenwood 1974; Hibberd and Norris 1984; Maier 1992) and contains a vestigial genome composed of 300 to 450 genes mainly involved in its own replication and expression (Maier et al. 2000; Zauner et al. 2019). In summary, pre-proteins that have to be relocated to complex red plastids must pass through three to four membranes. Several molecular studies have shown that, in all lineages with a complex plastid, nuclear encoded pre-proteins are prefixed by a bipartite N-terminal topogenic signal (BTS); *i.e.* a combination of a signal peptide and a transit peptide. The relocation of pre-proteins is a two-step process involving the translocation into the secretory pathway followed by the import into the plastid stroma through Tic/Toc homologs. Both the signal peptide and the transit peptide are cleaved after preprotein translocation into the CER and into the stroma, respectively. (Grossman et al. 1990; Bhaya and Grossman 1991; Sulli et al. 1999; Waller et al. 2000; Rogers et al. 2004; S. Gould et al. 2006).

**FIG 1.**
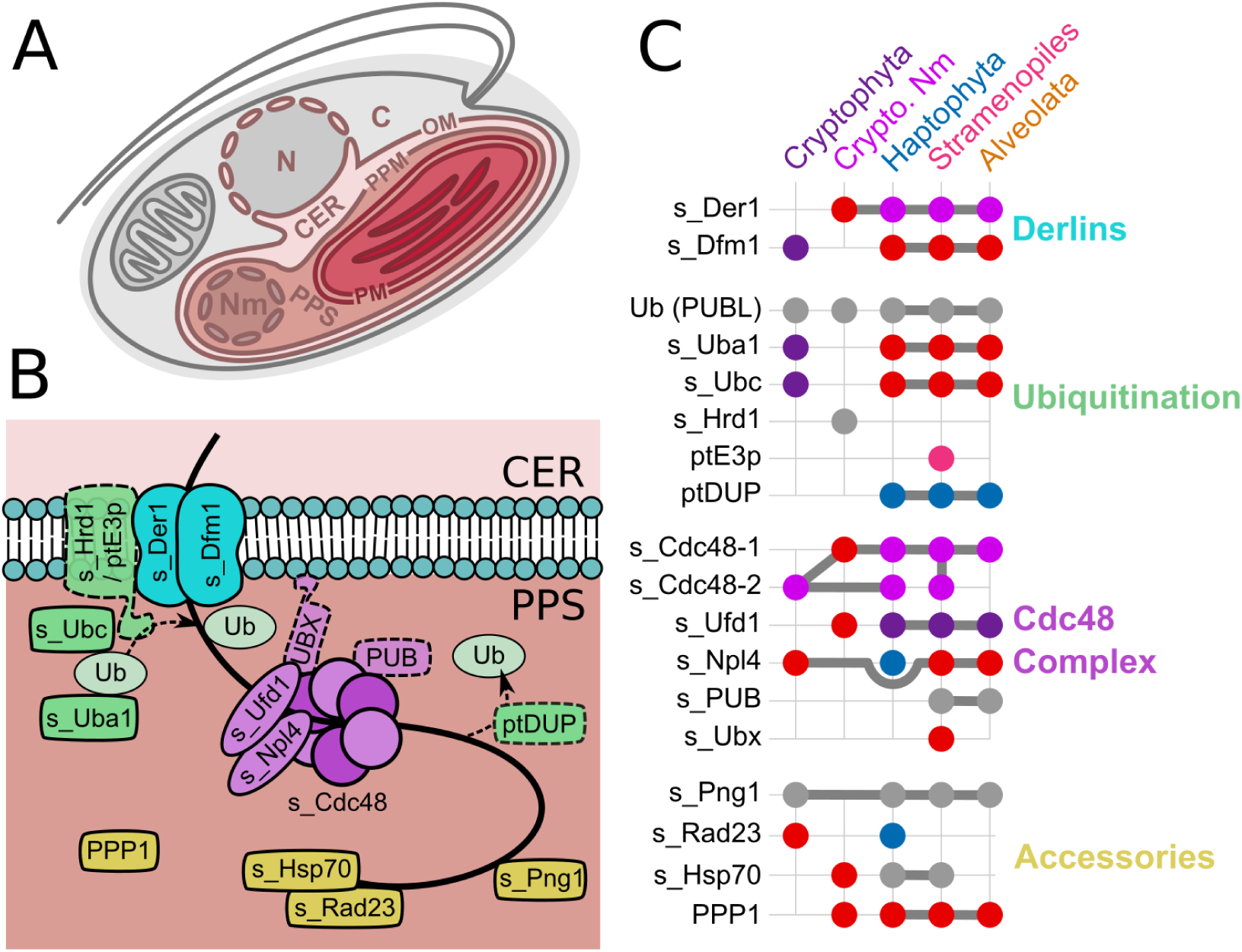
Composition of the SELMA transporter across CASH lineages. *(A)* Schematic organization of a photosynthetic Cryptophyta cell depicting organelles and membranes. N: nucleus, Nm: nucleomorph, C: cytoplasm, OM: plastid outer membrane, CER: chloroplast endoplasmic reticulum, PPM: periplastid membrane, PPS: periplastidial space, PM: plastid double membrane. *(B)* Schematic structure of the SELMA transporter, adapted from Lau et al (2016). Functional categories are annotated using different colors; light blue: Derlins, green: ubiquitination, purple: Cdc48 complex, yellow: accessory proteins. Please refer to the main text for details about the name and role of SELMA constituents. *(C)* Presence/absence, origin and phylogenetic relationships of putative SELMA components in CASH lineages. For each protein (line), a large dot indicates its existence in a lineage (column). The color of the dot corresponds to the inferred origin of the gene; dark purple: Cryptophyta nucleus, light purple: Cryptophyta nucleomorph, blue: Haptophyta, pink: Stramenopiles, brown: Alveolata. When two dots are linked with a thick line, the corresponding proteins share a common ancestor.

The nature of the transporter driving proteins across the PPM was unveiled much more recently. Its discovery resulted from the detection of a peculiar Endoplasmic Reticulum Associated Degradation (ERAD) complex encoded in the nucleomorph genome of Cryptophyta as well as in the nuclear genome of Apicomplexa and Stramenopiles (Sommer et al. 2007). ERAD is a ubiquitin dependent pathway used by all eukaryotic cells to extract miss-folded proteins from the endoplasmic reticulum back into the cytoplasm where they can be degraded by the proteasome (Meusser et al. 2005). Sommer et al (2007) speculated that this newly-found plastid-located additional copy of ERAD could have been inherited from red algae by EGT and tinkered to create a translocon that imports proteins across the PPM. The putative new complex was named SELMA for Symbiont-specific Erad-Like MAchinery (Hempel et al. 2009); its structure has been dissected and evidence supporting its function were produced both by *in silico* studies (Felsner et al. 2011; Moog et al. 2011; Stork et al. 2012) and by functional experiments (Agrawal et al. 2009; Hempel et al. 2009; Hempel et al. 2010; Agrawal et al. 2013; Lau et al. 2015; Fellows et al. 2017). Currently, SELMA is hypothesized to function using approximately 20 proteins that are, for the most part, shared by all available sequenced genomes of CASH species having a red plastid (Table 1, (Stork et al. 2012)). The discovery of this PPM protein translocator shared by all CASH and likely inherited by EGT from Rhodophyta brought new arguments into the debated question of the evolution of complex red plastids. Indeed, it seems very unlikely that the same multi-protein transporter evolved several times after independent secondary endosymbioses with different red algae. This idea is strengthened by the observation that, in Chlorarachniophyta, an evolutionary distinct green complex plastid containing lineage which also possesses a PPM, a PPS and a nucleomorph, there is no detectable SELMA-like complex (Hirakawa et al. 2012). The fact that SELMA components in Cryptophyta are partly encoded in the nucleomorph (thus undoubtedly inherited from secondary endosymbiosis), and partly encoded in the nucleus, prompted some authors to speculate that Cryptophyta represent an evolutionary intermediate of secondary red plastid integration (Zimorski et al. 2014; Gould et al. 2015; Cavalier-Smith 2018). Additionally, in the very restricted set of single protein phylogenies available in the literature, SELMA components of Alveolata, Stramenopiles and Haptophyta form a clade that is a sister group to the homologous protein of Cryptophyta (Felsner et al. 2011; Fellows et al. 2017). Those clues were used to argue that ancestors of Cryptophyta were likely the symbionts involved in the endosymbiotic series that gave rise to other photosynthetic CASH phyla. However, no systematic phylogenetic analysis of the entire SELMA complex was ever attempted. Such an analysis could assert if genes involved in SELMA were indeed commonly inherited from red algae; if they were transferred between CASH phyla, and if some of those transfers might indicate the timing and order of complex endosymbioses. In this work, we attempted to reconstruct phylogenies of all described protein components of the SELMA complex. We confirm the existence and homology of SELMA in all CASH phyla and propose an interpretation of the phylogenetic tree topologies in the context of complex red plastid evolution.

## RESULTS

### Data mining and phylogenetic analysis of SELMA-related proteins

Stork et al. (2012) gathered, using sequence similarity searches, a comprehensive sequence collection of putative SELMA components from all CASH genomes available in 2012. To differentiate SELMA from ERAD proteins, they used *in silico* predictions of BTS as well as BLAST dissimilarity as proxy of evolutionary distance between sequences. Members of the SELMA complex can be classified into four functional categories that will be fully described thereafter: (1) the Derlin pore, (2) the ubiquitination machinery, (3) a Cdc48-Ufd1-Npl4 complex and (4) a set of accessory proteins (fig. 1b). For the sake of clarity, the following terminology will be used throughout the rest of the manuscript: genes and proteins putatively involved in ERAD will have the prefix “e_” while those potentially involved in SELMA will be prefixed “s_”. We used protein sequence identifiers from supplementary table S1 in Stork et al. as a starting set for our own data mining procedure. We assembled a custom database containing protein sequences predicted from 864 genomes and transcriptomes of species distributed across the 3 domains of life with an emphasis on photosynthetic eukaryotes (supplementary table S3). We used the starting set to query our database using reciprocal BLASTP (Altschul et al. 1990) in order to retrieve as many non-redundant similar protein sequences as possible for each potential ERAD/SELMA component. We then used those collections of similar proteins to reconstruct and refine phylogenetic trees and managed to obtain informative phylogenies for 16 of the 21 putative SELMA components analyzed (supplementary table S1). SELMA is considered to have evolved from the refunctionalization of ERAD genes of Rhodophyta obtained by EGT (Bolte et al. 2011). Therefore, it is expected that eukaryotes lacking a complex plastid should only have one copy of each ERAD components while photosynthetic CASH genomes should carry at least 2 copies: one involved in the cytosolic/endoplasmic ERAD complex and the other one in the plastid-located SELMA complex. We looked for the corresponding patterns in our trees and included *in silico* predictions of cellular localization to determine which paralogous group of proteins might be implicated in SELMA or in ERAD. By doing so, we could detect several classification errors in Stork et al. study, probably due to the fact that they used BLAST “distances” in lieu of phylogenetic relationships, which is known to be unreliable (Koski and Golding 2001). We provide an updated table of ERAD/SELMA protein references in supplementary Table S2. We describe thereafter the results we obtained for each of the functional groups composing the SELMA transporter.

### The SELMA Derlin translocon likely originated by EGT from Rhodophyta

There is still an open debate about the nature of the pore through which proteins are excreted out of the ER when they are processed by ERAD. For some authors, Derlin transmembrane proteins, which are present in all eukaryotes, are the main constituent of this channel (Rao et al. 2021). Those proteins have been extensively studied in the model *Saccharomyces cerevisiae*, whose genome encodes two Derlin paralogs, namely Der1 (for “degradation in the ER”) and Dfm1 (for “Der1-like family member”) (Knop et al. 1996; Hitt and Wolf 2004). Analyses of yeast single mutants for each isoform indicate that Der1 is mainly implicated in the retrotranslocation of luminal proteins (ERAD-L) while Dfm1 would rather transport membrane integral proteins (ERAD-M) (Neal et al. 2018). It seems, however, that each isoform is able to compensate for the absence of the other, explaining the lack of altered growth phenotype in single mutant lines (Knop et al. 1996; Hitt and Wolf 2004). Additionally, double mutants impaired for both Derlins are able to grow, indicating that there might also be alternative ways to export misfolded proteins out of the ER (the E3 ubiquitin ligase Hrd1 is one candidate (Wu and Rapoport 2018)). Although all heterotrophic eukaryotes (and Archaeplastida) only have one pair of Derlins dedicated to ERAD, additional genes encoding Der1 and Dfm1 homologs have been detected in all photosynthetic CASH lineages (Sommer et al. 2007; Spork et al. 2009; Felsner et al. 2011). Interestingly, both are absent in the apicoplast-lacking parasite *Cryptosporidium*, suggesting a specific role of these proteins in the plastid (Agrawal et al. 2009). The gene coding s_Der1 was first detected in the nucleomorph genome of *Guillardia theta* and used to successfully complement the yeast *der1* mutant, indicating a similar function (Sommer et al. 2007). s_Der1 and s_Dfm1 proteins are encoded in the nuclear genome of all other CASH and were proven to be located in the PPM in Apicomplexa and Stramenopiles (Hempel et al. 2009; Kalanon et al. 2009; Spork et al. 2009). A deletion of the nuclear gene *s_der1* in *Toxoplasma gondii* hinders the relocation of proteins into the apicoplast, leading eventually to the death of the parasite (Agrawal et al. 2009), definitely proving the role of s_Derlins in SELMA.

To our knowledge, three publications have reported phylogenetic trees of the s_Derlin proteins (Hirakawa et al. 2012; Petersen et al. 2014; Cavalier-Smith 2018). However, they all suffer from poor taxon sampling or miss many appropriate orthologs. We reconstructed several wide sampled phylogenetic trees of Der1+Dfm1, which allowed us to confirm that all CASH lineages possess both s_Der1 and s_Dfm1, but also to determine that e_Dfm1 and e_Der1 are ancient paralogs that were likely duplicated in the Last Eukaryotic Common Ancestor. We present in supplementary fig. S1 a phylogenetic tree rooted where we believe is the correct split between Dfm1 and Der1 paralogs. We observed that the internal topology of each paralog is unstable, probably because of long branch attraction (LBA) artifacts when including both paralogs in the same tree. Hence, we split the dataset into two sub-trees (fig. 2; supplementary fig. S2 and S3). Our phylogenetic tree of Der1 supports for the first time that *s_der1* was inherited by serial EGT (fig. 2a and supplementary fig. S2). Similarly, we observe that s_Dfm1 proteins of Alveolata, Stramenopiles and Haptophyta form a single monophyletic group that is sister to Rhodophyta, although with low bootstrap support (fig. 2b and supplementary fig. S3). This indicates that *s_dfm1* in these phyla likely derives from *e_dfm1* of Rhodophyta and was inherited by EGT. Dfm1 homologs in Cryptophyta are, however, found as two paralogous sister clades, one being nested in the other and having a root branch significantly longer. The plastid lacking species *Goniomonas* sp. seems to harbor only one of the two Dfm1 paralogs. We suggest that the short-branching group corresponds to e_Dfm1 and that the long-branching one is likely s_Dfm1 that evolved by duplication and replaced a putative protein derived originally from red alga. We observed this pattern for other SELMA components and will discuss it afterwards.

### Genes involved in the ubiquitylation of SELMA-translocated proteins have conflicting origins

Ubiquitin (Ub) is a protein generally composed of 76 amino-acids, whose sequence is extremely conserved among eukaryotes (Swatek and Komander 2016). Ubiquitination is a post-translational modification consisting of the covalent addition of ubiquitin moieties onto lysine residues of targeted proteins. This modification plays a pivotal role in signaling and regulation of many metabolic pathways of the eukaryotic cell (Komander and Rape 2012; Swatek and Komander 2016). During the ERAD process, ubiquitination of misfolded proteins on the cytosol side is essential to their Derlin/Hrd1 mediated extrusion as well as their relocation to the proteasome (Claessen et al. 2012; Lemus and Goder 2014). Ubiquitination is performed thanks to the consecutive action of three enzymes: Ub activating enzyme E1, Ub conjugating enzyme E2 and Ub ligase E3. Ubiquitin activation is handled by the versatile enzyme Uba1 that uses ATP to first adenylate Ub and subsequently transfers it to one of its cysteine residues, forming an E1∼Ub complex (McGrath et al. 1991). Ubiquitin is then transferred to the active site of one of the many ubiquitin conjugating E2 enzymes. Finally, a ubiquitin protein ligase (E3) interacts with both E2∼Ub and the final protein substrate to transfer Ub to a lysine residue of the latter. The nature of the E2/E3 couple determines the specificity of ubiquitination toward the substrate and differentiates ubiquitin-using pathways. In the case of ERAD, this couple consists predominantly of Ubc7 and Hrd1, with Hdr1 catalyzing the transfer of ubiquitin to misfolded proteins at the ER luminal side, but also regulating the extrusion process by self-ubiquitination (Baldridge and Rapoport 2016). The Ubiquitin moieties are then recognized by Ub-interacting chaperones and cofactors to address misfolded proteins to the proteasome.

SELMA transport across the PPM also requires the ubiquitination of precursor proteins on the PPS side. A specific Plastidial UBiquitin Like protein (PUBL) has been identified and its plastidial localization experimentally confirmed in *Phaeodactylum tricornutum*, *Plasmodium falciparum* and *Toxoplasma gondii* (Sommer et al. 2007; Spork et al. 2009; Fellows et al. 2017). The specific implication of PUBL in SELMA has been demonstrated in a conditional mutant of *T. gondii* (Fellows et al. 2017) whose phenotype displays a decrease in plastid-located mature proteins and eventually induces the death of the parasite. PUBL proteins are longer than Ub, with extensions on the N-terminal side (comprising the BTS), but also on the C-terminal side, which eliminates the regular Ub C-terminal glycine and seems incompatible with polyubiquitination. A published phylogenetic tree of PUBL proteins, comprising conventional Ub sequences as outgroup, showed with moderate support that PUBL is shared by all photosynthetic CASH (excluding Cryptophyta) and constitutes a monophyletic group, indicating a common ancestry (Fellows et al. 2017). We could reconstruct an equivalent tree (supplementary fig. S4a) using only the conserved positions of Ub and PUBL and with the addition of PUBL sequences of *Plasmodium* species that were originally excluded by Fellows et al. (2017). In our tree, those proteins are found nested within the Apicomplexa, with a very long basal branch, indicating a strong acceleration of the evolutionary rate of their coding genes. Although *Plasmodium* PUBL sequences are highly divergent, it appears that structural models predicted by Alphafold can be successfully aligned with those of *Toxoplasma*. (Jumper et al. 2021; Varadi et al. 2022) (supplementary fig. S4b). Contrary to previous observations, our survey of PUBL similar proteins did identify homologs encoded in the nucleomorph genome, but only for some Cryptophyta species (*Chroomonas mesostigmatica*, *Cryptomonas curvata* and *Hemiselmis andersenii*). Indeed, *Rhodomonas abbreviata*, *Geminigera cryophila* and *G. theta* share a nuclear encoded Ub homolog with a N-terminal BTS extension, suggesting that those species could have replaced a nucleomorph encoded Ub with a nucleus encoded plastid targeted PUBL. However, probably because of insufficient positions in the alignment, our tree does not support an evolutionary relationship between nucleomorph-encoded Ub, nuclear-encoded Ub-like proteins and PUBL, whose exact origin remains undetermined.

SELMA homologs of Uba1 have been reported on several occasions (Sommer et al. 2007; Spork et al. 2009; Felsner et al. 2011; Stork et al. 2012). A *s_uba1* gene can be found in all photosynthetic CASH nuclear genomes and the PPS localization of s_Uba1 was demonstrated experimentally in *Plasmodium* (Spork et al. 2009). A phylogenetic tree of e_Uba1 and s_Uba1 proteins is available in Felsner et al. (2011) where CASH s_Uba1 sequences form a poorly supported monophyletic group that also comprises the unique red alga *Cyanidiochyzon merolae*. Our own phylogenetic tree of Uba1 proteins shows a more complex evolutionary history (supplementary fig. S5). First, the tree displays evident signs of an ancient paralogy with differential losses. For instance, Rhodophyta and Glaucophyta share an ortholog that differs from the one found in Viridiplantae and Alveolata. Moreover, Haptophyta and Cryptophyta possess two copies of e_Uba, one related to the isoform of Viridiplantae/Alveolata and the second shared with all other eukaryotes. We also observe an additional homolog of Uba1 shared by all photosynthetic CASH, except Cryptophyta, and forming a monophyletic group that branches in between the two previously described paralogous groups. Proteins of this clade have putative BTS sequences and are likely SELMA s_Uba1 enzymes. Interestingly, this group also contains orthologous proteins of Chlorarachniophyta species, suggesting the existence of plastid-located ubiquitin using pathways in this phylum. Finally, we observe in Cryptophyta the same topology as for s_Dfm1: a specific duplication with one paralog being e_Uba1 (found also in the plastid-lacking species *Goniomonas* sp.), and the other one being likely s_Uba1.

The main experimental evidence of the involvement of a specific E2 gene in SELMA was published by Agrawal et al (2013), who showed that a *T. gondii* mutant line affected at the *TGME49_295990* locus was unable to import proteins in the apicoplast. E2 ubiquitin conjugating enzymes compose a very large family of orthologous genes, each having one or many E3 partners, and acting at different areas of the cell metabolism (Michelle et al. 2009). For instance, the genome of *S. cerevisiae* contains 13 E2 *ubc* genes; *C. elegans* encodes 20, and *A. thaliana* has no less than 37 (Jones et al. 2002; Kraft et al. 2005; Finley et al. 2012). Unfortunately, their naming is not coherent between organisms and their classification is misleading (Michelle et al. 2009). Additionally, it is hard to determine if putative SELMA E2 enzymes that were previously detected in the literature using sequence similarity are in fact all related and orthologous to the experimentally evidenced gene *TGME49_295990* (Sommer et al. 2007; Spork et al. 2009; Hempel et al. 2010; Stork et al. 2012). To overcome this issue, we used every *S. cerevisiae* Ubc protein sequence to search our database for homologs and combined all non-redundant hits to produce a general tree of E2 enzymes. Fig. 3a presents this general tree where each isoform group is delimited and named based either on *S. cerevisiae* numbering (Ubc+N) or using *Arabidopsis thaliana* E2 genes naming convention (Atg+N) (Kraft et al. 2005). The position of all nucleomorph-encoded Cryptophyta E2-like proteins is depicted: they fall into 3 families: Atg9-Ubc10/Pex4, Atg6/13-Ubc4/5 and Atg3/Ubc2, which are respectively described as involved in peroxisome biogenesis and functioning; protein degradation / anaphase-promoting complex; and the N-end rule pathway / DNA repair (Finley et al. 2012). Two other sub-trees contain proteins found encoded in CASH genomes and which are predicted to be targeted to the plastid: Atg14.1 comprising a clade of Stramenopiles; and Atg7 which contains all E2 proteins previously reported as potentially involved in SELMA, including the product of *TGME49_295990*. Fig. 3b and supplementary fig. S6 focuses on this Atg7 sub-group and show that photosynthetic CASH E2-like proteins (excluding Cryptophyta) form a monophyletic group closely related to Rhodophyta, suggesting an acquisition by EGT. Additionally, we could again observe a pattern of specific duplication in Cryptophyta, with one paralog having a longer basal branch and possibly being s_Ubc.

**FIG 2.**
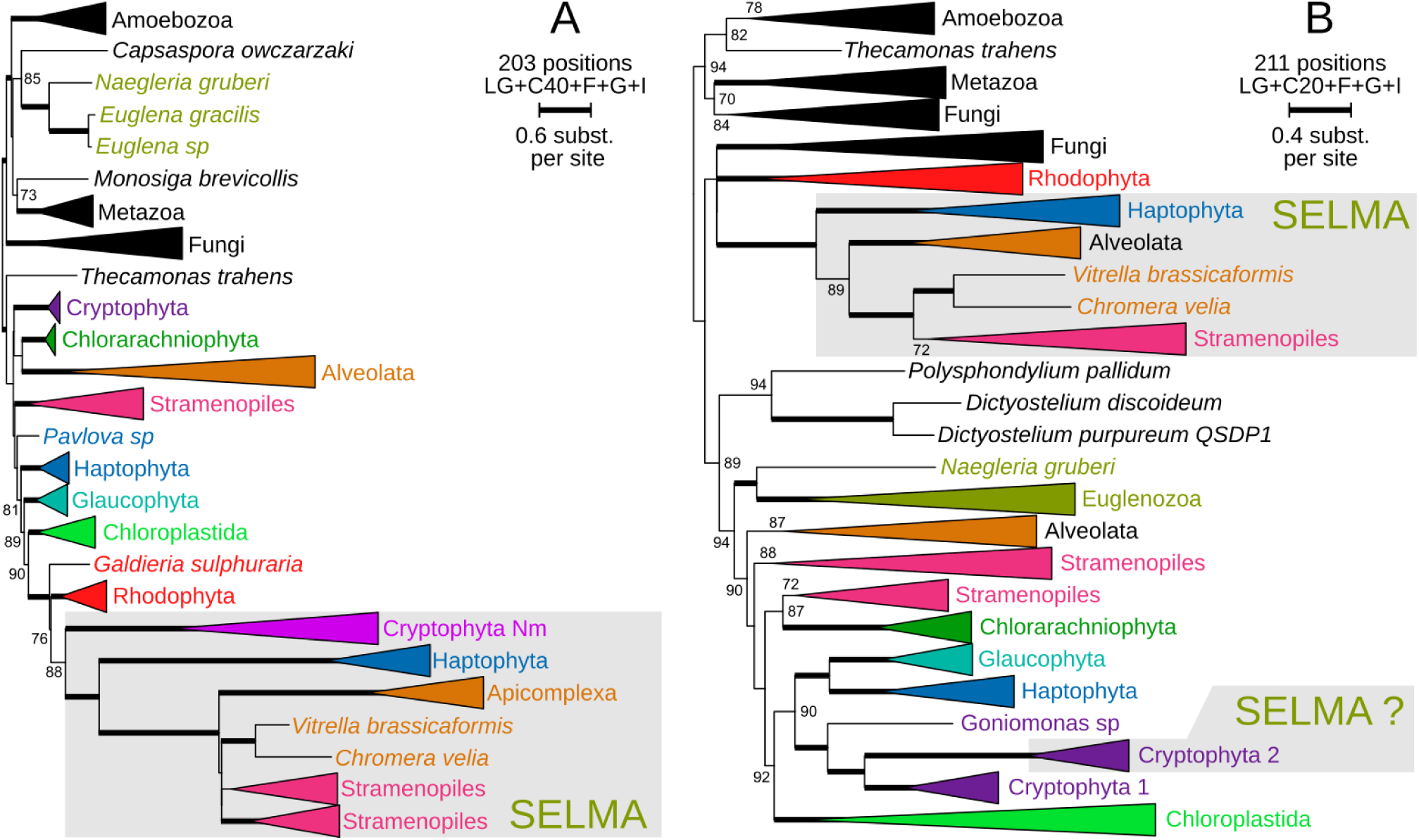
Unrooted condensed maximum likelihood phylogenetic trees of Der1 *(A)* and Dfm1 *(B)* proteins. The corresponding expanded trees are available in supplementary fig. S2 and S3 respectively. For the sake of readability, monophyletic phyla are depicted as triangles. Ultrafast bootstrap branch support values (1000 replicates) are indicated except when lower than 70% or as a thick branch when the value is maximum. The number of amino acid positions in the corresponding sequence alignment, the substitution model and parameters used for IQ-TREE reconstruction as well as the branch length scale are indicated at the top right of each tree. Clades corresponding to proteins putatively implicated in SELMA are shadowed in gray.

**FIG 3.**
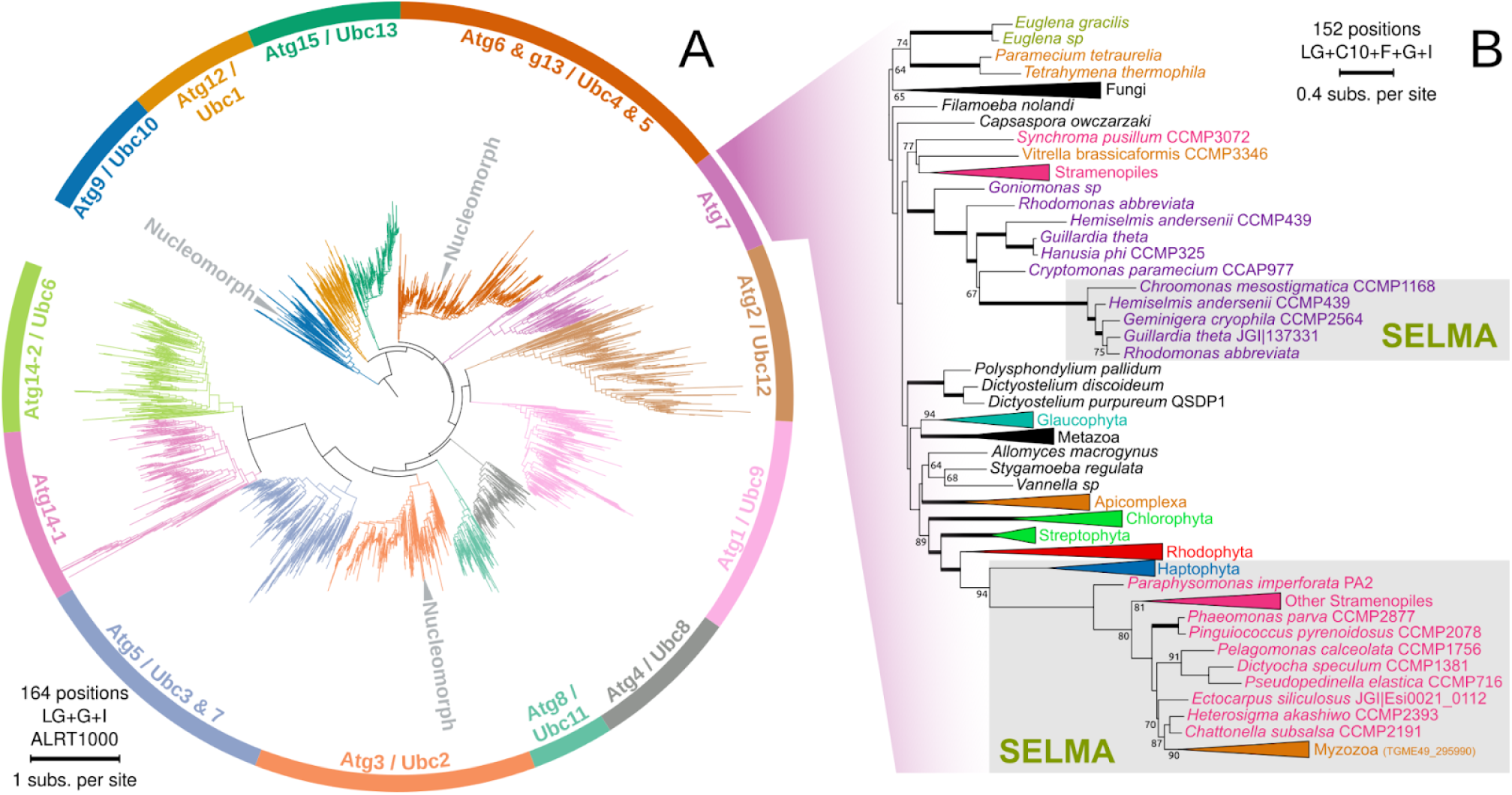
Phylogenetic analysis of the E2 Ubiquitin conjugating (Ubc) enzymes. The number of amino acid positions in the corresponding sequence alignment, the substitution model and parameters used for IQ-TREE reconstruction as well as the branch length scale are indicated for each tree *(A)* Maximum likelihood phylogenetic tree of the whole Ubc protein family. The tree is arbitrarily rooted. Each isoform is depicted with a different color and named using the convention for *S. cerevisiae* (UbcN) and *A. thaliana* (AtgN). The positions of nucleomorph encoded Ubc enzymes are indicated in gray. *(B)* Unrooted condensed maximum likelihood phylogenetic tree focusing on Ubc enzymes of the Atg7 family. Monophyletic phyla are depicted as triangles. The corresponding expanded tree is available in supplementary fig. S6. Ultrafast bootstrap branch support values (1000 replicates) are indicated except when lower than 70% or as a thick branch when the value is maximum. Clades corresponding to proteins putatively implicated in SELMA are shadowed in gray.

The E3 component of the potential SELMA ubiquitination cascade is more elusive. In ERAD, ubiquitin transfer is performed by e_Hrd1, which also regulates the process of protein extrusion. A SELMA equivalent to Hrd1 has been repetitively reported to exist only in Cryptophyta (Sommer et al. 2007; Spork et al. 2009; Hempel et al. 2010). We were also unable to detect any orthologs of *s_hrd1* in other CASH genomes. Our attempts at reconstructing the phylogeny of Hrd1 proteins recovered nucleomorph-encoded s_Hrd1 nested in the Archaeplastida supergroup but not as a sister group to Rhodophyta (supplementary fig. S7). Our results also invalidate the candidate s_Hrd1 protein of *Emiliania huxleyi* that was reported by Stork et al (2012). The absence of s_Hrd1 in most CASH lineages is unexpected and some authors have tried to identify alternatives by screening for proteins having trans-membrane domains attached to an E3-ligase module. The ptE3p protein of the diatom *P. tricornutum* has been proposed as an equivalent to s_Hrd1 in Stramenopiles. This protein has a verified E3-ligase activity and seems to be an actual physical component of the purified SELMA complex of *P. tricornutum* (Hempel et al. 2010). We gathered homologs of ptE3p and tried to reconstruct their phylogeny but all our trees showed signs of a weak signal leading to unresolved topologies (data not shown). Nevertheless, this protein seems to only exist in Stramenopiles, which means that, if it is indeed involved in SELMA, it would not give any information about the origin of the specific E3 enzyme of the putative common ancestor of CASH plastids.

Although the fate of ubiquitinated proteins in ERAD is their destruction, proteins translocated through SELMA should not be degraded, and it seems logical to suppose that PPS-located and stromal proteins would be de-ubiquitinated to function properly. Hempel *et al*. (2010) queried the genome of *P. tricornutum* for a putative PPS located deubiquitinating enzyme and proposed one candidate that they named ptDUP. We collected homologs of the ptDUP protein encoded in CASH genomes and reconstructed the corresponding phylogenetic tree (supplementary fig. S8). Proteins similar to this enzyme are present in most eukaryotic genomes, including *S. cerevisiae* (named Ubp15p and involved in the peroxisomal export machinery (Debelyy et al. 2011)) and *A. thaliana* (named UBP12 and 13 and involved in several regulation processes (Cui et al. 2013; Lindbäck et al. 2022)). Our tree shows the existence of a monophyletic paralogous group containing ptDUP as well as similar proteins (with predicted BTS) of Haptophyta, Stramenopiles and *Vitrella brassicaformis* (Chromerida) but not of Cryptophyta nor Apicomplexa. The closest sister group to this ptDUP clade is not Rhodophyta but Haptophyta (with very low support) which, if not erroneous or due to reconstruction artifacts, may be an interesting observation that will be discussed below.

Overall, the SELMA-specific ubiquitination pathway seems to have a monophyletic ancestry in Haptophyta, Stramenopiles and Apicomplexa, but the initial origin of each element appears contradictory (fig. 1c). Additionally, most of the corresponding proteins in Cryptophyta seem to have evolved from ERAD components via ancient phylum specific duplications that are not related to proteins of the other CASH lineages. We will discuss below how these replacements might affect our analysis.

### The SELMA Cdc48-Ufd1-Npl4 complex was likely inherited from Rhodophyta

Cdc48 (p97 in mammals) belongs to the AAA protein family (ATPases Associated with diverse cellular Activities), whose members consume ATP to perform mechanical actions on macromolecules (Bodnar and Rapoport 2017). More precisely, Cdc48 achieves the physical displacement of ubiquitinated proteins in various cellular processes, with a specificity that depends on the nature of its cofactors (Bays and Hampton 2002). In the case of ERAD, Cdc48 associates with the couple Ufd1 (ubiquitin fusion degradation) / Npl4 (nuclear pore localization). Together, they pull misfolded ubiquitinated proteins out of the ER and transfer them to the proteasome. A single SELMA-putative copy of Cdc48 was initially detected in the nucleomorph genome of Cryptophyta and in the nuclear genome of other CASH species (Sommer et al. 2007). Phylogenetic analyses have shown that those s_Cdc48 proteins cluster in a monophyletic group directly related to Rhodophyta (Agrawal et al. 2009; Felsner et al. 2011; Petersen et al. 2014). A deeper mining of genomic data uncovered a second copy of *s_cdc48* in all CASH except in Alveolata. s_Cdc48-1 and sCdc48-2 are both targeted to the plastid in *P. tricornutum* where they physically interact (Spork et al. 2009; Lau et al. 2016). Thus, contrary to ERAD, the Cdc48 complex involved in SELMA is likely a heterohexamer. Our own survey confirms the existence of both isoforms of s_Cdc48 in all photosynthetic CASH except Alveolata, which only have one. Additionally, our phylogenetic analysis demonstrates that s_Cdc48-1 and s_Cdc48-2 are paralogs that derive from the duplication of a single protein originally inherited from Rhodophytes (fig. 4a and supplementary fig. S9). However, the topology of the tree is not compatible with an early duplication of isoforms 1 and 2 before the diversification of CASH but rather with a more complex history of lineage-specific duplications. Lau et al. (2016) described in *P. tricornutum* a specific motif (DDDLYS*) present at the C-terminal side of s_Cdc48-1 but absent from s_Cdc48-2. We also observe this motif on many s_Cdc48 proteins, including some encoded in the nucleomorph of Cryptophyta and in Rhodophyta, which indicate, given the topology of the tree, that the presence of this motif was the ancestral state of s_Cdc48. Additionally, there seems to be a correlation between the pattern of presence/absence of the motif and the delimitation of isoform 1 and isoform 2 in our phylogenetic tree (marked with green Ⓜ signs in supplementary fig. S9). However, if the topology of the tree reflects the true history of the gene, the loss of the motif would then be a convergence. SELMA specific copies of Npl4 and Ufd1 also exist in the genomes of photosynthetic CASH species. In Cryptophyta, *s_npl*4 is found in the nuclear genomes, while *s_ufd1* is encoded in the nucleomorph genome. Our phylogenetic analyses show that both genes in Cryptophyta originated from Rhodophyta genes via secondary EGT (fig. 1c, fig. 4b and 4c, supplementary figs. S10 and S11). Interestingly, s_Ufd1 in Stramenopiles, Apicomplexa and Haptophyta derive from the same ancestral protein but does not seem to be related to the nucleomorph-encoded protein of Cryptophyta, but rather to e_Ufd1 (fig. 4c, supplementary fig. S11). The case of s_Npl4 is slightly different: the corresponding gene has a common origin in Stramenopiles and Alveolata, and the group is related to Rhodophyta and Cryptophyta, indicating a possible inheritance by EGT (fig. 4b, supplementary fig. S10). In Haptophyta, s_Npl4 is likely a specific duplication of e_Npl4, as it displays a similar topology than what we could observe for several proteins of the Ubiquitination pathway in Cryptophyta. Finally, we also analyzed two putative, PPS located, additional components of the Cdc48 complex, s_UBX and s_PUB, that might be respectively involved in its interaction with Derlins or in its regulation (Spork et al. 2009; Moog et al. 2011; 2016). s_PUB is composed of a combination of a thioredoxin domain and a PUB domain (supplementary fig. S12a). After a thorough search for similar sequences in our database, we observed that only Stramenopiles and Chromerida (Alveolata) possess a protein with this combination of domains. The closest similar proteins are thioredoxins of the Ybbn family: they present a similar thioredoxin domain but are otherwise completely different in structure. We reconstructed a phylogenetic tree using only the portions that align (supplementary fig. S12b). This tree shows that s_PUB sequences cluster in a monophyletic group that is nested into a group of Ybbn-like proteins of Stramenopiles. Thus, s_PUB is a specific innovation that is only shared by Chromerida and Stramenopiles, and might play a role that does not exist in other CASH. On the other hand, s_UBX is related to a widely distributed member of the UBX-like gene family that comprises Ubx3p of *S. cerevisiae* and PUX10 of *A. thaliana*, which are cofactors of Cdc48 and are respectively involved in clathrin-dependent endocytosis and in the dislocation of oleosins from lipid droplets (Farrell et al. 2015; Deruyffelaere et al. 2018). We reconstructed a phylogenetic tree of those Ubx-like proteins (supplementary fig. S13) and observed that s_Ubx proteins are only found in Stramenopiles, forming a monophyletic group nested within Rhodophyta, suggesting a transmission by secondary EGT. If this protein is indeed involved in SELMA, it would be specific to Stramenopiles and may not be an ancestral member of the transporter that evolved initially in the secondary red plastid.

**FIG 4.**
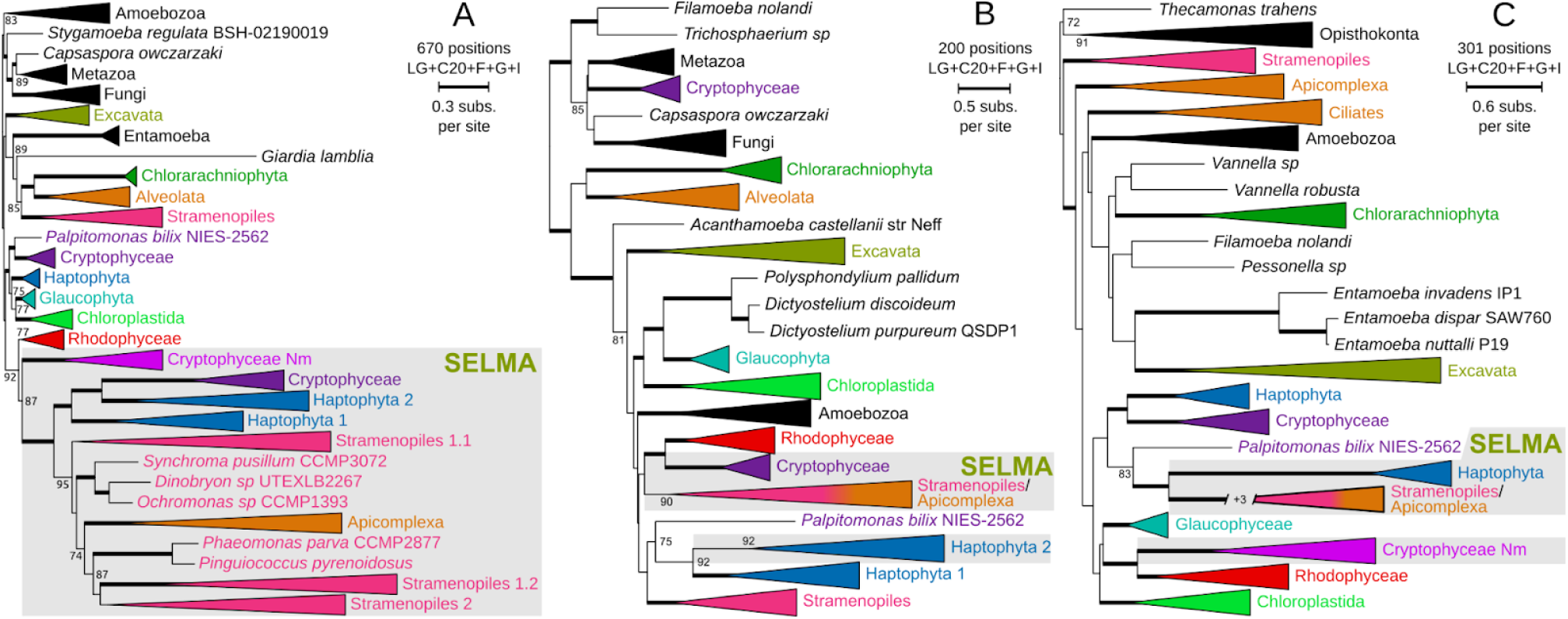
Unrooted condensed maximum likelihood phylogenetic trees of Cdc48 *(A)*, Npl4 *(B)* and Ufd1 (*C*) proteins. The corresponding expanded trees are available in supplementary figs. S9, S10 and S11, respectively. Monophyletic phyla are depicted as triangles, ultrafast bootstrap branch support values (1000 replicates) are indicated except when lower than 70% or as a thick branch when the value is maximum. The number of amino acid positions in the corresponding sequence alignment, the substitution model and parameters used for IQ-TREE reconstruction as well as the branch length scale are indicated at the top right of each tree. Clades corresponding to proteins putatively implicated in SELMA are shadowed in gray.

### Accessory proteins

ERAD is dedicated to the retrotranslocation of misfolded ER proteins. When they reach the cytosol, these proteins are tagged with ubiquitin and sent to the proteasome for degradation. Png1 and its interacting partner Rad23 are suspected to play an important role in the functional link between ERAD and the proteasome (Suzuki et al. 2001). Png1 is the dominant deglycosylating enzyme of the eukaryotic cell and is active in the cytoplasm and the nucleus. Efficient degradation of glycosylated proteins requires a trimming of their glycosidic moieties (Kim et al. 2006). Rad23 was first described as a lesion recognition factor involved in DNA repair. With its ubiquitin-interacting domains, Rad23 also plays the role of cargo for ubiquitinated proteins toward the proteasome (Dantuma et al. 2009). SELMA imported pre-proteins are not supposed to be degraded, and the need to immediately drive them to a putative PPS-located proteasome appears irrelevant. However, trimming pre-protein that might have been accidentally glycosylated in the ER could be required for their proper functioning in the plastid. We gathered orthologs of Png1 and reconstructed a phylogenetic tree of those proteins (supplementary fig. S14). This tree is highly unresolved, with no phylum being properly assembled. We could, however, observe an outlier clade composed of proteins encoded exclusively in CASH genomes, with some being putatively addressed to the plastid. The origin of this paralog, which is likely linked to secondary endosymbioses and could be an accessory protein of SELMA, is unfortunately non-traceable. We also searched for Rad23 homologs in our database. Contrary to Png1, the phylogenetic tree of Rad23 properly assembles all major eukaryotic phyla, but does not contain a derived clade that resembles those we observed for other SELMA genes that evolved conjointly in all CASH (supplementary fig. S15). Only Cryptophyta and Haptophyta have a second copy of Rad23, which was inherited from Rhodophyta in the case of Cryptophyta but was duplicated from the canonical ortholog in Haptophyta. However, none of those proteins are predicted to be targeted to the plastid, which leaves interrogations about their role in the cell let alone in SELMA.

The protein chaperone Hsp70 of the Ssa family, together with cochaperone Hsp40 (Ydj1 in *Saccharomyces*), is a post-ERAD factor that also participates in the degradation of cytoplasmic misfolded proteins. It recognizes misfolded protein domains and ensures their solubility while they are being delivered to the proteasome (Park et al. 2007; Lee et al. 2016). A PPS located Hsp70 protein was first described in studies deciphering the structure of the BTS in Cryptophyta and diatoms (S.B. Gould et al. 2006) and was proposed to be a cofactor of SELMA but with no determined function (Stork et al. 2012). Our phylogenetic tree of Hsp70 shows that all photosynthetic CASH except Alveolata have a second copy of Hsp70 (supplementary fig. S16). In Cryptophyta, s_Hsp70 is encoded in the nucleomorph and is related to e_Hsp70 of Rhodophyta. Alternatively, Haptophyta and Stramenopiles share the same s_Hsp70 paralog that appears unrelated to the nucleomorph encoded protein and whose position in the tree does not allow to infer its origin. Finally, using a systematic screening procedure in *Toxoplasma*, Sheiner et al. (2011) have identified a gene encoding a PPS-located protein that, when disrupted, leads to the blockage of protein import into the plastid and the death of the parasite. This protein, named PPP1 is related to the Sey1 protein of *Saccharomyces* that is involved in homotypic ER vesicles fusion (Anwar et al. 2012). Cryptophyta nucleomorphs encode three to four *sey1*-like copies (Zauner et al. 2019), each being related to a specific paralog of Rhodophyta, and three of them (including PP1) being also present and probably inherited by EGT in CASH genomes (supplementary figures 17a, b and c). This observation suggests that those proteins probably have an important role in the functioning of the red plastid, and because all paralogs already existed in Rhodophyta before secondary endosymbiosis, they may not be directly involved in SELMA.

## DISCUSSION

The route taken by proteins to travel across the periplastidial membrane of complex plastids has long remained a mystery. The discovery of the SELMA transporter was a smart intuition, inspired by the detection of members of an ERAD-like complex encoded in the nucleomorph of Cryptophyta. Much experimental evidence has now proven that a multi-protein complex located in the PPM of all CASH species containing a red-alga derived plastid is responsible for the translocation of pre-proteins from the CER to the PPS. The requirement of pre-protein ubiquitination for their effective translocation is also demonstrated and highlights the existence of ubiquitin-using pathways in secondary plastids that may extend beyond protein import. The composition of SELMA across CASH species and the origin of its components have been the subject of previous studies and are explained by endosymbiotic gene transfers and refunctionalization of ERAD genes from secondary red algal endosymbionts. The complexity of SELMA, its intricate evolution from ERAD and its similarity in all CASH has prompted authors to argue, on the principle of parsimony, that it must have had a single evolutionary origin (Zimorski et al. 2014; Gould et al. 2015; Cavalier-Smith 2018). This single origin conjecture has, however, not been thoroughly and systematically evaluated using phylogenetics. In this work, we reevaluated the origin of 21 of the demonstrated or putative components of the SELMA complex. This analysis offers, for the first time, a complete image of the molecular history of this complex and provides information about the evolution of red algae-derived plastids.

### The composition of the SELMA complex is highly conserved

Fig. 1c summarizes the presence/absence of each SELMA component in CASH genomes, the inferred origin of those components in each lineage as well as how they relate to one another. As previously reported by Stork et al (2012), we observe that the level of conservation of the SELMA complex is remarkable, both in composition and origins, with 13 proteins being present in at least 3 CASH lineages, and a majority being of red algal origin directly or indirectly. First, the two Derlins are always present, even in the more reduced genomes of Apicomplexa parasites, underlining a complementary role of each subunit in the structure or functioning of the pore complex. Secondly, within the inferred PPS-localized ubiquitination machinery, PUBL, s_Uba1 and s_Ubc are conserved, and we could show that the E2 enzyme s_Ubc evolved from the same family of Ubc-like genes in all CASH (fig. 3b). The identity of the E3 enzyme is however still elusive. Considering the conservation pattern that we generally observe, but also the functional specificity of the E2/E3 couple in ubiquitin-using pathways, it seems reasonable to imagine that the ancestral SELMA E3 enzyme is neither s_Hrd1 nor ptE3p. Several publications have evidenced an endomembrane network in the PPS of Cryptophyta as well as a proteasome (Stork et al. 2012; Cavalier-Smith 2018). We could also determine that Cryptophyta have kept many Ubc homologs encoded in the nucleomorph and involved in protein degradation pathways. In *S. cerevisiae*, Hrd1 is able to single-handedly export misfolded proteins when overexpressed *in vivo* or in reconstituted vesicle experiments (Wu and Rapoport 2018). In Cryptophyta, s_Hrd1 might be involved in a PPS localized protein degradation pathway, and may even be sufficient to process a remnant ERAD-like process. On the other hand, ptE3p only exists in Stramenopiles where it likely evolved from the duplication of another E3 enzyme. Thus, there might be an alternative SELMA E3 enzyme that we tried to identify. We referred to the literature to determine which E3 enzymes interact with ATG7-type Ubc in *A. thaliana* and *Homo sapiens* (Kraft et al. 2005; Markson et al. 2009). Unfortunately, none of the known interactors had homologs showing a phylogenetic topology similar to other SELMA components. Besides, the putative deubiquitination of pre-proteins after their translocation is supported by the presence and common origin of ptDUP in Haptophyta, Stramenopiles and Alveolata. Its absence in Cryptophyta is surprising and could be due to the existence of an alternative enzyme for this function; an enzyme that might be shared with other processes utilizing Ubiquitin in the plastid that did not persist in other CASH. Concerning the Cdc48 complex, its four main components are conserved in CASH (Cdc48-1, Cdc48-2, Ufd1 and Npl4) and, similarly to Derlins, the preservation of both Cdc48 isoforms (except for Alveolata that lack isoform 2) indicates a possible specialization as experimentally observed by Lau et al. (2016). The two potential additional components of the Cdc48 complex detected by interaction assays (s_PUB and s_Ubx) are restricted to Stramenopiles and Alveolata. If these two proteins have a role in SELMA, they were not part of the original set that derived from secondary endosymbiosis and probably evolved a posteriori. Finally, three of the four putative accessory proteins that we analyzed are conserved in at least 3 CASH lineages. The enzyme s_Png1 is conserved in all photosynthetic CASH, including Chromerida but not Apicomplexa, in which glycosylation is altered (Samuelson and Robbins 2015). s_Png1 has a unique but unresolved origin in CASH, indicating that deglycosylation of pre-proteins after their translocation is likely an important step. Similarly, the Hsp70 chaperone is conserved but only absent in Alveolata, and PPP1 is of red algal origin in all CASH. However, the exact function of both proteins in SELMA remains undetermined. Altogether, the composition of the SELMA complex appears very stable for as far as we can trace its composition using comparative genomics.

### The history of SELMA genes in relation to Cryptophyta is likely masked

Cryptophyta possess 14 of the 21 analyzed SELMA components: six of them are encoded by nucleomorph genes and seven by nuclear genes. The case of the 14th component, the plastid specific ubiquitin PUBL, is peculiar, because the corresponding gene is found in the nucleomorph of some species and in the nucleus of others, illustrating that the gene content of the nucleomorph is not uniform across Cryptophyta (Moore 2013). Out of the 6 nucleomorph encoded SELMA proteins, the origin of five could be traced back to Rhodophyta, but the phylogeny of only two (s_Der1 and s_Cdc48-1) is compatible with an endosymbiotic transmission of the corresponding nucleomorph genes to the other CASH genomes. Similarly, only three of the nucleus encoded SELMA components of Cryptophyta are closely related to their homologs in CASH species (s_Npl4, s_Png1 and Cdc48-2), and only two (s_Npl4 and s_Rad23) are related to red algae (figure 1c). Altogether, the phylogenies of only five proteins support the possibility of a serial transmission of SELMA from Cryptophyta to other CASH lineages. We have, however, several reasons to consider that some proteins produce misleading phylogenetic trees. First, we observe that 3 of the nuclear-encoded SELMA genes (*s_dfm1*, *s_uba1* and *s_ubc*) were likely replaced by duplication and refunctionalization of homologous nuclear-encoded ERAD genes. The corresponding trees show that Haptophyta, Stramenopiles and Alveolata seem to have inherited those 3 genes by EGT from Rhodophyta (supplementary figs. S3, S5 and S6). We can speculate that the original red-alga derived genes did exist in Cryptophyta but became invisible due to their subsequent replacement by nuclear homologs. Consequently, the true serial EGT origin of those genes in CASH genomes is masked. Additionally, even if the phylogenetic tree of PUBL is poorly resolved (supplementary fig. S4), we observe that the closest, yet weakly supported, homolog to PUBL of Alveolata+Stramenopiles+Haptophyta is the nucleomorph-encoded Ub gene of *C. mesostigmatica,* adding another candidate accounting for the serial endosymbiosis scheme. Finally, in the phylogeny of both s_Hsp70 and s_Ufd1, Haptophyta, Stramenopiles (and Alveolata for s_Ufd1) are monophyletic and branch close to the nuclear ERAD paralog of Cryptophyta (supplementary figs. S16 and S11). If those trees are correct, we might speculate that, during a putative process of tertiary endosymbiosis involving a Cryptophyta ancestor, the host recruited and refunctionalized the ERAD homologs by EGT instead of the nucleomorph-encoded SELMA ones. The fact that four other components of SELMA (s_Dfm1, s_Uba1, s_Ubc and s_Npl4) were actually replaced by ERAD homologs in Cryptophyta or Haptophyta outlines the versatility of such substitutions. Altogether, we conclude that there might be more evidence than previously thought indicating that ancestors of Cryptophyta were the donors of the SELMA complex, and consequently of the red plastid, during tertiary endosymbiosis events.

### Plastids of Stramenopiles, Haptophyta and Alveolata have entangled origins

We reported above that Stramenopiles, Haptophyta and Alveolata have acquired components of the SELMA transporter from mostly three genomic sources: Cryptophyta nucleomorph SELMA genes (*s_der1*, *s_cdc48-1, ub*), Cryptophyta nuclear SELMA genes (e.g. s_npl4, s_png1, s_cdc48-2) or Cryptophyta nuclear ERAD genes (*s_ufd1*, *s_hsp70*). Additionally, out of the 16 SELMA elements successfully analyzed in our work, 11 are shared by Stramenopiles, Haptophyta and Alveolata, and for 10 of those, phylogenetic trees show that these phyla have acquired the same ancestral gene. This strongly argues for the unlikeliness of multiple secondary endosymbiosis involving different red algae, as was proposed recently by Pietluch et al. (Pietluch et al. 2024), but rather for a single origin of the secondary red plastid which likely arose in an ancestor of Cryptophyta. Interestingly, none of the ERAD-to-SELMA gene replacements that we observed in Cryptophyta were transmitted to any other CASH lineage. This reduces the time window of tertiary endosymbiosis involving a Cryptophyta symbiont to a moment predating those replacements, which is before the diversification of their known extant representatives. We also recover the nested relationship of Myzozoa (Alveolata) into Stramenopiles for 12 SELMA proteins which, together with the strikingly unique shared domain composition of s_PUB, strongly argues that the apicoplast derived from the plastid of Stramenopiles, as previously hypothesized (Ševčíková et al. 2015; Pietluch et al. 2024). The solid evolutionary connection between Stramenopiles and Myzozoa discards the model of plastid evolution published by Bodył et al. where the apicoplast of Myzozoa would have derived from an ancestral plastid of Haptophyta (Bodył et al. 2009). This leaves only three possibilities for the number and order of transmissions of the red alga derived plastid. (A) Plastids of ancestral Cryptophyta were acquired separately by the ancestors of Haptophyta and Ochrophyta (Stramenopiles); (B) a single tertiary endosymbiosis occurred between an ancestral Cryptophyta and the ancestor of Ochrophyta who then transmitted their plastid independently to the Haptophyta and Myzozoa (Alveolata) (as proposed by Stiller et al. (2014)); (C) Haptophyta emerged first from a tertiary endosymbiosis involving Cryptophyta, transmitted their plastid to ancestors of Stramenopiles by quaternary endosymbiosis, who later passed on to Myzozoa. Figure 5 presents schematic phylogenetic trees that illustrate those evolutionary hypotheses; upon which the hypothetical dynamic of SELMA genes between and along lineages is displayed. Note that EGT origins (represented in dotted boxes by the color of the donor lineage) are not in accordance with what is presented in figure 1c. Indeed, the latter exposes what can be inferred from our trees while figure 5 incorporates the logic of inter-lineages transfers to further determine gene donors. The fact that the exact same collection of SELMA/ERAD genes was inherited by Stramenopiles, Haptophyta and Alveolata (especially the recruitment of e_ufd1 and e_hsp70 in place of the nucleomorph isoforms), would argue in favor of a single tertiary endosymbiosis, eliminating scenario (A) depicted in figure 5a. The existence of a monophyletic ptDUP in all CASH except Cryptophyta adds another argument in favor of a single tertiary endosymbiosis. Alternative propositions (B) and (C) are very similar in terms of gene dynamics, with the only notable difference that the emergence of ptDUP occurred either in Ochrophyta (case B) or in Haptophyta (case C). The phylogeny of ptDUP (supplementary fig S8) indicates that the corresponding gene likely derived from the duplication of a nuclear gene in Haptophyta (although with moderate support); giving a little more weight to scenario (C). Altogether, we consider that our data point to the following hypothesis for the evolution of the SELMA transporter, and consequently for the evolution of complex plastids (fig. 5c): an ancestor of Cryptophyta acquired its plastid by secondary endosymbiosis and tinkered the ERAD transporter of the red algal symbiont into SELMA to allow the transport of plastid targeted proteins across de PPM. The secondary plastid was then transmitted to an ancestor of Haptophyta by tertiary endosymbiosis, and a specific set of SELMA and ERAD genes were gathered in the nucleus to recompose a slightly different SELMA transporter with the later addition of ptDUP. The whole set was then transmitted together with the plastid to an ancestor of Ochrophyta through another endosymbiosis. s_PUB was then constructed by domain reshuffling in Stramenopiles, and finally, the plastid of Ochrophyta was acquired by an ancestor of Myzozoa where SELMA was later reduced by the loss of s_cdc48-2, s_Hsp70 (plus s_Png1 in Apicomplexa). Later in Cryptophyta, SELMA experienced several modifications by replacements involving ERAD homologs. The differential transfer of the Ubiquitin gene from the nucleomorph to the nucleus is probably the most recent change in SELMA composition in Cryptophyta. It is interesting to mention that none of those hypotheses can simply explain the peculiar taxonomic distribution of *rpl36* isoforms in the plastid genomes of Haptophyta and Cryptophyta (Rice and Palmer 2006). Indeed, the most parsimonious way to account for this distribution is to accept two separate secondary endosymbioses with different red algae: one possessing the canonical *rpl36-p* (at the origin of Stramenopiles) and the other one having acquired *rpl36-c* by horizontal gene transfer before being engulfed by the ancestor of Cryptophyta and Haptophyta. Our study of SELMA tends, however, to exclude dual secondary endosymbioses. We propose in Figure 5, an hypothetical set of gain and loss events concerning those *rpl36* genes that is compatible with the history of SELMA and implies the coexistence of both isoforms in the ancestor of the plastid of Cryptophyta followed by differential losses.

**FIG 5.**
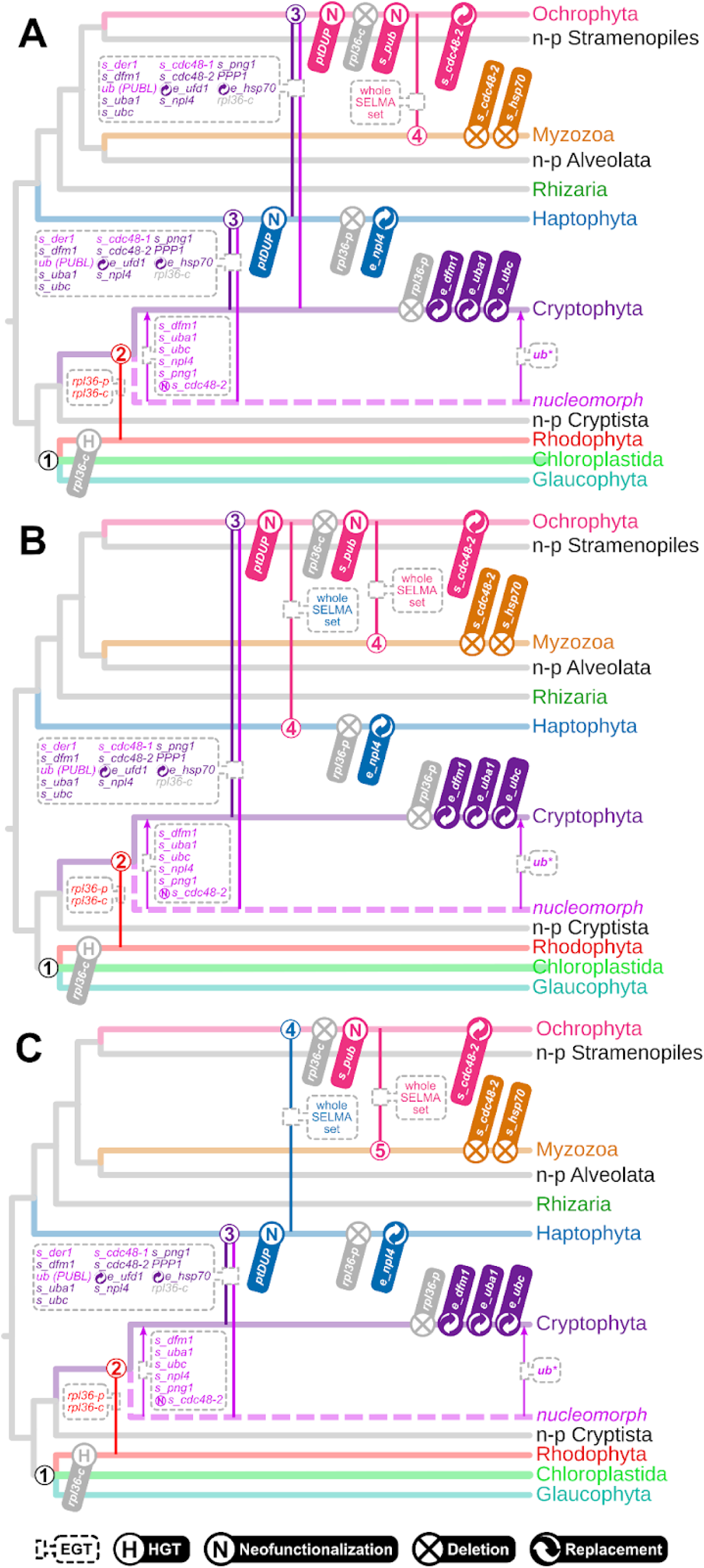
Hypothetical scenarios of the evolution of complex plastids and of the SELMA complex. Panels (*A)*, *(B)* and (*C)* present schematic phylogenetic trees, each displaying an alternative scenario for the evolution of photosynthetic CASH lineages in the form of sequences of endosymbioses. Endosymbiosis events are depicted by vertical lines colored depending on the symbiont lineage and labeled with a number describing the level of endosymbiosis. Genes transferred by EGT are indicated inside dotted boxes and colored depending on the donor lineage. Other types of events affecting genes involved in SELMA such as Horizontal Gene Transfers (HGT), neofunctionalizations, deletions and replacements are indicated by specific signs specified at the bottom of the figure. *EGT of the Ubiquitin coding gene from the nucleomorph to the nucleus only occurred in some species. Hypothetical movements of *rpl36* isoforms are indicated in gray.

## Conclusion

The Chromalveolate hypothesis, proposed by Cavalier-smith in 1997, argued that all photosynthetic lineages having plastids of red-algal origin emerged from a single secondary endosymbiosis. One of the prevailing arguments of that model was that fully integrated endosymbioses must have been very rare, because they are very complex evolutionary events that require tremendous modifications of the cell structure as well as important reshufflings of genomes and metabolic networks. In the meantime, as our understanding of microbial diversity improved, it has become clear that symbioses, and even intracellular symbioses, are very common, and that many of them have remarkably shaped the course of evolution (López-García et al. 2017). Although presented 20 years ago, the first proposals challenging the Chromalveolate hypothesis have long been neglected before being brought back in the spotlight by the accumulation of molecular data falsifying the monophyly of Chromalveolates (Bodył 2005; Sanchez-Puerta et al. 2007; Bodył et al. 2009; Baurain et al. 2010; Petersen et al. 2014; Stiller et al. 2014; Strassert et al. 2020). The idea that each photosynthetic phylum of this group could have been the result of tertiary, quaternary or even more complex serial endosymbioses, passing the same red plastid along the way, is now considered a more plausible scenario. Among the many innovations that are required to transform symbionts into fully integrated organelles, the ability to address proteins encoded by nuclear genes back into the symbiont-derived compartment is crucial. Implementing such mechanisms is complex enough to consider that non-related endosymbiotic events are very unlikely to develop the same solution by chance. Thus, the observation that all Archaeplastida use the same Tic/Toc transporter to translocate proteins into their plastids has been determinant in establishing the monophyly of primary plastids (Steiner et al. 2005). In this study, we provide new evidence that the SELMA transporter, which is responsible for the transport of proteins across the second outermost membrane of complex red plastids, has very likely a single origin, taking place after the secondary endosymbiosis between a red alga and an ancestor of Cryptophyta. We also show that SELMA is present in all other photosynthetic CASH, and that many of its components are homologous to those found in Cryptophyta. Moreover, we pinpoint the relative timing of subtle modifications of the composition of the complex and the horizontal transmission of those innovations between lineages. The evolutionary scenario that we infer from our results supports the idea of serial endosymbioses and of the transfer of the red plastid from Cryptophyta to Haptophyta, from Haptophyta to Stramenopiles, and from Stramenopiles to Alveolata (fig. 5c). This chain of events slightly differs from what was proposed by Bodył et al. (2009) and Stiller et al. (2014) (fig. 5b), but is not contradictory with the estimated dating of endosymbioses inferred by Strassert et al. (2020). Our proposal is based on the interpretation of a collection of single gene phylogenies, which might be seen as moderately resolutive considering the evolutionary timescale of plastid endosymbioses. These interpretations are, however, constrained and reinforced by the need for conservation of elements that must interact properly to secure the very important process of protein targeting. This new glance at the evolution of complex plastids would need to be challenged using wider phylogenomic data.

## MATERIAL AND METHODS

### Dataset and phylogenetic tree reconstruction

A custom SQL database was used to store and organize a set of about 5,5 million protein sequences predicted from 864 genomes and transcriptomes of species representing the diversity of the three domains of life, with an emphasis on photosynthetic eukaryotes. The complete list of assemblies that were used in this database is available in supplementary table S3. The starting material of all analyses in the present work is the set of accession numbers reported in supplementary table S1 in Stork et al. (2012). Protein sequences corresponding to these accession numbers were retrieved and used to query our local custom database using reciprocal BLASTP (Altschul et al. 1990), allowing for 5 rounds of sequence searches with an e-value threshold of 1e-5. All hits for each reference protein were collected in a multiple sequence file and used to produce phylogenetic trees as follows: (1) sequences were aligned using mafft v7.205 with default parameters (Katoh and Toh 2010); (2) a custom python script was used to trim the alignments using relaxed conditions: deletion of columns showing more than 20% of gaps and pruning of sequences having a length coverage lower than 10% of the trimmed MSA. A first set of phylogenetic trees were reconstructed using FastTree v2.1.7 (LG + Gamma + CAT20) (Price et al. 2010). These first trees were manually inspected and pruned to remove duplicate sequences as well as sub-groups of similar proteins that were not directly related to the homologous group of interest. The remaining sequences were re-extracted from the database, realigned and trimmed using the same previous conditions, and the manual inspection and elimination of distant sequences was repeated. The final sets of homologous protein sequences that we selected are available as supplementary material. For the final tree reconstructions, these sets were realigned, and subjected to different strategies of trimming (using trimAl (Capella-Gutiérrez et al. 2009)) depending on the size of each analyzed SELMA component in order to keep enough informative positions and recover phylogenies where major eukaryotic phyla were retrieved as solid clades with moderate to high bootstrap support values. Supplementary table 2 indicates the chosen strategy for each protein set. Final trees were reconstructed using IQ-TREE stable version 1.6.12, a combination of mixture model parameters detailed in supplementary table 2, as well as 1000 ultrafast bootstrap replicates (Minh et al. 2013; Nguyen et al. 2014). The only variation between models used for each alignment was the number of estimated classes in mixture models (C10/C20/C40). We decided to keep the parameter that maximized the support for the root branch of all widely accepted eukaryotic phyla or superphyla. The final trees were edited using FigTree version 1.4 (Rambaut 2009) and the vector graphics program Inkscape (https://inkscape.org/).

### Prediction of proteins sub-cellular localization

All non-aligned, untrimmed protein sequences used for producing final phylogenetic trees were submitted online to HECTAR˄SEC - https://webtools.sb-roscoff.fr/root?tool_id=abims_hectar - (Gschloessl et al. 2008) and locally to ASAFind (Gruber et al. 2015) to determine their putative sub-cellular localization, either toward the secretion pathway or to the secondary plastid. Many protein sequences proved to be truncated in their N-terminal portion, in particular for those generated from transcriptome data, and were therefore ignored by those detection programs.

### Conserved domain searches and Alphafold data

Conserved domains of ptDUP and related proteins were determined using the search engine of the Conserved Domain Database (Wang et al. 2023) hosted on the NCBI website (https://www.ncbi.nlm.nih.gov/Structure/cdd/wrpsb.cgi). The coordinates of the detected domains were exported to manually create the maps presented in supplementary figure S12 using the vector graphics program Inkscape (https://inkscape.org/). Alphafold 3D model representations of PUBL proteins of *Toxoplasma gondii* (TGME49_223125, Uniprot S8F891) and *Plasmodium falciparum* (PF3D7_0815700, Uniprot C0H4U7) presented in supplementary figure S4 were retrieved from the AlphaFold Protein Structure Database (Varadi et al. 2022). Alignment of those same 3D models was achieved using the online tool ICN3D (Wang et al. 2022).

## SUPPLEMENTARY INFORMATION AND DATA AVAILABILITY

Supplementary figures S1 to S17 and supplementary tables S1 to S3 are available on the publisher website. The same supplementary files, as well as multiple protein sequence alignments in fasta format and phylogenetic trees in Newick format are also available at https://data.deemteam.fr/

## AUTHOR CONTRIBUTIONS

P.D. and D.M. designed the study. R.P.T. conducted all phylogenetic experiments under the supervision of P.D. & D.M. All authors contributed to the writing and review of this manuscript. D.M. and P.L.-G. were supported by grants from the European Research Council (ERC Advanced grants 787904 and 101141745, respectively).

