## Supplementary_material for "Molecular phylogeny of the SELMA translocation machinery recounts the evolution of complex photosynthetic eukaryotes"

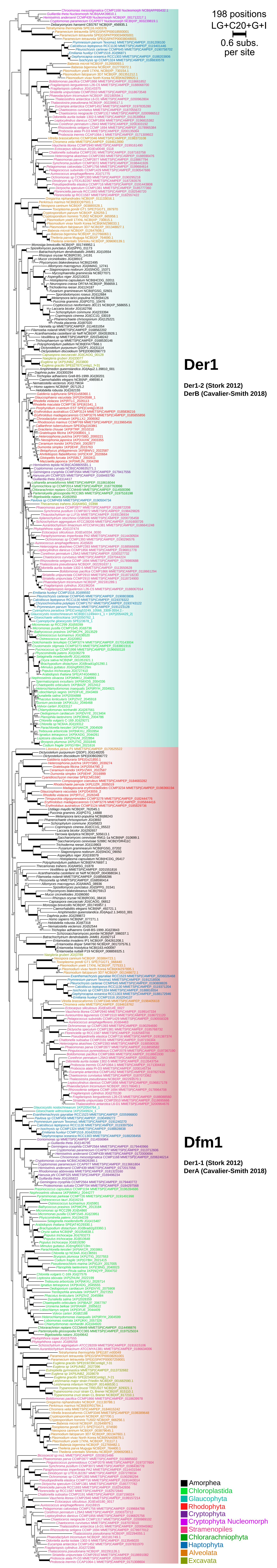

**Supplementary fig. S1. Maximum likelihood consensus phylogenetic tree of Derlin proteins.**
The tree is artificially rooted beetwen Der1 and Dfm1 paralog clades. Ultrafast bootstrap branch support values (1000 replicates) are indicated except when lower than 70% or as a thick line when value is maximum. The number of amino acid positions in the corresponding sequence alignment, the substitution model and parameters used for IQ-TREE reconstruction as well as the branch length scale are indicated at the top right. Phyla color legend is provided at the bottom right. Protein identifiers indicate the source and the reference separated by a pipe symbol. Der1 paralog is highlighted in blue; Dfm1 in red.

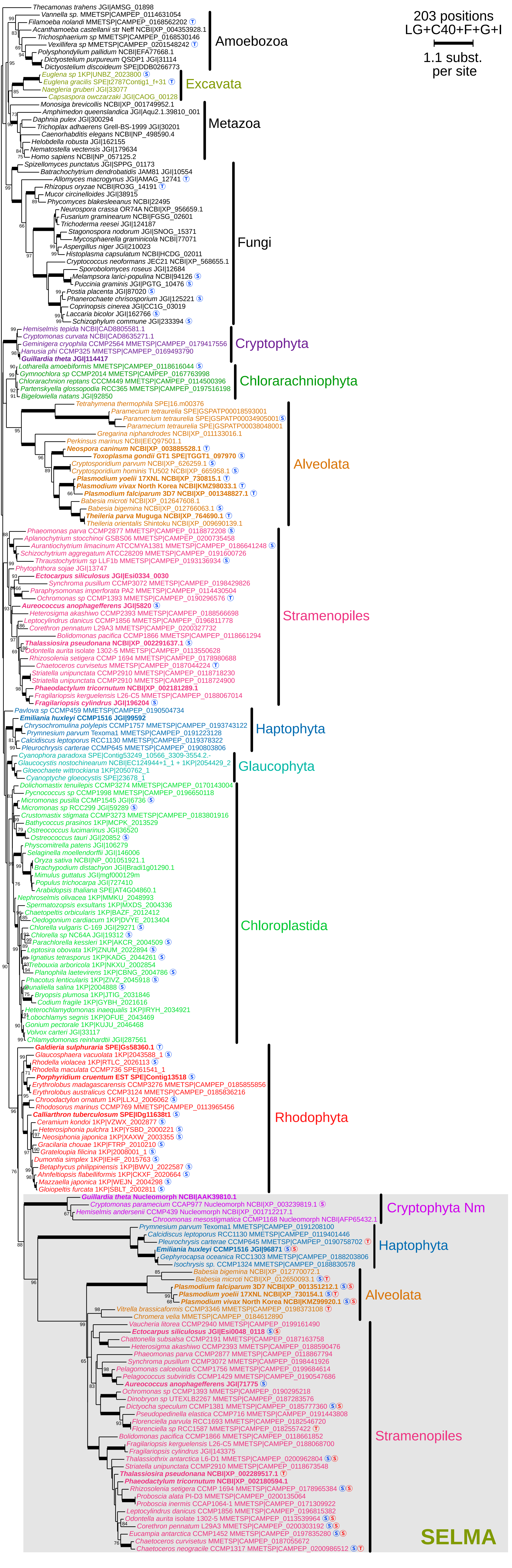

**Supplementary fig. S2. Unrooted maximum likelihood consensus phylogenetic tree of Der1 proteins.** Ultrafast bootstrap branch support values (1000 replicates) are indicated except when lower than 70% or as a thick line when value is maximum. The number of amino acid positions in the corresponding sequence alignment, the substitution model and parameters used for IQ-TREE reconstruction as well as the branch length scale are indicated at the top right. Protein identifiers indicate the source and the reference separated by a pipe symbol. Leaves corresponding to proteins reported in Stork et al. 2012 are in bold. For each protein, the predicted cellular localisation is tagged by a S for "Signal peptide" and a T for "bipartite Transit sequence" as predicted by Hectar in blue and ASAfind in red. Clades corresponding to proteins putatively implicated in SELMA are shadowed in gray.

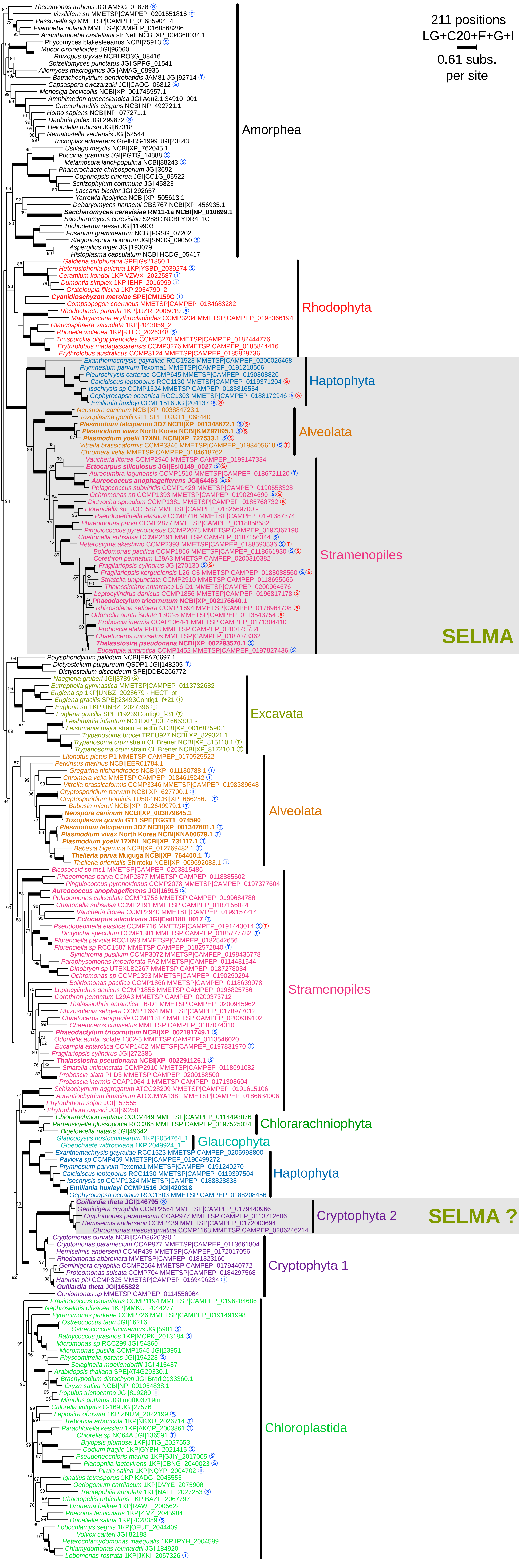

**Supplementary fig. S3. Unrooted maximum likelihood consensus phylogenetic tree of Dfm1 proteins.** Ultrafast bootstrap branch support values (1000 replicates) are indicated except when lower than 70% or as a thick line when value is maximum. The number of amino acid positions in the corresponding sequence alignment, the substitution model and parameters used for IQ-TREE reconstruction as well as the branch length scale are indicated at the top right. Protein identifiers indicate the source and the reference separated by a pipe symbol. Leaves corresponding to proteins reported in Stork et al. 2012 are in bold. For each protein, the predicted cellular localisation is tagged by a S for "Signal peptide" and a T for "bipartite Transit sequence" as predicted by Hectar in blue and ASAfind in red. Clades corresponding to proteins putatively implicated in SELMA are shadowed in gray.

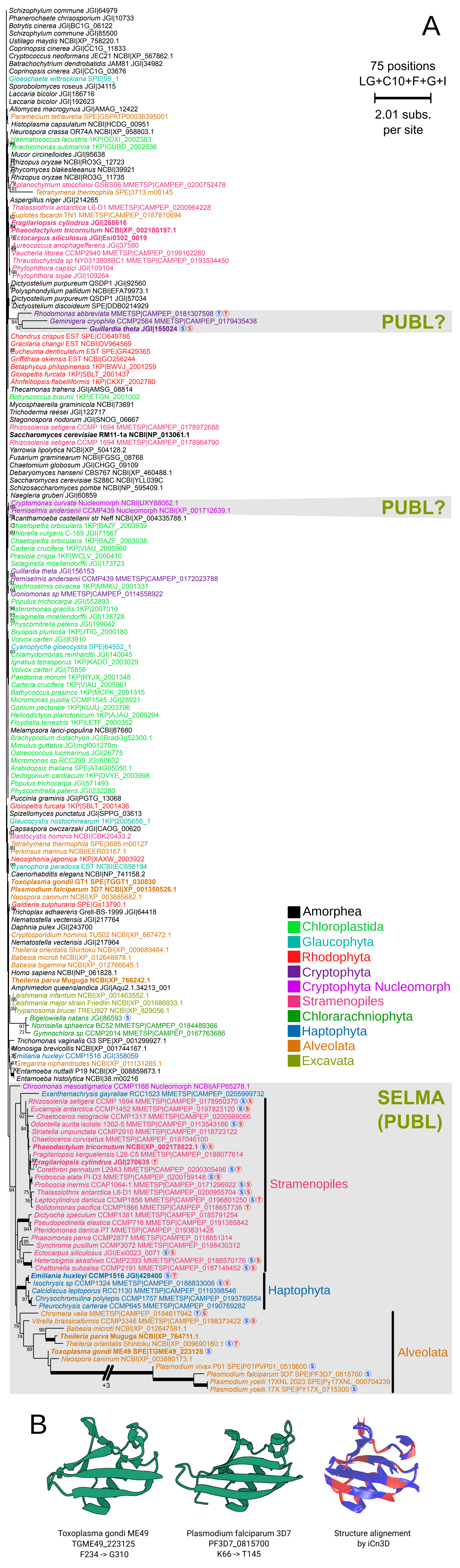

**Supplementary fig. S4. (A) Unrooted maximum likelihood consensus phylogenetic tree of Ubiquitin proteins.** Ultrafast bootstrap branch support values (1000 replicates) are indicated except when lower than 70% or as a thick line when value is maximum. The number of amino acid positions in the corresponding sequence alignment, the substitution model and parameters used for IQ-TREE reconstruction as well as the branch length scale are indicated at the top right. Protein identifiers indicate the source and the reference separated by a pipe symbol. Leaves corresponding to proteins reported in Stork et al. 2012 are in bold. For each protein, the predicted cellular localisation is tagged by a S for "Signal peptide" and a T for "bipartite Transit sequence" as predicted by Hectar in blue and ASAfind in red. Clades corresponding to proteins putatively implicated in SELMA are shadowed in gray. **(B) Comparison of AlphaFold predicted structural models** of the Ubiquitin-like portions of the PUBL protein from Toxoplasma gondii ME49 Plasmodium falciparum 3D7. The model on the right displays the structural alignment of both proteins as computed using ICN3D.

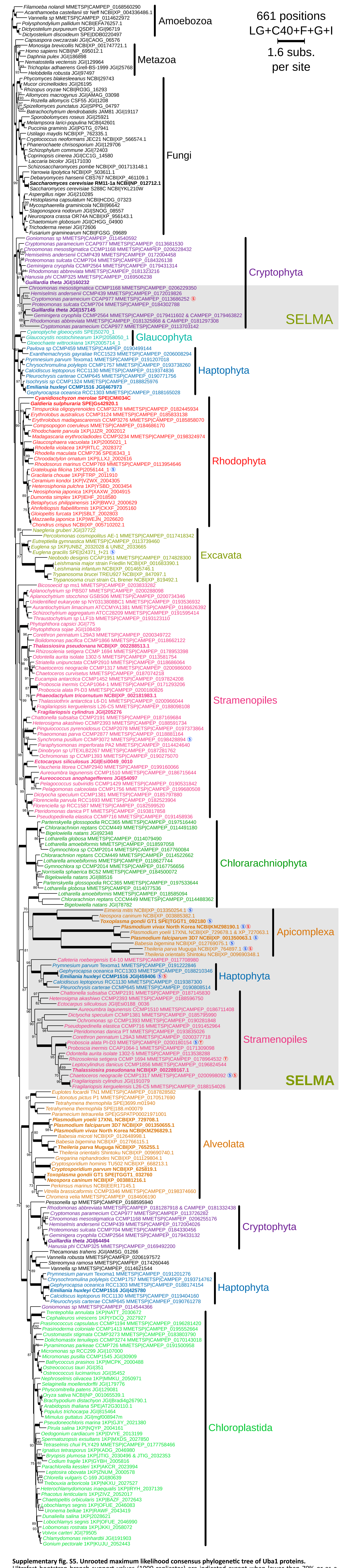

**Supplementary fig. S5. Unrooted maximum likelihood consensus phylogenetic tree of Uba1 proteins.**
Ultrafast bootstrap branch support values (1000 replicates) are indicated except when lower than 70% or as a thick line when value is maximum. The number of amino acid positions in the corresponding sequence alignment, the substitution model and parameters used for IQ-TREE reconstruction as well as the branch length scale are indicated at the top right. Protein identifiers indicate the source and the reference separated by a pipe symbol. Leaves corresponding to proteins reported in Stork et al. 2012 are in bold. For each protein, the predicted cellular localisation is tagged by a S for “Signal peptide” and a T for “bipartite Transit sequence” as predicted by Hectar in blue and ASAfind in red. Clades corresponding to proteins putatively implicated in SELMA are shadowed in gray.

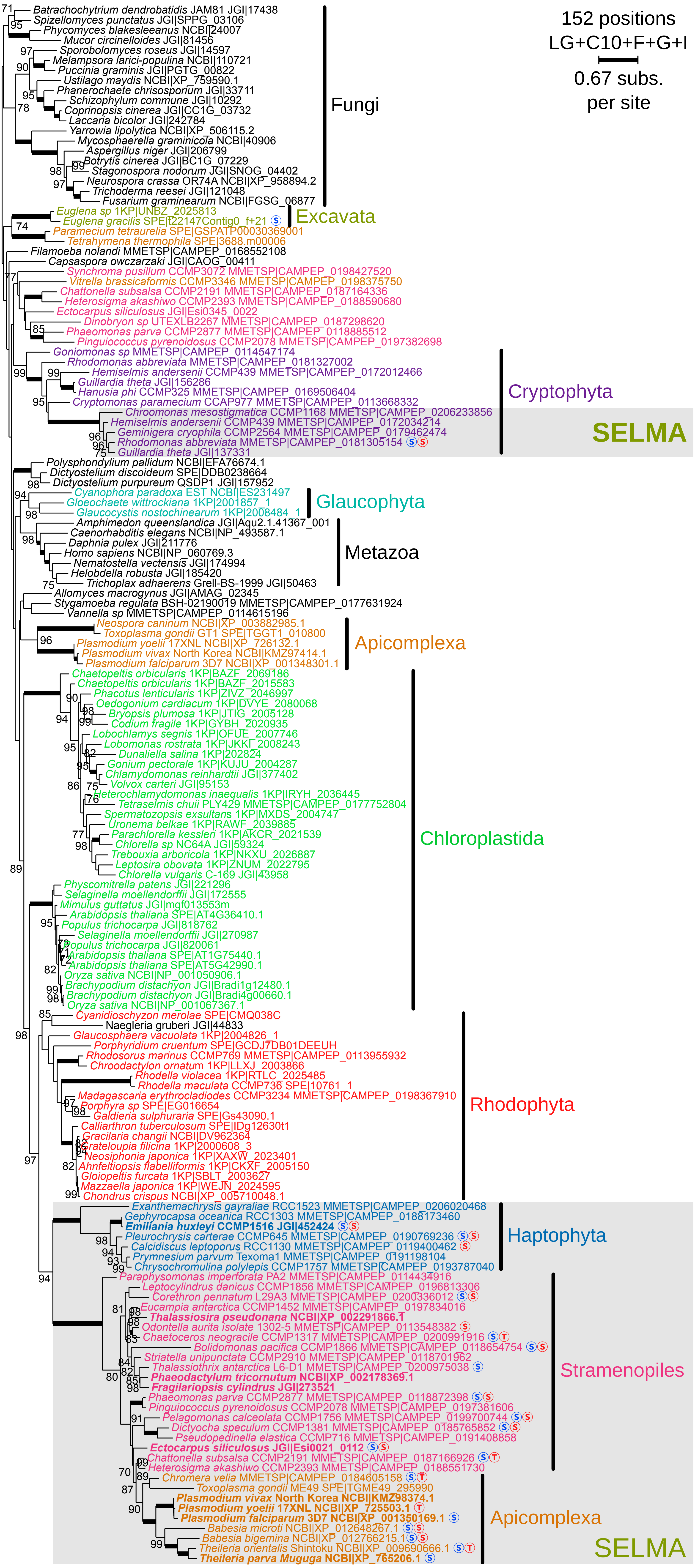

**Supplementary fig. S6. Unrooted maximum likelihood consensus phylogenetic tree of Atg7 type Ubc proteins.** Ultrafast bootstrap branch support values (1000 replicates) are indicated except when lower than 70% or as a thick line when value is maximum. The number of amino acid positions in the corresponding sequence alignment, the substitution model and parameters used for IQ-TREE reconstruction as well as the branch length scale are indicated at the top right. Protein identifiers indicate the source and the reference separated by a pipe symbol. Leaves corresponding to proteins reported in Stork et al. 2012 are in bold. For each protein, the predicted cellular localisation is tagged by a S for "Signal peptide" and a T for "bipartite Transit sequence" as predicted by Hectar in blue and ASAfind in red. Clades corresponding to proteins putatively implicated in SELMA are shadowed in gray.

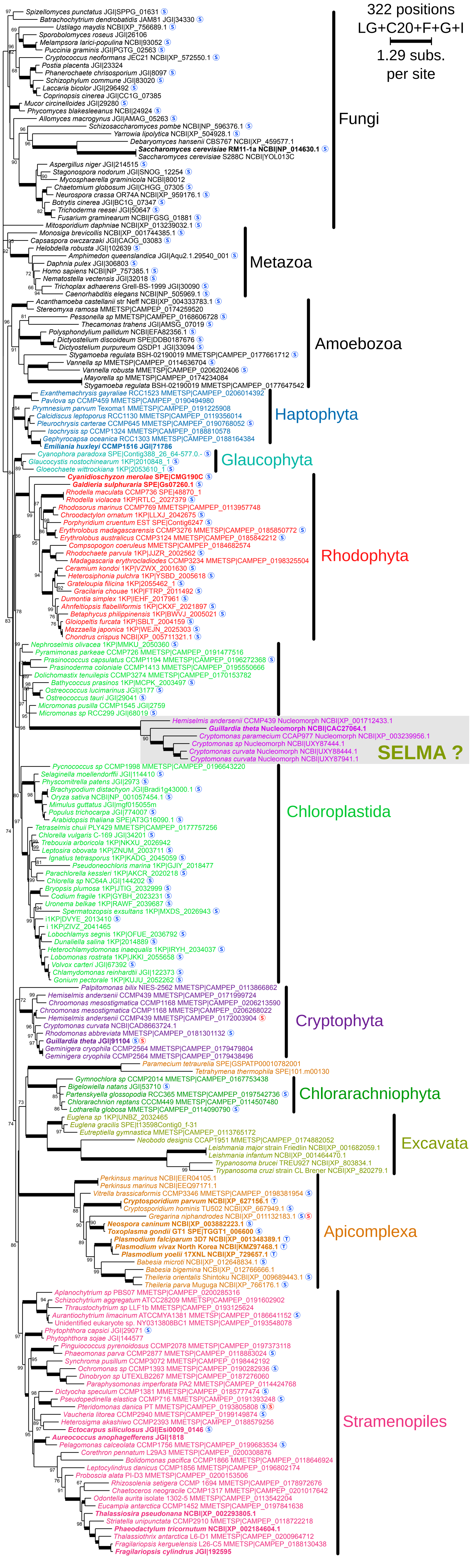

**Supplementary fig. S7. Unrooted maximum likelihood consensus phylogenetic tree of Hrd1 proteins.**
Ultrafast bootstrap branch support values (1000 replicates) are indicated except when lower than 70% or as a thick line when value is maximum. The number of amino acid positions in the corresponding sequence alignment, the substitution model and parameters used for IQ-TREE reconstruction as well as the branch length scale are indicated at the top right. Protein identifiers indicate the source and the reference separated by a pipe symbol. Leaves corresponding to proteins reported in Stork et al. 2012 are in bold. For each protein, the predicted cellular localisation is tagged by a S for "Signal peptide" and a T for "bipartite Transit sequence" as predicted by Hectar in blue and ASAfind in red. Clades corresponding to proteins putatively implicated in SELMA are shadowed in gray.

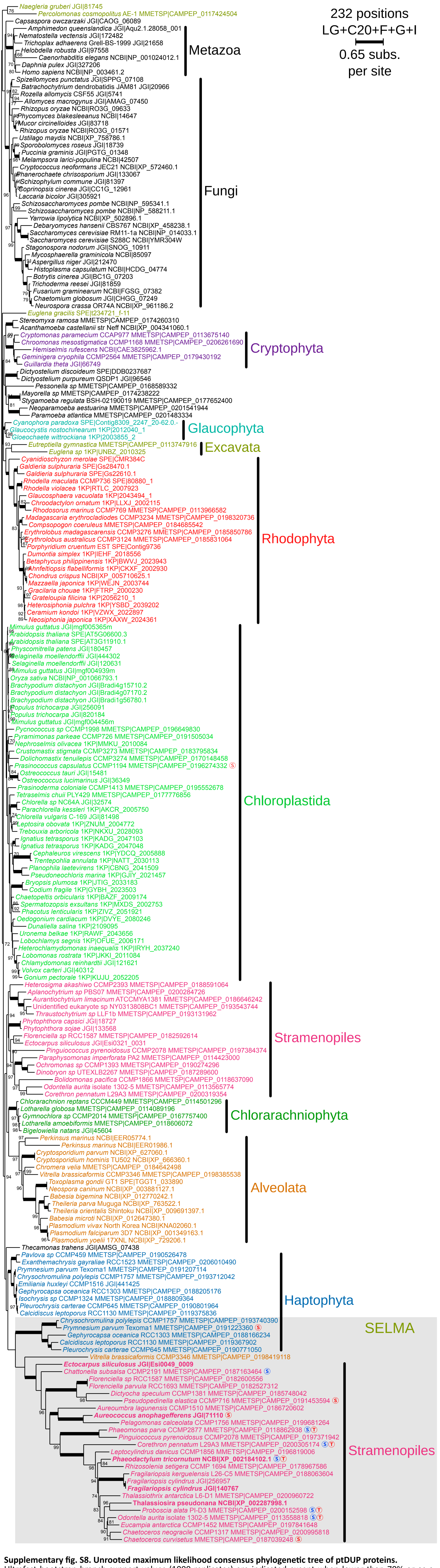

**Supplementary fig. S8. Unrooted maximum likelihood consensus phylogenetic tree of ptDUP proteins.** Ultrafast bootstrap branch support values (1000 replicates) are indicated except when lower than 70% or as a thick line when value is maximum. The number of amino acid positions in the corresponding sequence alignment, the substitution model and parameters used for IQ-TREE reconstruction as well as the branch length scale are indicated at the top right. Protein identifiers indicate the source and the reference separated by a pipe symbol. Leaves corresponding to proteins reported in Stork et al. 2012 are in bold. For each protein, the predicted cellular localisation is tagged by a S for "Signal peptide" and a T for "bipartite Transit sequence" as predicted by Hectar in blue and ASAfind in red. Clades corresponding to proteins putatively implicated in SELMA are shadowed in gray.

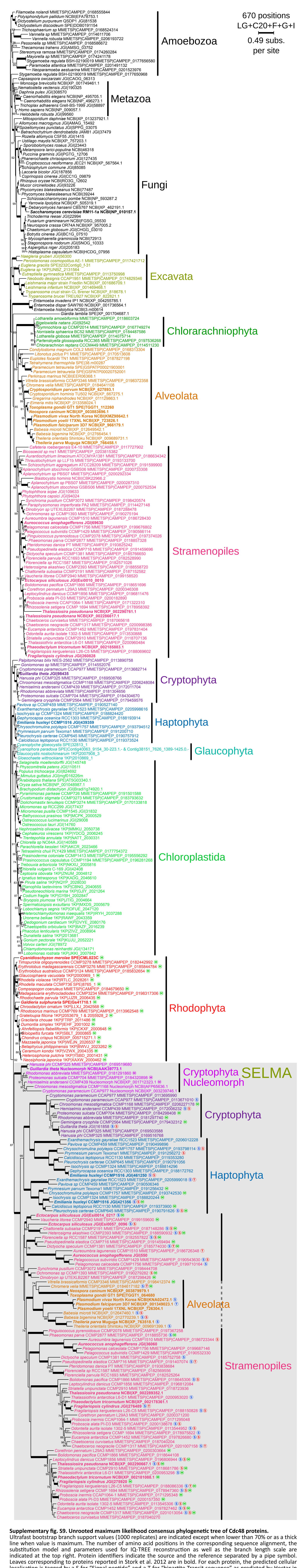

**Supplementary fig. S9. Unrooted maximum likelihood consensus phylogenetic tree of Cdc48 proteins.** Ultrafast bootstrap branch support values (1000 replicates) are indicated except when lower than 70% or as a thick line when value is maximum. The number of amino acid positions in the corresponding sequence alignment, the substitution model and parameters used for IQ-TREE reconstruction as well as the branch length scale are indicated at the top right. Protein identifiers indicate the source and the reference separated by a pipe symbol. Leaves corresponding to proteins reported in Stork et al. 2012 are in bold. For each protein, the predicted cellular localisation is tagged by a S for “Signal peptide” and a T for “bipartite Transit sequence” as predicted by Hectar in blue and ASAfind in red. Additionally, the presence of the <DDDLYS*> motif described by Lau et al. (2016) is tagged by a M letter. Clades corresponding to proteins putatively implicated in SELMA are shadowed in gray.

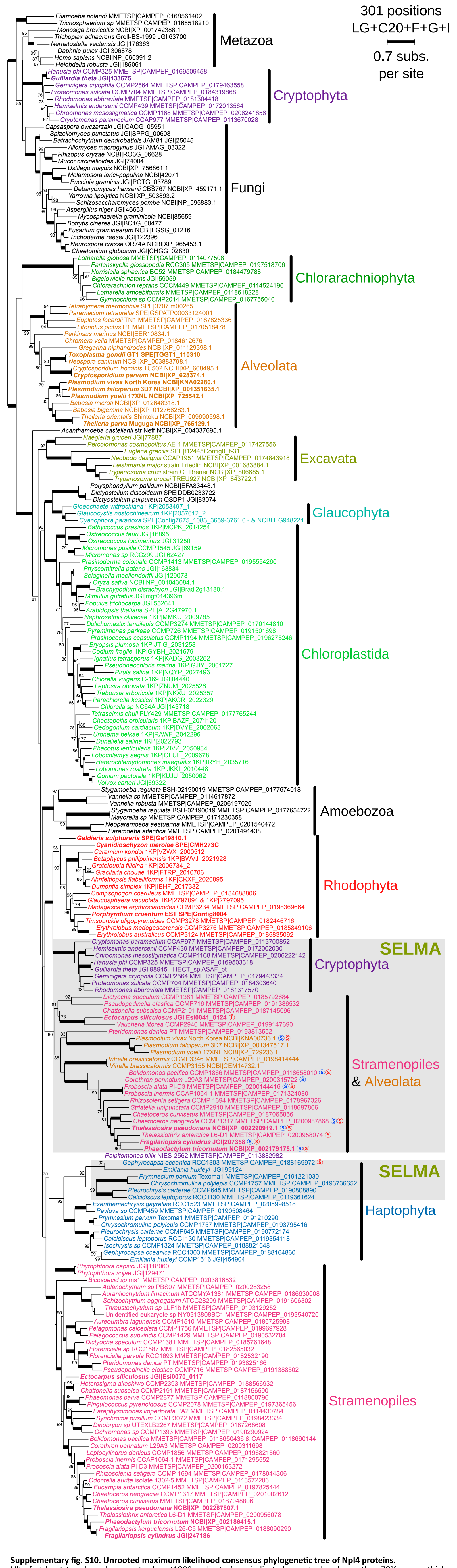

**Supplementary fig. S10. Unrooted maximum likelihood consensus phylogenetic tree of Npl4 proteins.**
Ultrafast bootstrap branch support values (1000 replicates) are indicated except when lower than 70% or as a thick line when value is maximum. The number of amino acid positions in the corresponding sequence alignment, the substitution model and parameters used for IQ-TREE reconstruction as well as the branch length scale are indicated at the top right. Protein identifiers indicate the source and the reference separated by a pipe symbol. Leaves corresponding to proteins reported in Stork et al. 2012 are in bold. For each protein, the predicted cellular localisation is tagged by a S for "Signal peptide" and a T for "bipartite Transit sequence" as predicted by Hectar in blue and ASAfind in red. Clades corresponding to proteins putatively implicated in SELMA are shadowed in gray.

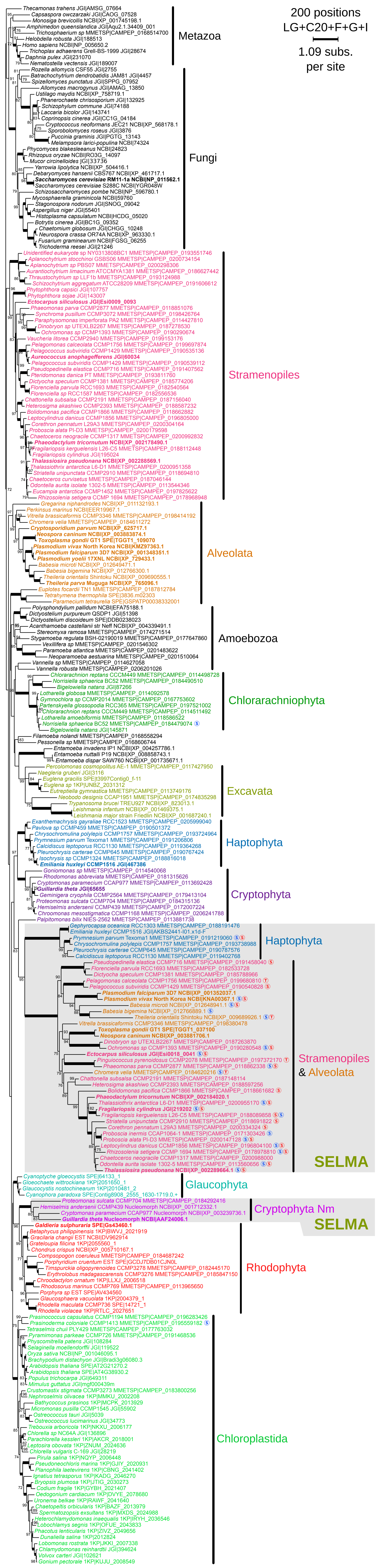

**Supplementary fig. S11. Unrooted maximum likelihood consensus phylogenetic tree of Ufd1 proteins.** Ultrafast bootstrap branch support values (1000 replicates) are indicated except when lower than 70% or as a thick line when value is maximum. The number of amino acid positions in the corresponding sequence alignment, the substitution model and parameters used for IQ-TREE reconstruction as well as the branch length scale are indicated at the top right. Protein identifiers indicate the source and the reference separated by a pipe symbol. Leaves corresponding to proteins reported in Stork et al. 2012 are in bold. For each protein, the predicted cellular localisation is tagged by a S for “Signal peptide” and a T for “bipartite Transit sequence” as predicted by Hectar in blue and ASAfind in red. Clades corresponding to proteins putatively implicated in SELMA are shadowed in gray.

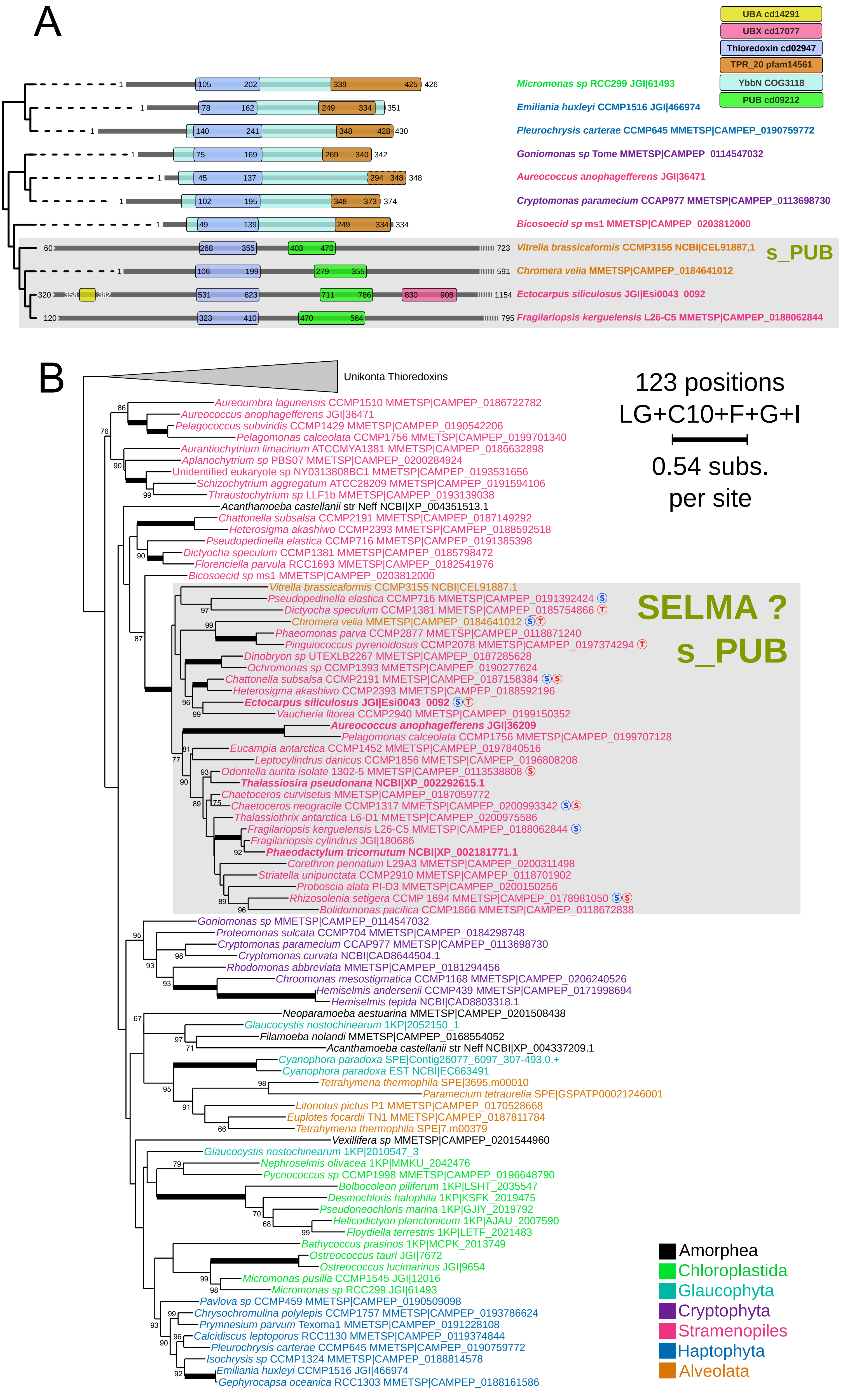

**Supplementary fig. S12. (A) Conserved domains composition of s_PUB proteins compared to thioredoxins of the YbbN family.** Conserved domains were detected using the ncbi CDD online search tool. Proteins are represented as thick black lines with coordinates, aligned to the position of the shared thioredoxin domain and ordered with respect to their inferred phylogenetic relationships. Domains are represented by colored boxes with absolute coordinates, the color legend is provided at the top right of the panel. **(B) Unrooted maximum likelihood consensus phylogenetic tree of s_PUB and thioredoxin proteins.** The supporting multiple alignment is restricted to the shared amino acids in and around the thioredoxin domain. Ultrafast bootstrap branch support values (1000 replicates) are indicated except when lower than 70% or as a thick line when value is maximum. The number of amino acid positions in the corresponding sequence alignment, the substitution model and parameters used for IQ-TREE reconstruction as well as the branch length scale are indicated at the top right. Protein identifiers indicate the source and the reference separated by a pipe symbol. Leaves corresponding to proteins reported in Stork et al. 2012 are in bold. For each protein, the predicted cellular localisation is tagged by a S for “Signal peptide” and a T for “bipartite Transit sequence” as predicted by Hectar in blue and ASAfind in red. Clades corresponding to proteins putatively implicated in SELMA are shadowed in gray.

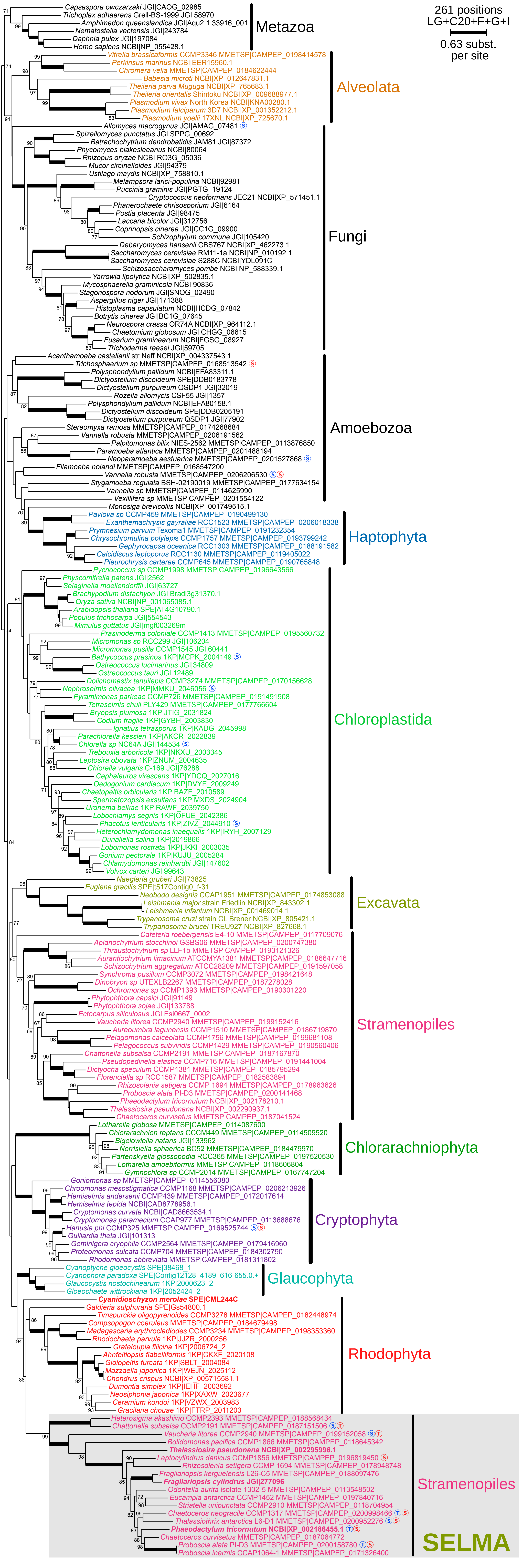

**Supplementary fig. S13. Unrooted maximum likelihood consensus phylogenetic tree of Ubx proteins.** Ultrafast bootstrap branch support values (1000 replicates) are indicated except when lower than 70% or as a thick line when value is maximum. The number of amino acid positions in the corresponding sequence alignment, the substitution model and parameters used for IQ-TREE reconstruction as well as the branch length scale are indicated at the top right. Protein identifiers indicate the source and the reference separated by a pipe symbol. Leaves corresponding to proteins reported in Stork et al. 2012 are in bold. For each protein, the predicted cellular localisation is tagged by a S for "Signal peptide" and a T for "bipartite Transit sequence" as predicted by Hectar in blue and ASAfind in red. Clades corresponding to proteins putatively implicated in SELMA are shadowed in gray.

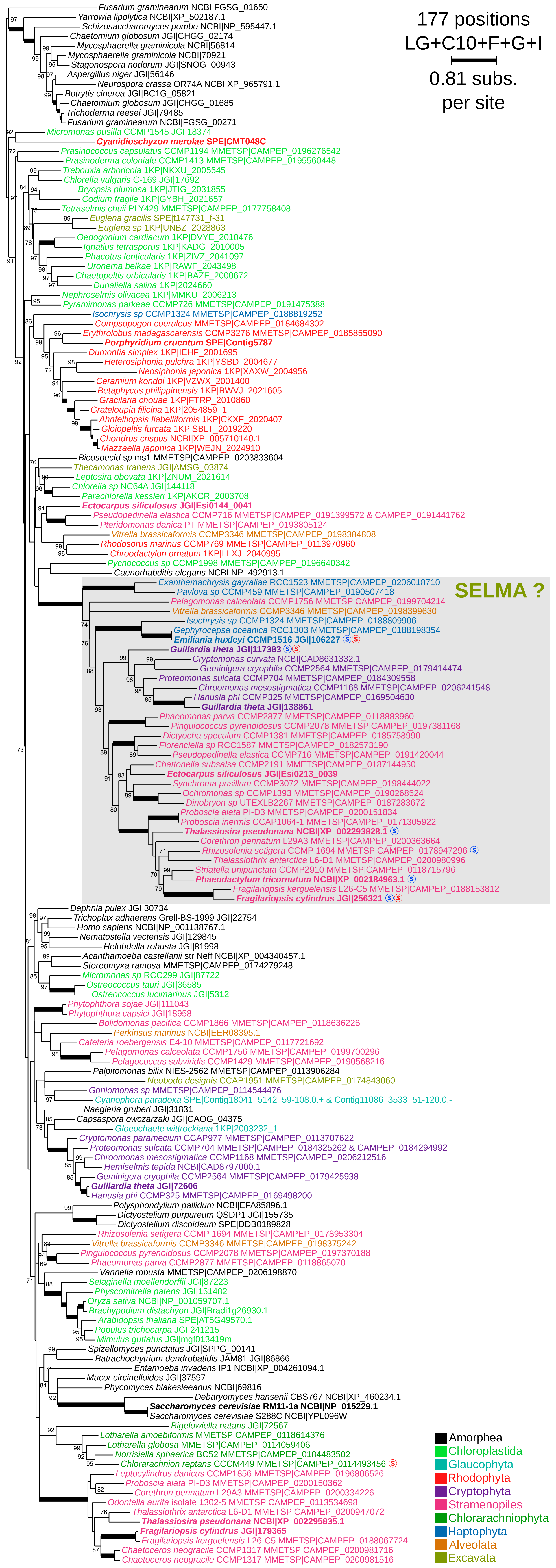

**Supplementary fig. S14. Unrooted maximum likelihood consensus phylogenetic tree of Png1 proteins.** Ultrafast bootstrap branch support values (1000 replicates) are indicated except when lower than 70% or as a thick line when value is maximum. The number of amino acid positions in the corresponding sequence alignment, the substitution model and parameters used for IQ-TREE reconstruction as well as the branch length scale are indicated at the top right. Protein identifiers indicate the source and the reference separated by a pipe symbol. Leaves corresponding to proteins reported in Stork et al. 2012 are in bold. For each protein, the predicted cellular localisation is tagged by a S for "Signal peptide" and a T for "bipartite Transit sequence" as predicted by Hectar in blue and ASAfind in red. Clades corresponding to proteins putatively implicated in SELMA are shadowed in gray.

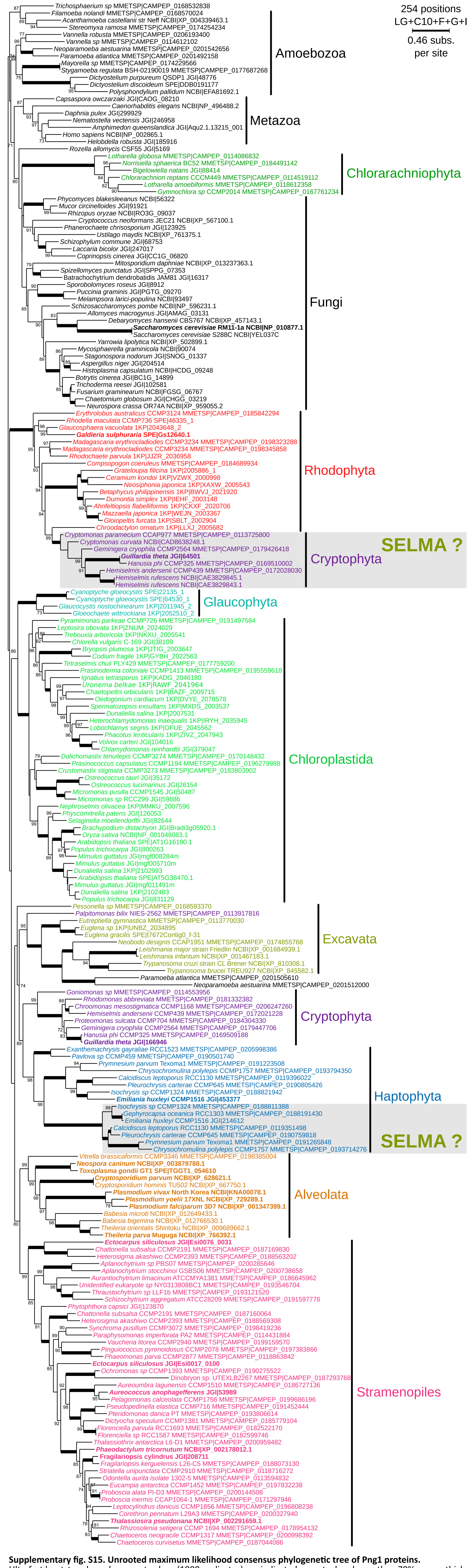

**Supplementary fig. S15. Unrooted maximum likelihood consensus phylogenetic tree of Png1 proteins.**
Ultrafast bootstrap branch support values (1000 replicates) are indicated except when lower than 70% or as a thick line when value is maximum. The number of amino acid positions in the corresponding sequence alignment, the substitution model and parameters used for IQ-TREE reconstruction as well as the branch length scale are indicated at the top right. Protein identifiers indicate the source and the reference separated by a pipe symbol. Leaves corresponding to proteins reported in Stork et al. 2012 are in bold. For each protein, the predicted cellular localisation is tagged by a S for "Signal peptide" and a T for "bipartite Transit sequence" as predicted by Hectar in blue and ASAfind in red. Clades corresponding to proteins putatively implicated in SELMA are shadowed in gray.

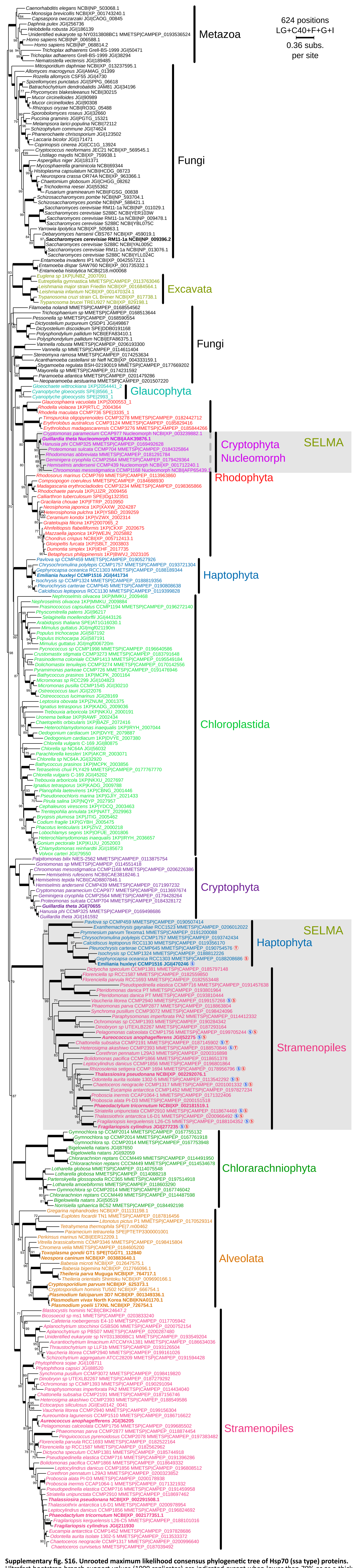

**Supplementary fig. S16. Unrooted maximum likelihood consensus phylogenetic tree of Hsp70 (ssa type) proteins.** Ultrafast bootstrap branch support values (1000 replicates) are indicated except when lower than 70% or as a thick line when value is maximum. The number of amino acid positions in the corresponding sequence alignment, the substitution model and parameters used for IQ-TREE reconstruction as well as the branch length scale are indicated at the top right. Protein identifiers indicate the source and the reference separated by a pipe symbol. Leaves corresponding to proteins reported in Stork et al. 2012 are in bold. For each protein, the predicted cellular localisation is tagged by a S for “Signal peptide” and a T for “bipartite Transit sequence” as predicted by Hectar in blue and ASAfind in red. Clades corresponding to proteins putatively implicated in SELMA are shadowed in gray.

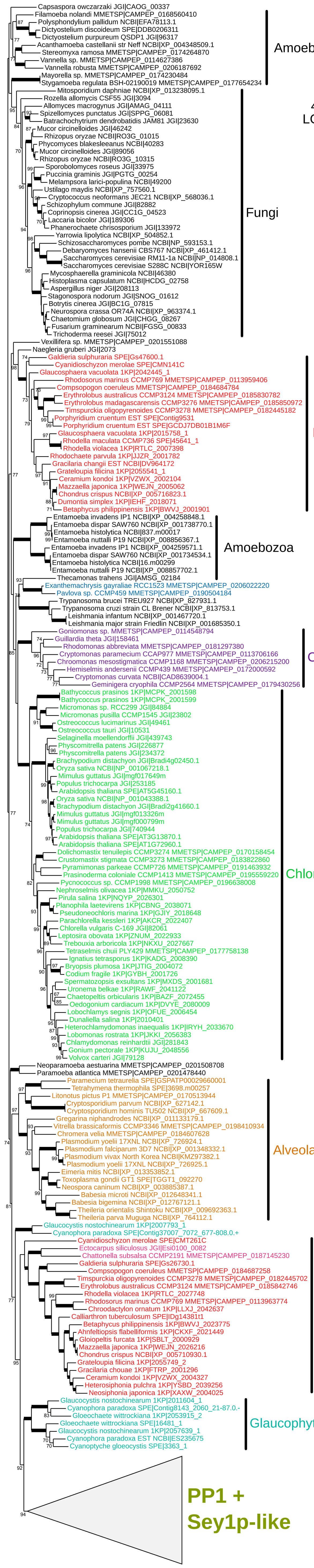

**Supplementary fig. S17-A. Unrooted maximum likelihood consensus phylogenetic trees of PPP1 and Sey1p-like proteins, encompassing the whole diversity of Sey1p-like proteins.**
Ultrafast bootstrap branch support values (1000 replicates) are indicated except when lower than 70% or as a thick line when value is maximum. The number of amino acid positions in the corresponding sequence alignment, the substitution model and parameters used for IQ-TREE reconstruction as well as the branch length scale are indicated at the top right. Protein identifiers indicate the source and the reference separated by a pipe symbol. Leaves corresponding to proteins reported in Stork et al. 2012 are in bold. For each protein, the predicted cellular localisation is tagged by a S for "Signal peptide" and a T for "bipartite Transit sequence" as predicted by Hectar in blue and ASAfind in red. Clades corresponding to proteins putatively implicated in SELMA are shadowed in gray.

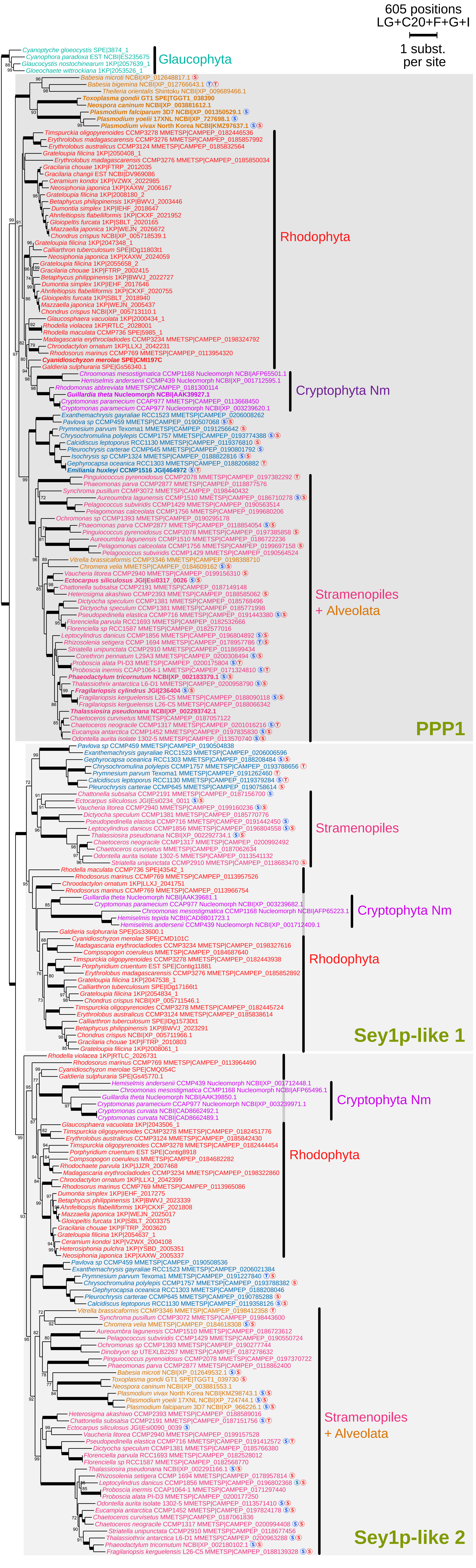

**Supplementary fig. S17-B. Maximum likelihood consensus phylogenetic trees of PPP1 and Sey1p-like proteins, tree restricted to PPP1 homologs and their the closest Sey1p-like similar proteins.** Glaucophyta are used as outgroup. Ultrafast bootstrap branch support values (1000 replicates) are indicated except when lower than 70% or as a thick line when value is maximum. The number of amino acid positions in the corresponding sequence alignment, the substitution model and parameters used for IQ-TREE reconstruction as well as the branch length scale are indicated at the top right. Protein identifiers indicate the source and the reference separated by a pipe symbol. Leaves corresponding to proteins reported in Stork et al. 2012 are in bold. For each protein, the predicted cellular localisation is tagged by a S for "Signal peptide" and a T for "bipartite Transit sequence" as predicted by Hectar in blue and ASAfind in red. Clades corresponding to proteins putatively implicated in SELMA are shadowed in gray.

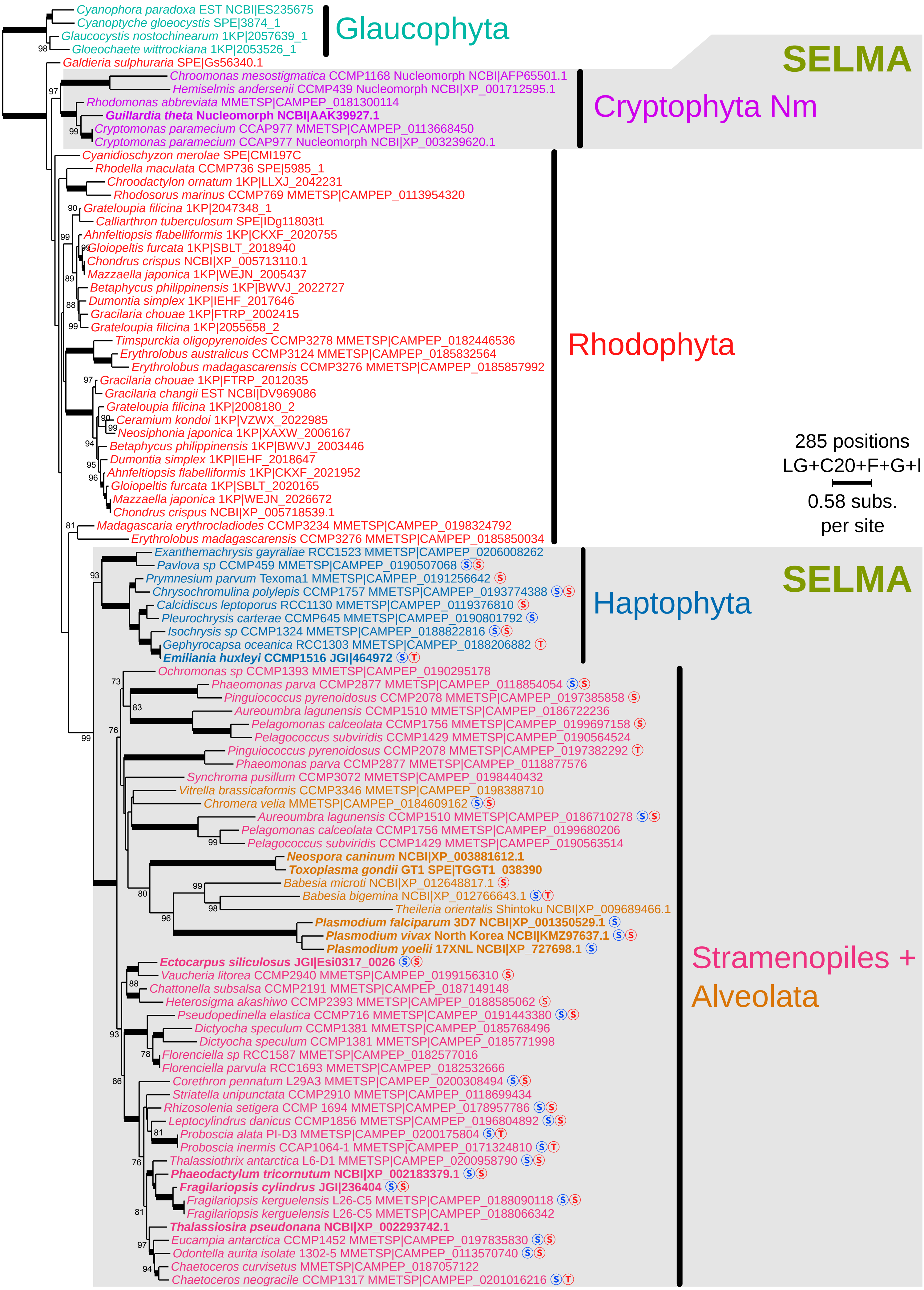

**Supplementary fig. S17-C. Maximum likelihood consensus phylogenetic trees of PPP1 and Sey1p-like proteins, tree restricted to PPP1 protein homologs only.**
Glaucophyta are used as outgroup. Ultrafast bootstrap branch support values (1000 replicates) are indicated except when lower than 70% or as a thick line when value is maximum. The number of amino acid positions in the corresponding sequence alignment, the substitution model and parameters used for IQ-TREE reconstruction as well as the branch length scale are indicated at the top right. Protein identifiers indicate the source and the reference separated by a pipe symbol. Leaves corresponding to proteins reported in Stork et al. 2012 are in bold. For each protein, the predicted cellular localisation is tagged by a S for "Signal peptide" and a T for "bipartite Transit sequence" as predicted by Hectar in blue and ASAfind in red. Clades corresponding to proteins putatively implicated in SELMA are shadowed in gray.

**Supplementary Table S1. List of all potential SELMA components studied in this work.**
Proteins for which a complete analysis could not be performed are grayed and the reason for the failure is specified. Details provided concerning proteins for which a phylogenetic tree is presented in this publication are: the strategy used to trim the final protein alignment, the length of the final alignment, the program and parameters used to produce the phylogenetic tree, the reference of the figures where the corresponding trees can be found, and a summary of the presence/absence status of each component in CASH phyla.

| | Tree Reconstruction | | | | | | | Component Presence/Absence | | | |
|---|---|---|---|---|---|---|---|---|---|---|---|
| **Functionnal group** | **Protein Name** | **MSA trimming** | **MSA lenght** | **Rec. program** | **Bootstrap method** | **Substitution Model** | **Figure** | ***Cryptophyta*** | ***Haptophyta*** | ***Stramenopiles*** | ***Alveolata*** |
| **Derlins** | Der1+Dfm1 | trimal gt 0.7 | 198 | iqtree | 1000x UFboot | LG+C20+F+G+I | sup fig. S01 | N/A | N/A | N/A | N/A |
| | Der1 | trimal gappyout | 203 | iqtree | 1000x UFboot | LG+C40+F+G+I | figure 2a sup fig. S02 | + | + | + | + |
| | Dfm1 | trimal gappyout | 211 | iqtree | 1000x UFboot | LG+G+I | figure 2b sup fig. S03 | + | + | + | + |
| **Ubiquitination** | PUBL (Ubiquitin) | manual | 75 | iqtree | 1000x UFboot | LG+C10+F+G+I | sup fig. S04 | + | + | + | + |
| | Uba1 (E1) | trimal gt 0.9 | 661 | iqtree | 1000x UFboot | LG+C40+F+G+I | sup fig. S05 | + | + | + | + |
| | Ubc (E2), all isoforms | trimal gt 0.7 | 164 | iqtree | 1000x alrt | LG+G+I | figure 3a | N/A | N/A | N/A | N/A |
| | UbcX (E2) | trimal gt 0.6 | 152 | iqtree | 1000x UFboot | LG+C10+F+G+I | figure 3b sup fig. S06 | + | + | + | + |
| | Hrd1 (E3) | trimal gt 0.8 | 322 | iqtree | 1000x UFboot | LG+C20+F+G+I | sup fig. S07 | + | - | - | - |
| | ptE3P | Unresolved phylogenetic tree | | | | | | - | - | + | - |
| | ptDUP | trimal gt 0.9 | 232 | iqtree | 1000x UFboot | LG+C20+F+G+I | sup fig. S08 | - | + | + | + |
| | Hrd3 | No potential SELMA paralog | | | | | | N/A | N/A | N/A | N/A |
| | Doa 10 | No potential SELMA paralog | | | | | | N/A | N/A | N/A | N/A |
| **Cgc48 complex** | Cdc48-1 & 2 | trimal gappyout | 670 | iqtree | 1000x UFboot | LG+C20+F+G+I | figure 4a sup fig. S09 | + | + | + | + |
| | | | | | | | | + | + | + | - |
| | Npl4 | trimal gappyout | 200 | iqtree | 1000x UFboot | LG+C20+F+G+I | figure 4b sup fig. S10 | + | + | + | + |
| | Ufd1 | trimal gappyout | 301 | iqtree | 1000x UFboot | LG+C20+F+G+I | figure 4c sup fig. S11 | + | + | + | + |
| | PUB | trimal gt 0.6 | 123 | iqtree | 1000x UFboot | LG+C10+F+G+I | sup fig. S12 | - | - | + | + |
| | UBX | trimal gt 0.8 | 261 | iqtree | 1000x UFboot | LG+C20+F+G+I | sup fig. S13 | - | - | + | - |
| **Accessories** | Png1 | trimal gt 0.8 | 177 | iqtree | 1000x UFboot | LG+C10+F+G+I | sup fig. S14 | + | + | + | + |
| | Rad23 | trimal gt 0.7 | 254 | iqtree | 1000x UFboot | LG+C10+F+G+I | sup fig. S15 | + | + | - | - |
| | Hsp70 | trimal gt 0.8 | 624 | iqtree | 1000x UFboot | LG+C40+F+G+I | sup fig. S16 | + | + | + | - |
| | PPP1 | trimal gt 0.7 | 428 | iqtree | 1000x UFboot | LG+C10+F+G+I | sup fig. S17a | + | + | + | + |
| | | trimal gt 0.6 | 605 | iqtree | 1000x UFboot | LG+C20+F+G+I | sup fig. S17b | + | + | + | + |
| | | trimal gt 0.8 | 285 | iqtree | 1000x UFboot | LG+C20+F+G+I | sup fig. S17c | + | + | + | + |
| | Hsp40 | No potential SELMA paralog | | | | | | N/A | N/A | N/A | N/A |
| | sDPC | References in Stork et al. 2004 are not related | | | | | | N/A | N/A | N/A | N/A |

**UFBoot**: ultrafast bootstrap approximation (Minh et al., 2013; Hoang et al., 2018)
**alrt**: SH-like approximate likelihood ratio test (Guindon et al., 2010)

**Supplementary Table S2. List of protein sequence identifiers corresponding to ERAD and SELMA components.**
The species set presented in this table is the same as in Table S1 from Stork et al. 2014.

| | Derlins | | | |
|---|---|---|---|---|
| | Der1 | | Dfm1 | |
| | ERAD | SELMA | ERAD | SELMA |
| **Saccharomyces cerevisiae S288C** | NCBI\|NP_009760.1 | | NCBI\|NP_010699.1 | |
| **Saccharomyces cerevisiae RM11-1a** | NCBI\|YBR201W | | NCBI\|YDR411C | |
| **Cyanidioschyzon merolae** | SPE\|CMK163C | | SPE\|CMI159C | |
| **Galdieria sulphuraria** | SPE\|Gs58360.1 | | SPE\|Gs21850.1 | |
| **Porphyridium cruentum** | SPE\|Contig13518 | | ND | |
| **Calliarthron tuberculosum** | SPE\|IDg11638t1 | | SPE\|IDg9967t1 | |
| **Chondrus crispus** | ND | | ND | |
| **Arabidopsis thaliana** | SPE\|AT4G04860.1 | | SPE\|AT4G29330.1 | |
| **Chlamydomonas reinhardtii** | JGI\|287561 | | JGI\|184920 | |
| **Ostreococcus tauri** | JGI\|20852 | | JGI\|16216 | |
| **Volvox carteri** | JGI\|33117 | | JGI\|82188 | |
| **Cyanophora paradoxa** | SPE\|Contig53249_10566_3309-3554.2.- | | ND | |
| **Glaucocystis nostochinearum** | NCBI\|EC124944+1_1<br>1KP\|2054429_2 | | 1KP\|2054764_1 | |
| **Phaeodactylum tricornutum** | NCBI\|XP_002181289.1 | NCBI\|XP_002180594.1 | NCBI\|XP_002181749.1 | NCBI\|XP_002176640.1 |
| **Thalassiosira pseudonana** | NCBI\|XP_002291637.1 | NCBI\|XP_002289517.1 | NCBI\|XP_002291126.1<br>NCBI\|XP_002294055.1 | NCBI\|XP_002293570.1 |
| **Fragilariopsis cylindrus** | JGI\|196204 | JGI\|143375 | JGI\|272386 | JGI\|270130 |
| **Ectocarpus siliculosus** | JGI\|Esi0334_0030 | JGI\|Esi0048_0118 | JGI\|Esi0180_0017 | JGI\|Esi0149_0027 |
| **Aureococcus anophagefferens** | JGI\|5820 | JGI\|71775 | JGI\|16915 | JGI\|64463 |
| **Emiliania huxleyi CCMP1516** | JGI\|99592 | JGI\|96871 | JGI\|420318 | JGI\|204137 |
| **Prymnesium parvum Texoma1** | MMETSP\|CAMPEP_0191223128 | MMETSP\|CAMPEP_0191208100 | MMETSP\|CAMPEP_0191240270 | MMETSP\|CAMPEP_0191218506 |
| **Guillardia theta** | JGI\|114417 | NCBI\|AAK39810.1 | JGI\|165822 | JGI\|146795 |
| **Hemiselmis andersenii CCMP439** | ND | NCBI\|XP_001712217.1 | MMETSP\|CAMPEP_0172017056 | MMETSP\|CAMPEP_0172000694 |
| **Vitrella brassicaformis CCMP3346** | ND | MMETSP\|CAMPEP_0198373108 | MMETSP\|CAMPEP_0198389648 | MMETSP\|CAMPEP_0198405618 |
| **Chromera velia** | ND | MMETSP\|CAMPEP_0184612890 | MMETSP\|CAMPEP_0184615242 | MMETSP\|CAMPEP_0184618762 |
| **Plasmodium falciparum 3D7** | NCBI\|XP_001348827.1 | NCBI\|XP_001351212.1 | NCBI\|XP_001347601.1 | NCBI\|XP_001348672.1 |
| **Plasmodium vivax North Korea** | NCBI\|KMZ98033.1 | NCBI\|KMZ99920.1 | NCBI\|KNA00679.1 | NCBI\|KMZ97895.1 |
| **Plasmodium yoelii 17XNL** | NCBI\|XP_730815.1 | NCBI\|XP_730154.1 | NCBI\|XP_731117.1 | NCBI\|XP_727533.1 |
| **Cryptosporidium parvum** | NCBI\|XP_626259.1 | ND | NCBI\|XP_627700.1 | ND |
| **Theileria parva Muguga** | NCBI\|XP_764690.1 | ND | NCBI\|XP_764400.1 | ND |
| **Neospora caninum** | NCBI\|XP_003885528.1 | ND | NCBI\|XP_003879645.1 | NCBI\|XP_003884723.1 |
| **Toxoplasma gondii GT1** | SPE\|TGGT1_097970 | ND | SPE\|TGGT1_074590 | SPE\|TGGT1_068440 |

| | Ubiquitinylation | | | |
|---|---|---|---|---|
| | Ubiquitin | | Uba1 | |
| | Ubiquitin | PUBL | ERAD | SELMA |
| **Saccharomyces cerevisiae S288C** | NCBI\|YLL039C | | NCBI\|NP_012712.1 | |
| **Saccharomyces cerevisiae RM11-1a** | NCBI\|NP_013061.1 | | NCBI\|YKL210W | |
| **Cyanidioschyzon merolae** | ND | | SPE\|CMI034C | |
| **Galdieria sulphuraria** | SPE\|Gs13790.1 | | SPE\|Gs42920.1 | |
| **Porphyridium cruentum** | ND | | SPE\|Contig13481 | |
| **Calliarthron tuberculosum** | ND | | SPE\|IDg16397t1 | |
| **Chondrus crispus** | SPE\|CO649786 | | NCBI\|XP_005710202.1 | |
| **Arabidopsis thaliana** | SPE\|AT4G05050.1 | | SPE\|AT2G30110.1 | |
| **Chlamydomonas reinhardtii** | JGI\|140045 | | JGI\|191903 | |
| **Ostreococcus tauri** | ND | | JGI\|351 | |
| **Volvox carteri** | JGI\|75856 | | JGI\|79505 | |
| **Cyanophora paradoxa** | NCBI\|EC658194 | | SPE\|Contig53781_10949_8218-8269.0.+ | |
| **Glaucocystis nostochinearum** | 1KP\|2005656_1 | | 1KP\|2058050_1 | |
| **Phaeodactylum tricornutum** | XP_002180197.1 | NCBI\|XP_002178822.1 | NCBI\|XP_002181983.1 | ND |
| **Thalassiosira pseudonana** | ND | ND | NCBI\|XP_002288513.1 | NCBI\|XP_002289167.1 |
| **Fragilariopsis cylindrus** | JGI\|268616 | JGI\|270635 | JGI\|205276 | JGI\|191079 |
| **Ectocarpus siliculosus** | JGI\|Esi0302_0019 | JGI\|Esi0023_0071 | JGI\|Esi0049_0010 | JGI\|Esi0188_0036 |
| **Aureococcus anophagefferens** | JGI\|37580 | ND | JGI\|54097 | ND |
| **Emiliania huxleyi CCMP1516** | JGI\|358059 | JGI\|428400 | JGI\|425780<br>JGI\|467973 | JGI\|459406 |
| **Prymnesium parvum Texoma1** | ND | ND | MMETSP\|CAMPEP_0191207018<br>MMETSP\|CAMPEP_0191201276 | MMETSP\|CAMPEP_0191222846 |
| **Guillardia theta** | JGI\|156153 | JGI\|155024 | JGI\|64494<br>JGI\|160232 | JGI\|157145 |
| **Hemiselmis andersenii CCMP439** | NCBI\|XP_001712639.1 | NCBI\|XP_001712639.1 | MMETSP\|CAMPEP_0172004026<br>MMETSP\|CAMPEP_0172004458 | MMETSP\|CAMPEP_0172019826 |
| **Vitrella brassicaformis CCMP3346** | ND | MMETSP\|CAMPEP_0198373422 | MMETSP\|CAMPEP_0198374660 | ND |
| **Chromera velia** | ND | MMETSP\|CAMPEP_0184617942 | MMETSP\|CAMPEP_0184606190 | ND |
| **Plasmodium falciparum 3D7** | NCBI\|XP_001350526.1 | SPE\|PF3D7_0815700 | NCBI\|XP_001350655.1 | NCBI\|XP_001350063.1 |
| **Plasmodium vivax North Korea** | ND | P01 SPE\|P01PVP01_0519600 | NCBI\|KMZ96829.1 | NCBI\|KMZ98190.1 |
| **Plasmodium yoelii 17XNL** | ND | SPE\|PY17X_0715300 | NCBI\|XP_729708.1 | NCBI\|XP_729678.1<br>XP_727063.1 |
| **Cryptosporidium parvum** | ND | ND | NCBI\|XP_625819.1 | ND |
| **Theileria parva Muguga** | NCBI\|XP_766242.1 | NCBI\|XP_764711.1 | NCBI\|XP_765255.1 | NCBI\|XP_764897.1 |
| **Neospora caninum** | NCBI\|XP_003885682.1 | NCBI\|XP_003880173.1 | NCBI\|XP_003881216.1 | NCBI\|XP_003885382.1 |
| **Toxoplasma gondii** | SPE\|TGGT1_030830 | SPE\|TGME49_223125 | SPE\|TGGT1_032760 | SPE\|TGGT1_092180 |

| | Ubiquitinylation | | | |
|---|---|---|---|---|
| | UbcX | | Hrd1 | |
| | HOST | SELMA | ERAD | SELMA |
| **Saccharomyces cerevisiae S288C** | - | | NCBI\|NP_014630.1 | |
| **Saccharomyces cerevisiae RM11-1a** | - | | NCBI\|YOL013C | |
| **Cyanidioschyzon merolae** | SPE\|CMQ038C | | SPE\|CMG190C | |
| **Galdieria sulphuraria** | SPE\|Gs43090.1 | | SPE\|Gs07260.1 | |
| **Porphyridium cruentum** | SPE\|GCDJ7DB01DEEUH | | SPE\|Contig6247 | |
| **Calliarthron tuberculosum** | SPE\|IDg12630t1 | | ND | |
| **Chondrus crispus** | NCBI\|XP_005710048.1 | | NCBI\|XP_005711321.1 | |
| **Arabidopsis thaliana** | SPE\|AT1G75440.1 SPE\|AT5G42990.1 | | SPE\|AT3G16090.1 | |
| **Chlamydomonas reinhardtii** | JGI\|377402 | | JGI\|122373 | |
| **Ostreococcus tauri** | ND | | JGI\|29041 | |
| **Volvox carteri** | JGI\|95153 | | JGI\|67392 | |
| **Cyanophora paradoxa** | NCBI\|ES231497 | | SPE\|Contig388_26_64-577.0.- | |
| **Glaucocystis nostochinearum** | 1KP\|2008484_1 | | 1KP\|2010848_1 | |
| **Phaeodactylum tricornutum** | ND | NCBI\|XP_002178369.1 | NCBI\|XP_002184604.1 | ND |
| **Thalassiosira pseudonana** | ND | NCBI\|XP_002291866.1 | NCBI\|XP_002293805.1 | ND |
| **Fragilariopsis cylindrus** | ND | JGI\|273521 | JGI\|192595 | ND |
| **Ectocarpus siliculosus** | JGI\|Esi0345_0022 | JGI\|Esi0021_0112 | JGI\|Esi0009_0146 | ND |
| **Aureococcus anophagefferens** | ND | ND | JGI\|1818 | ND |
| **Emiliania huxleyi CCMP1516** | ND | JGI\|452424 | JGI\|248733 | ND |
| **Prymnesium parvum Texoma1** | ND | MMETSP\|CAMPEP_0191198104 | MMETSP\|CAMPEP_0191225908 | ND |
| **Guillardia theta** | JGI\|156286 | JGI\|137331 | JGI\|91104 | NCBI\|CAC27064.1 |
| **Hemiselmis andersenii CCMP439** | MMETSP\|CAMPEP_0172012466 | MMETSP\|CAMPEP_0172034214 | MMETSP\|CAMPEP_0172003904 NCBI\|ZP_08256689.1 | NCBI\|XP_001712433.1 |
| **Vitrella brassicaformis CCMP3346** | MMETSP\|CAMPEP_0198375750 | ND | MMETSP\|CAMPEP_0198381954 | ND |
| **Chromera velia** | ND | MMETSP\|CAMPEP_0184605158 | ND | ND |
| **Plasmodium falciparum 3D7** | NCBI\|XP_001348301.1 | NCBI\|XP_001350169.1 | NCBI\|XP_001348389.1 | ND |
| **Plasmodium vivax North Korea** | NCBI\|KMZ97414.1 | NCBI\|KMZ98374.1 | NCBI\|KMZ97468.1 | ND |
| **Plasmodium yoelii 17XNL** | NCBI\|XP_726132.1 | NCBI\|XP_725503.1 | NCBI\|XP_729657.1 | ND |
| **Cryptosporidium parvum** | ND | ND | NCBI\|XP_627156.1 | ND |
| **Theileria parva Muguga** | ND | NCBI\|XP_765206.1 | NCBI\|XP_766176.1 | ND |
| **Neospora caninum** | NCBI\|XP_003882985.1 | ND | NCBI\|XP_003882223.1 | ND |
| **Toxoplasma gondii** | SPE\|TGGT1_010800 | SPE\|TGME49_295990 | SPE\|TGGT1_006600 | ND |

| | Ubiquitinylation | |
|---|---|---|
| | ptDUP | |
| | HOST | SELMA |
| **Saccharomyces cerevisiae S288C** | NCBI\|YMR304W | |
| **Saccharomyces cerevisiae RM11-1a** | NCBI\|NP_014033.1 | |
| **Cyanidioschyzon merolae** | SPE\|CMR384C | |
| **Galdieria sulphuraria** | SPE\|Gs28470.1 | |
| **Porphyridium cruentum** | SPE\|Contig9736 | |
| **Calliarthron tuberculosum** | ND | |
| **Chondrus crispus** | NCBI\|XP_005710625.1 | |
| **Arabidopsis thaliana** | SPE\|AT5G06600.3 SPE\|AT3G11910.1 | |
| **Chlamydomonas reinhardtii** | JGI\|121621 | |
| **Ostreococcus tauri** | JGI\|15481 | |
| **Volvox carteri** | JGI\|40312 | |
| **Cyanophora paradoxa** | SPE\|Contig8309_2247_20-62.0.- | |
| **Glaucocystis nostochinearum** | 1KP\|2012040_1 | |
| **Phaeodactylum tricornutum** | ND | NCBI\|XP_002184102.1 |
| **Thalassiosira pseudonana** | ND | NCBI\|XP_002287998.1 |
| **Fragilariopsis cylindrus** | ND | JGI\|140767 |
| **Ectocarpus siliculosus** | JGI\|Esi0321_0031 | JGI\|Esi0049_0009 |
| **Aureococcus anophagefferens** | ND | JGI\|71110 |
| **Emiliania huxleyi CCMP1516** | JGI\|441425 | ND |
| **Prymnesium parvum Texoma1** | MMETSP\|CAMPEP_0191207114 | MMETSP\|CAMPEP_0191223360 |
| **Guillardia theta** | JGI\|66749 | ND |
| **Hemiselmis andersenii CCMP439** | ND | ND |
| **Vitrella brassicaformis CCMP3346** | MMETSP\|CAMPEP_0198385538 | MMETSP\|CAMPEP_0198419118 |
| **Chromera velia** | MMETSP\|CAMPEP_0184642498 | ND |
| **Plasmodium falciparum 3D7** | NCBI\|XP_001349163.1 | ND |
| **Plasmodium vivax North Korea** | NCBI\|KNA02060.1 | ND |
| **Plasmodium yoelii 17XNL** | NCBI\|XP_729206.1 | ND |
| **Cryptosporidium parvum** | NCBI\|XP_627060.1 | ND |
| **Theileria parva Muguga** | NCBI\|XP_763522.1 | ND |
| **Neospora caninum** | NCBI\|XP_003881127.1 | ND |
| **Toxoplasma gondii** | SPE\|TGGT1_033890 | ND |

| | Cdc48 complex | | |
|---|---|---|---|
| | Cdc48 | | |
| | ERAD | SELMA-1 | SELMA-2 |
| Saccharomyces cerevisiae S288C | NCBI\|NP_010157.1 | | |
| Saccharomyces cerevisiae RM11-1a | NCBI\|YDL126C | | |
| Cyanidioschyzon merolae | SPE\|CML023C | | |
| Galdieria sulphuraria | SPE\|Gs41710.1 | | |
| Porphyridium cruentum | ND | | |
| Calliarthron tuberculosum | SPE\|IDg2684t1 | | |
| Chondrus crispus | NCBI\|XP_005715271.1 | | |
| Arabidopsis thaliana | SPE\|AT5G03340.1 | | |
| Chlamydomonas reinhardtii | JGI\|134171 | | |
| Ostreococcus tauri | JGI\|14760 | | |
| Volvox carteri | JGI\|78972 | | |
| Cyanophora paradoxa | SPE\|Contig38151_7626_1389-1425.0.- SPE\|Contig40063_9154_30-223.1.- | | |
| Glaucocystis nostochinearum | 1KP\|2007908_3 | | |
| Phaeodactylum tricornutum | NCBI\|XP_002185883.1 | NCBI\|XP_002181088.1 | NCBI\|XP_002178361.1 |
| Thalassiosira pseudonana | NCBI\|XP_002290761.1 NCBI\|XP_002286617.1 | NCBI\|XP_002290667.1 | NCBI\|XP_002289352.1 |
| Fragilariopsis cylindrus | JGI\|260828 | JGI\|275920 | JGI\|276489 |
| Ectocarpus siliculosus | JGI\|Esi0010_0010 | JGI\|Esi0014_0217 | JGI\|Esi0057_0096 |
| Aureococcus anophagefferens | JGI\|69630 | JGI\|550 | JGI\|36060 |
| Emiliania huxleyi CCMP1516 | JGI\|439359 | JGI\|421356 | JGI\|461250 |
| Prymnesium parvum Texoma1 | MMETSP\|CAMPEP_0191200710 | MMETSP\|CAMPEP_0191256432 | MMETSP\|CAMPEP_0191256272 |
| Guillardia theta | JGI\|98435 | NCBI\|AAK39773.1 | JGI\|161858 |
| Hemiselmis andersenii CCMP439 | MMETSP\|CAMPEP_0172011704 | NCBI\|XP_001712323.1 | MMETSP\|CAMPEP_0172006232 |
| Vitrella brassicaformis CCMP3346 | MMETSP\|CAMPEP_0198372358 | MMETSP\|CAMPEP_0198412374 | ND |
| Chromera velia | MMETSP\|CAMPEP_0184641108 | MMETSP\|CAMPEP_0184617182 | ND |
| Plasmodium falciparum 3D7 | NCBI\|XP_966179.1 | NCBI\|XP_001349023.1 | ND |
| Plasmodium vivax North Korea | NCBI\|KMZ98642.1 | NCBI\|KNA02472.1 | ND |
| Plasmodium yoelii 17XNL | NCBI\|XP_723826.1 | NCBI\|XP_726304.1 | ND |
| Cryptosporidium parvum | NCBI\|XP_627893.1 | ND | ND |
| Theileria parva Muguga | NCBI\|XP_766458.1 | NCBI\|XP_763518.1 | ND |
| Neospora caninum | NCBI\|XP_003883696.1 | NCBI\|XP_003879979.1 | ND |
| Toxoplasma gondii GT1 | SPE\|TGGT1_112260 | SPE\|TGGT1_064680 | ND |

| | Cdc48 complex | | | |
|---|---|---|---|---|
| | Npl4 | | Ufd1 | |
| | ERAD | SELMA | ERAD | SELMA |
| Saccharomyces cerevisiae S288C | NCBI\|NP_009729.1 | | NCBI\|NP_011562.1 | |
| Saccharomyces cerevisiae RM11-1a | NCBI\|YBR170C | | NCBI\|YGR048W | |
| Cyanidioschyzon merolae | SPE\|CMH273C | | ND | |
| Galdieria sulphuraria | SPE\|Gs19810.1 | | SPE\|Gs43460.1 | |
| Porphyridium cruentum | SPE\|Contig8004 | | SPE\|GCDJ7DB01CJN0L | |
| Calliarthron tuberculosum | ND | | ND | |
| Chondrus crispus | ND | | NCBI\|XP_005710167.1 | |
| Arabidopsis thaliana | SPE\|AT2G47970.1 | | SPE\|AT4G38930.2 SPE\|AT2G21270.2 | |
| Chlamydomonas reinhardtii | ND | | JGI\|394624 | |
| Ostreococcus tauri | JGI\|16895 | | JGI\|5039 | |
| Volvox carteri | JGI\|69322 | | JGI\|102621 | |
| Cyanophora paradoxa | SPE\|Contig7675_1083_3659-3761.0.- NCBI\|FG948221 | | SPE\|Contig8908_2555_1630-1719.0.+ | |
| Glaucocystis nostochinearum | 1KP\|2057612_2 | | 1KP\|2010481_2 | |
| Phaeodactylum tricornutum | NCBI\|XP_002186415.1 | NCBI\|XP_002179175.1 | NCBI\|XP_002178490.1 | NCBI\|XP_002184020.1 |
| Thalassiosira pseudonana | NCBI\|XP_002287807.1 | NCBI\|XP_002290919.1 | NCBI\|XP_002288569.1 | NCBI\|XP_002289664.1 |
| Fragilariopsis cylindrus | JGI\|247186 | JGI\|207358 | JGI\|195024 | JGI\|219202 |
| Ectocarpus siliculosus | JGI\|Esi0070_0117 | JGI\|Esi0041_0124 | JGI\|Esi0009_0093 | JGI\|Esi0018_0041 |
| Aureococcus anophagefferens | ND | ND | JGI\|60034 | ND |
| Emiliania huxleyi CCMP1516 | JGI\|454904 | JGI\|99124 | JGI\|467386 | JGI\|AKBS2441-I01.x1d-F |
| Prymnesium parvum Texoma1 | MMETSP\|CAMPEP_0191210290 | MMETSP\|CAMPEP_0191221030 | MMETSP\|CAMPEP_0191206806 MMETSP\|CAMPEP_0191219060 | MMETSP\|CAMPEP_0191219060 |
| Guillardia theta | JGI\|133675 | JGI\|98945 | JGI\|65655 | NCBI\|AAF24006.1 |
| Hemiselmis andersenii CCMP439 | MMETSP\|CAMPEP_0172013564 | MMETSP\|CAMPEP_0172002030 | MMETSP\|CAMPEP_0172007224 | NCBI\|XP_001712332.1 |
| Vitrella brassicaformis CCMP3346 | | MMETSP\|CAMPEP_0198414444 | MMETSP\|CAMPEP_0198414192 | MMETSP\|CAMPEP_0198380478 |
| Chromera velia | MMETSP\|CAMPEP_0184612676 | ND | MMETSP\|CAMPEP_0184611272 | MMETSP\|CAMPEP_0184620216 |
| Plasmodium falciparum 3D7 | NCBI\|XP_001351635.1 | NCBI\|XP_001347517.1 | NCBI\|XP_001348351.1 | NCBI\|XP_001352037.1 |
| Plasmodium vivax North Korea | NCBI\|KNA02280.1 | NCBI\|KNA00736.1 | NCBI\|KMZ97363.1 | NCBI\|KNA00367.1 |
| Plasmodium yoelii 17XNL | NCBI\|XP_725542.1 | ND | NCBI\|XP_729433.1 | ND |
| Cryptosporidium parvum | NCBI\|XP_628374.1 | ND | NCBI\|XP_625717.1 | ND |
| Theileria parva Muguga | NCBI\|XP_765129.1 | ND | NCBI\|XP_765096.1 | ND |
| Neospora caninum | NCBI\|XP_003883798.1 | ND | NCBI\|XP_003883874.1 | NCBI\|XP_003881706.1 |
| Toxoplasma gondii GT1 | SPE\|TGGT1_110310 | ND | SPE\|TGGT1_109070 | GT1 SPE\|TGGT1_037100 |

| | Cdc48 complex | | | |
|---|---|---|---|---|
| | PUB | | UBX | |
| | HOST (Ybbn-like proteins) | SELMA | HOST | SELMA |
| Saccharomyces cerevisiae S288C | ND | | NCBI\|YDL091C | |
| Saccharomyces cerevisiae RM11-1a | ND | | NCBI\|NP_010192.1 | |
| Cyanidioschyzon merolae | ND | | SPE\|CML244C | |
| Galdieria sulphuraria | ND | | SPE\|Gs54800.1 | |
| Porphyridium cruentum | ND | | ND | |
| Calliarthron tuberculosum | ND | | ND | |
| Chondrus crispus | ND | | NCBI\|XP_005715581.1 | |
| Arabidopsis thaliana | ND | | SPE\|AT4G10790.1 | |
| Chlamydomonas reinhardtii | ND | | JGI\|147602 | |
| Ostreococcus tauri | JGI\|7672 | | JGI\|12489 | |
| Volvox carteri | ND | | JGI\|99643 | |
| Cyanophora paradoxa | SPE\|Contig26077_6097_307-493.0.+<br>EST NCBI\|EC663491 | | SPE\|Contig12128_4189_616-655.0.+ | |
| Glaucocystis nostochinearum | 1KP\|2052150_1<br>1KP\|2010547_3 | | 1KP\|2000623_2 | |
| Phaeodactylum tricornutum | ND | NCBI\|XP_002181771.1 | NCBI\|XP_002178210.1 | NCBI\|XP_002186455.1 |
| Thalassiosira pseudonana | ND | NCBI\|XP_002292615.1 | NCBI\|XP_002290937.1 | NCBI\|XP_002295996.1 |
| Fragilariopsis cylindrus | ND | JGI\|180686 | ND | JGI\|277096 |
| Ectocarpus siliculosus | ND | JGI\|Esi0043_0092 | JGI\|Esi0667_0002 | ND |
| Aureococcus anophagefferens | JGI\|36471 | ND | ND | ND |
| Emiliania huxleyi CCMP1516 | JGI\|466974 | ND | ND | ND |
| Prymnesium parvum Texoma1 | MMETSP\|CAMPEP_0191228108 | ND | MMETSP\|CAMPEP_0191232354 | ND |
| Guillardia theta | ND | ND | JGI\|101313 | ND |
| Hemiselmis andersenii CCMP439 | MMETSP\|CAMPEP_0171998694 | ND | MMETSP\|CAMPEP_0172017614 | ND |
| Vitrella brassicaformis CCMP3346 | ND | NCBI\|CEL91887,1 | MMETSP\|CAMPEP_0198414578 | ND |
| Chromera velia | ND | MMETSP\|CAMPEP_0184641012 | MMETSP\|CAMPEP_0184622444 | ND |
| Plasmodium falciparum 3D7 | ND | ND | NCBI\|XP_001352212.1 | ND |
| Plasmodium vivax North Korea | ND | ND | NCBI\|KNA00280.1 | ND |
| Plasmodium yoelii 17XNL | ND | ND | NCBI\|XP_725670.1 | ND |
| Cryptosporidium parvum | ND | ND | ND | ND |
| Theileria parva Muguga | ND | ND | NCBI\|XP_765683.1 | ND |
| Neospora caninum | ND | ND | ND | ND |
| Toxoplasma gondii GT1 | ND | ND | ND | ND |

| | Accessory proteins | | | |
|---|---|---|---|---|
| | Png1 | | Rad23 | |
| | ERAD | SELMA | ERAD | SELMA |
| Saccharomyces cerevisiae S288C | NCBI\|NP_015229.1 | | NCBI\|NP_010877.1 | |
| Saccharomyces cerevisiae RM11-1a | NCBI\|YPL096W | | NCBI\|YEL037C | |
| Cyanidioschyzon merolae | SPE\|CMT048C | | ND | |
| Galdieria sulphuraria | ND | | SPE\|Gs12640.1 | |
| Porphyridium cruentum | SPE\|Contig5787 | | ND | |
| Calliarthron tuberculosum | ND | | ND | |
| Chondrus crispus | NCBI\|XP_005710140.1 | | ND | |
| Arabidopsis thaliana | SPE\|AT5G49570.1 | | SPE\|AT5G38470.1<br>SPE\|AT1G16190.1 | |
| Chlamydomonas reinhardtii | JGI\|146964 | | JGI\|379047 | |
| Ostreococcus tauri | JGI\|36585 | | JGI\|35172 | |
| Volvox carteri | JGI\|103955 | | JGI\|104016 | |
| Cyanophora paradoxa | SPE\|Contig11086_3533_51-120.0.- | | ND | |
| Glaucocystis nostochinearum | 1KP\|2017498_1 | | 1KP\|2011945_2 | |
| Phaeodactylum tricornutum | ND | NCBI\|XP_002184963.1 | NCBI\|XP_002178012.1 | ND |
| Thalassiosira pseudonana | NCBI\|XP_002295835.1 | NCBI\|XP_002293828.1 | NCBI\|XP_002291659.1 | ND |
| Fragilariopsis cylindrus | JGI\|179365 | JGI\|256321 | JGI\|208711 | ND |
| Ectocarpus siliculosus | JGI\|Esi0144_0041 | JGI\|Esi0213_0039 | JGI\|Esi0017_0100<br>JGI\|Esi0076_0031 | ND |
| Aureococcus anophagefferens | ND | ND | JGI\|53989 | ND |
| Emiliania huxleyi CCMP1516 | ND | JGI\|106227 | JGI\|59557<br>JGI\|453377 | JGI\|214612 |
| Prymnesium parvum Texoma1 | ND | MMETSP\|CAMPEP_0191221280 | MMETSP\|CAMPEP_0191223508 | MMETSP\|CAMPEP_0191265848 |
| Guillardia theta | JGI\|72606 | JGI\|138861<br>JGI\|117383 | JGI\|166946 | JGI\|64501 |
| Hemiselmis andersenii CCMP439 | ND | ND | MMETSP\|CAMPEP_0172021228 | MMETSP\|CAMPEP_0172028030 |
| Vitrella brassicaformis CCMP3346 | MMETSP\|CAMPEP_0198384808 | MMETSP\|CAMPEP_0198399630 | MMETSP\|CAMPEP_0198385004 | ND |
| Chromera velia | ND | ND | ND | ND |
| Plasmodium falciparum 3D7 | ND | ND | NCBI\|XP_001347399.1 | ND |
| Plasmodium vivax North Korea | ND | ND | NCBI\|KNA00078.1 | ND |
| Plasmodium yoelii 17XNL | ND | ND | NCBI\|XP_729289.1 | ND |
| Cryptosporidium parvum | ND | ND | NCBI\|XP_628621.1 | ND |
| Theileria parva Muguga | ND | ND | NCBI\|XP_766392.1 | ND |
| Neospora caninum | ND | ND | NCBI\|XP_003879788.1 | ND |

| | Accessory proteins | | | |
|---|---|---|---|---|
| | Hsp70 | | PPP1 | |
| | ERAD | SELMA | ERAD | SELMA |
| **Saccharomyces cerevisiae S288C** | NCBI\|NP_009396.2<br>NCBI\|NP_013076.1 | | ND | |
| **Saccharomyces cerevisiae RM11-1a** | NCBI\|YAL005C<br>NCBI\|YAL005C | | ND | |
| **Cyanidioschyzon merolae** | SPE\|CMP145C | | SPE\|CMI197C | |
| **Galdieria sulphuraria** | SPE\|Gs16120.1 | | SPE\|Gs56340.1 | |
| **Porphyridium cruentum** | ND | | SPE\|GCDJ7DB01A44EB | |
| **Calliarthron tuberculosum** | SPE\|IDg13235t1 | | ND | |
| **Chondrus crispus** | NCBI\|XP_005712413.1 | | NCBI\|XP_005713110.1<br>NCBI\|XP_005718539.1 | |
| **Arabidopsis thaliana** | SPE\|AT1G16030.1 | | ND | |
| **Chlamydomonas reinhardtii** | JGI\|185673 | | ND | |
| **Ostreococcus tauri** | JGI\|22076 | | ND | |
| **Volvox carteri** | JGI\|79550 | | ND | |
| **Cyanophora paradoxa** | ND | | ND | |
| **Glaucocystis nostochinearum** | ND | | ND | |
| **Phaeodactylum tricornutum** | NCBI\|XP_002177351.1 | NCBI\|XP_002181519.1 | ND | NCBI\|XP_002183379.1 |
| **Thalassiosira pseudonana** | NCBI\|XP_002291508.1 | NCBI\|XP_002292076.1 | ND | NCBI\|XP_002293742.1 |
| **Fragilariopsis cylindrus** | JGI\|211930 | JGI\|277235 | ND | JGI\|236404 |
| **Ectocarpus siliculosus** | JGI\|Esi0142_0041 | ND | ND | JGI\|Esi0317_0026 |
| **Aureococcus anophagefferens** | JGI\|36205 | JGI\|52275 | ND | ND |
| **Emiliania huxleyi CCMP1516** | JGI\|441734 | JGI\|470246 | ND | JGI\|464972 |
| **Prymnesium parvum Texoma1** | ND | MMETSP\|CAMPEP_0191200088 | ND | MMETSP\|CAMPEP_0191256642 |
| **Guillardia theta** | JGI\|161592 | NCBI\|AAK39876.1 | ND | NCBI\|AAK39927.1 |
| **Hemiselmis andersenii CCMP439** | MMETSP\|CAMPEP_0171997232 | NCBI\|XP_001712240.1 | ND | NCBI\|XP_001712595.1 |
| **Vitrella brassicaformis CCMP3346** | MMETSP\|CAMPEP_0198415804 | ND | ND | MMETSP\|CAMPEP_0198388710 |
| **Chromera velia** | MMETSP\|CAMPEP_0184605200 | ND | ND | MMETSP\|CAMPEP_0184609162 |
| **Plasmodium falciparum 3D7** | NCBI\|XP_001349336.1 | ND | ND | NCBI\|XP_001350529.1 |
| **Plasmodium vivax North Korea** | NCBI\|KNA01170.1 | ND | ND | NCBI\|KMZ97637.1 |
| **Plasmodium yoelii 17XNL** | NCBI\|XP_726754.1 | ND | ND | NCBI\|XP_727698.1 |
| **Cryptosporidium parvum** | NCBI\|XP_625373.1 | ND | ND | ND |
| **Theileria parva Muguga** | NCBI\|XP_764717.1 | ND | ND | ND |
| **Neospora caninum** | NCBI\|XP_003883640.1 | ND | ND | NCBI\|XP_003881612.1 |

| Species | Domain | Taxonomy | Asembly type | N. proteins | Assembly source | Source link |
|---|---|---|---|---|---|---|
| Acidianus hospitalis W1 | Archaea | Crenarchaeota | genome | 2368 | NCBI | https://www.ncbi.nlm.nih.gov/genbank/ |
| Acidilobus saccharovorans 345-15 | Archaea | Crenarchaeota | genome | 1499 | NCBI | https://www.ncbi.nlm.nih.gov/genbank/ |
| Aeropyrum pernix K1 | Archaea | Crenarchaeota | genome | 1700 | NCBI | https://www.ncbi.nlm.nih.gov/genbank/ |
| Caldivirga maquilingensis IC_167 | Archaea | Crenarchaeota | genome | 1963 | NCBI | https://www.ncbi.nlm.nih.gov/genbank/ |
| Desulfurococcus kamchatkensis | Archaea | Crenarchaeota | genome | 1471 | NCBI | https://www.ncbi.nlm.nih.gov/genbank/ |
| Desulfurococcus mucosus DSM2162 | Archaea | Crenarchaeota | genome | 1345 | NCBI | https://www.ncbi.nlm.nih.gov/genbank/ |
| Hyperthermus butylicus DSM_5456 | Archaea | Crenarchaeota | genome | 1602 | NCBI | https://www.ncbi.nlm.nih.gov/genbank/ |
| Ignicoccus hospitalis KIN4_I | Archaea | Crenarchaeota | genome | 1434 | NCBI | https://www.ncbi.nlm.nih.gov/genbank/ |
| Ignisphaera aggregans DSM17230 | Archaea | Crenarchaeota | genome | 1930 | NCBI | https://www.ncbi.nlm.nih.gov/genbank/ |
| Metallosphaera cuprina Ar-4 | Archaea | Crenarchaeota | genome | 2029 | NCBI | https://www.ncbi.nlm.nih.gov/genbank/ |
| Metallosphaera sedula DSM_5348 | Archaea | Crenarchaeota | genome | 2256 | NCBI | https://www.ncbi.nlm.nih.gov/genbank/ |
| Methanobrevibacter smithii ATCC_35061 | Archaea | Crenarchaeota | genome | 1793 | NCBI | https://www.ncbi.nlm.nih.gov/genbank/ |
| Pyrobaculum aerophilum IM2 | Archaea | Crenarchaeota | genome | 2605 | NCBI | https://www.ncbi.nlm.nih.gov/genbank/ |
| Pyrobaculum arsenaticum DSM 13514 | Archaea | Crenarchaeota | genome | 2299 | NCBI | https://www.ncbi.nlm.nih.gov/genbank/ |
| Pyrobaculum calidifontis JCM 11548 | Archaea | Crenarchaeota | genome | 2149 | NCBI | https://www.ncbi.nlm.nih.gov/genbank/ |
| Pyrobaculum islandicum DSM 4184 | Archaea | Crenarchaeota | genome | 1978 | NCBI | https://www.ncbi.nlm.nih.gov/genbank/ |
| Pyrobaculum sp. 1860 | Archaea | Crenarchaeota | genome | 2827 | NCBI | https://www.ncbi.nlm.nih.gov/genbank/ |
| Pyrolobus fumarii 1A | Archaea | Crenarchaeota | genome | 1967 | NCBI | https://www.ncbi.nlm.nih.gov/genbank/ |
| Staphylothermus hellenicus DSM 12710 | Archaea | Crenarchaeota | genome | 1672 | JGI | https://mycocosm.jgi.doe.gov/ |
| Staphylothermus marinus F1 | Archaea | Crenarchaeota | genome | 1570 | NCBI | https://www.ncbi.nlm.nih.gov/genbank/ |
| Sulfolobus acidocaldarius DSM_639 | Archaea | Crenarchaeota | genome | 2223 | NCBI | https://www.ncbi.nlm.nih.gov/genbank/ |
| Sulfolobus islandicus L.S.2.15 | Archaea | Crenarchaeota | genome | 2737 | NCBI | https://www.ncbi.nlm.nih.gov/genbank/ |
| Sulfolobus solfataricus P2 | Archaea | Crenarchaeota | genome | 2977 | NCBI | https://www.ncbi.nlm.nih.gov/genbank/ |
| Sulfolobus tokodaii str 7 | Archaea | Crenarchaeota | genome | 2825 | NCBI | https://www.ncbi.nlm.nih.gov/genbank/ |
| Thermofilum pendens Hrk_5 | Archaea | Crenarchaeota | genome | 1876 | NCBI | https://www.ncbi.nlm.nih.gov/genbank/ |
| Thermoproteus neutrophilus V24Sta | Archaea | Crenarchaeota | genome | 1966 | NCBI | https://www.ncbi.nlm.nih.gov/genbank/ |
| Thermoproteus tenax Kra 1 | Archaea | Crenarchaeota | genome | 2049 | NCBI | https://www.ncbi.nlm.nih.gov/genbank/ |
| Thermoproteus uzoniensis 768-20 | Archaea | Crenarchaeota | genome | 2186 | NCBI | https://www.ncbi.nlm.nih.gov/genbank/ |
| Thermosphaera aggregans DSM 11486 | Archaea | Crenarchaeota | genome | 1387 | NCBI | https://www.ncbi.nlm.nih.gov/genbank/ |
| Vulcanisaeta distributa DSM14429 | Archaea | Crenarchaeota | genome | 2493 | NCBI | https://www.ncbi.nlm.nih.gov/genbank/ |
| Vulcanisaeta moutnovskia 768-28 | Archaea | Crenarchaeota | genome | 2320 | NCBI | https://www.ncbi.nlm.nih.gov/genbank/ |
| Aciduliprofundum boonei T469 | Archaea | Euryarchaeota | genome | 2949 | NCBI | https://www.ncbi.nlm.nih.gov/genbank/ |
| Archaeoglobus fulgidus DSM_4304 | Archaea | Euryarchaeota | genome | 2420 | NCBI | https://www.ncbi.nlm.nih.gov/genbank/ |
| Archaeoglobus profundus DSM 5631 | Archaea | Euryarchaeota | genome | 1823 | NCBI | https://www.ncbi.nlm.nih.gov/genbank/ |
| Archaeoglobus veneficus SNP6 | Archaea | Euryarchaeota | genome | 2090 | NCBI | https://www.ncbi.nlm.nih.gov/genbank/ |
| Candidatus Methanoregula boonei 6A8 | Archaea | Euryarchaeota | genome | 2450 | NCBI | https://www.ncbi.nlm.nih.gov/genbank/ |
| Candidatus Micrarchaeum acidiphilum ARMAN-2 | Archaea | Euryarchaeota | genome | 1037 | NCBI | https://www.ncbi.nlm.nih.gov/genbank/ |
| Candidatus Nanosalina sp. J07AB43 | Archaea | Euryarchaeota | genome | 1677 | NCBI | https://www.ncbi.nlm.nih.gov/genbank/ |
| Candidatus Nanosalinarum sp. J07AB56 | Archaea | Euryarchaeota | genome | 1407 | NCBI | https://www.ncbi.nlm.nih.gov/genbank/ |
| Candidatus Nitrosoarchaeum koreensis MY1 | Archaea | Euryarchaeota | genome | 1945 | NCBI | https://www.ncbi.nlm.nih.gov/genbank/ |
| Candidatus Nitrosoarchaeum limnia SFB1 | Archaea | Euryarchaeota | genome | 2038 | NCBI | https://www.ncbi.nlm.nih.gov/genbank/ |
| Candidatus Parvarchaeum acidiphilum ARMAN-4 | Archaea | Euryarchaeota | genome | 911 | NCBI | https://www.ncbi.nlm.nih.gov/genbank/ |
| Candidatus Parvarchaeum acidophilus ARMAN-5 | Archaea | Euryarchaeota | genome | 1042 | NCBI | https://www.ncbi.nlm.nih.gov/genbank/ |
| Ferroglobus placidus DSM 10642 | Archaea | Euryarchaeota | genome | 2480 | NCBI | https://www.ncbi.nlm.nih.gov/genbank/ |
| Ferroplasma acidarmanus fer1 | Archaea | Euryarchaeota | genome | 1986 | NCBI | https://www.ncbi.nlm.nih.gov/genbank/ |
| Halalkalicoccus jeotgali B3 | Archaea | Euryarchaeota | genome | 3873 | NCBI | https://www.ncbi.nlm.nih.gov/genbank/ |
| Haloarcula hispanica ATCC 33960 | Archaea | Euryarchaeota | genome | 3859 | NCBI | https://www.ncbi.nlm.nih.gov/genbank/ |
| Haloarcula marismortui ATCC_43049 | Archaea | Euryarchaeota | genome | 4240 | NCBI | https://www.ncbi.nlm.nih.gov/genbank/ |
| Halobacterium salinarum R1 | Archaea | Euryarchaeota | genome | 2749 | NCBI | https://www.ncbi.nlm.nih.gov/genbank/ |
| Haloferax volcanii DS2 | Archaea | Euryarchaeota | genome | 4015 | NCBI | https://www.ncbi.nlm.nih.gov/genbank/ |
| Halogeometricum borinquense DSM 11551 | Archaea | Euryarchaeota | genome | 4005 | NCBI | https://www.ncbi.nlm.nih.gov/genbank/ |
| Halomicrobium mukohataei DSM 12286 | Archaea | Euryarchaeota | genome | 3349 | NCBI | https://www.ncbi.nlm.nih.gov/genbank/ |
| halophilic archaeon DL31 | Archaea | Euryarchaeota | genome | 3476 | NCBI | https://www.ncbi.nlm.nih.gov/genbank/ |
| Halopiger xanaduensis SH-6 | Archaea | Euryarchaeota | genome | 4221 | NCBI | https://www.ncbi.nlm.nih.gov/genbank/ |
| Haloquadratum walsbyi DSM_16790 | Archaea | Euryarchaeota | genome | 2646 | NCBI | https://www.ncbi.nlm.nih.gov/genbank/ |
| Halorhabdus utahensis DSM 12940 | Archaea | Euryarchaeota | genome | 2998 | NCBI | https://www.ncbi.nlm.nih.gov/genbank/ |
| Halorubrum lacusprofundi ATCC 49239 | Archaea | Euryarchaeota | genome | 3560 | NCBI | https://www.ncbi.nlm.nih.gov/genbank/ |
| Haloterrigena turkmenica DSM 5511 | Archaea | Euryarchaeota | genome | 5113 | NCBI | https://www.ncbi.nlm.nih.gov/genbank/ |
| Marine group II euryarchaeote SCGC AB-629-J06 | Archaea | Euryarchaeota | genome | 550 | NCBI | https://www.ncbi.nlm.nih.gov/genbank/ |
| Methanobacterium sp. AL-21 | Archaea | Euryarchaeota | genome | 2493 | NCBI | https://www.ncbi.nlm.nih.gov/genbank/ |
| Methanobrevibacter ruminantium M1 | Archaea | Euryarchaeota | genome | 2217 | NCBI | https://www.ncbi.nlm.nih.gov/genbank/ |
| Methanocaldococcus fervens AG86 | Archaea | Euryarchaeota | genome | 1581 | NCBI | https://www.ncbi.nlm.nih.gov/genbank/ |
| Methanocaldococcus infernus ME | Archaea | Euryarchaeota | genome | 1441 | NCBI | https://www.ncbi.nlm.nih.gov/genbank/ |
| Methanocaldococcus jannaschii DSM 2661 | Archaea | Euryarchaeota | genome | 1786 | NCBI | https://www.ncbi.nlm.nih.gov/genbank/ |
| Methanocaldococcus sp. FS406-22 | Archaea | Euryarchaeota | genome | 1816 | NCBI | https://www.ncbi.nlm.nih.gov/genbank/ |
| Methanocaldococcus vulcanius M7 | Archaea | Euryarchaeota | genome | 1742 | NCBI | https://www.ncbi.nlm.nih.gov/genbank/ |
| Methanocella paludicola SANAE | Archaea | Euryarchaeota | genome | 3004 | NCBI | https://www.ncbi.nlm.nih.gov/genbank/ |
| Methanococcoides burtonii DSM_6242 | Archaea | Euryarchaeota | genome | 2273 | NCBI | https://www.ncbi.nlm.nih.gov/genbank/ |
| Methanococcus aeolicus Nankai_3 | Archaea | Euryarchaeota | genome | 1490 | NCBI | https://www.ncbi.nlm.nih.gov/genbank/ |
| Methanococcus maripaludis C5 | Archaea | Euryarchaeota | genome | 1840 | NCBI | https://www.ncbi.nlm.nih.gov/genbank/ |
| Methanococcus vannielii | Archaea | Euryarchaeota | genome | 1678 | NCBI | https://www.ncbi.nlm.nih.gov/genbank/ |
| Methanococcus voltae A3 | Archaea | Euryarchaeota | genome | 1690 | NCBI | https://www.ncbi.nlm.nih.gov/genbank/ |
| Methanocorpusculum labreanum Z | Archaea | Euryarchaeota | genome | 1739 | NCBI | https://www.ncbi.nlm.nih.gov/genbank/ |

| Species | Domain | Taxonomy | Asembly type | N. proteins | Assembly source | Source link |
|---|---|---|---|---|---|---|
| Methanoculleus marisnigri JR1 | Archaea | Euryarchaeota | genome | 2489 | NCBI | https://www.ncbi.nlm.nih.gov/genbank/ |
| Methanohalobium evestigatum Z-7303 | Archaea | Euryarchaeota | genome | 2254 | NCBI | https://www.ncbi.nlm.nih.gov/genbank/ |
| Methanohalophilus mahii DSM 5219 | Archaea | Euryarchaeota | genome | 1987 | NCBI | https://www.ncbi.nlm.nih.gov/genbank/ |
| Methanoplanus petrolearius DSM11571 | Archaea | Euryarchaeota | genome | 2785 | NCBI | https://www.ncbi.nlm.nih.gov/genbank/ |
| Methanopyrus kandleri AV19 | Archaea | Euryarchaeota | genome | 1687 | NCBI | https://www.ncbi.nlm.nih.gov/genbank/ |
| Methanosaeta concilii GP6 | Archaea | Euryarchaeota | genome | 2850 | NCBI | https://www.ncbi.nlm.nih.gov/genbank/ |
| Methanosaeta harundinacea 6Ac | Archaea | Euryarchaeota | genome | 2462 | NCBI | https://www.ncbi.nlm.nih.gov/genbank/ |
| Methanosaeta thermophila PT | Archaea | Euryarchaeota | genome | 1696 | NCBI | https://www.ncbi.nlm.nih.gov/genbank/ |
| Methanosalsum zhilinae DSM 4017 | Archaea | Euryarchaeota | genome | 1976 | NCBI | https://www.ncbi.nlm.nih.gov/genbank/ |
| Methanosarcina acetivorans C2A | Archaea | Euryarchaeota | genome | 4540 | NCBI | https://www.ncbi.nlm.nih.gov/genbank/ |
| Methanosarcina barkeri str. Fusaro | Archaea | Euryarchaeota | genome | 3624 | NCBI | https://www.ncbi.nlm.nih.gov/genbank/ |
| Methanosarcina mazei Go1 | Archaea | Euryarchaeota | genome | 3368 | NCBI | https://www.ncbi.nlm.nih.gov/genbank/ |
| Methanosphaera stadtmanae DSM_3091 | Archaea | Euryarchaeota | genome | 1534 | NCBI | https://www.ncbi.nlm.nih.gov/genbank/ |
| Methanosphaerula palustris E1-9c | Archaea | Euryarchaeota | genome | 2655 | NCBI | https://www.ncbi.nlm.nih.gov/genbank/ |
| Methanospirillum hungatei JF_1 | Archaea | Euryarchaeota | genome | 3139 | NCBI | https://www.ncbi.nlm.nih.gov/genbank/ |
| Methanothermobacter marburgensis str. Marburg | Archaea | Euryarchaeota | genome | 1757 | NCBI | https://www.ncbi.nlm.nih.gov/genbank/ |
| Methanothermobacter thermautotrophicus | Archaea | Euryarchaeota | genome | 1907 | NCBI | https://www.ncbi.nlm.nih.gov/genbank/ |
| Methanothermococcus okinawensis IH1 | Archaea | Euryarchaeota | genome | 1729 | NCBI | https://www.ncbi.nlm.nih.gov/genbank/ |
| Methanothermus fervidus | Archaea | Euryarchaeota | genome | 1283 | NCBI | https://www.ncbi.nlm.nih.gov/genbank/ |
| Methanotorris igneus Kol 5 | Archaea | Euryarchaeota | genome | 1772 | NCBI | https://www.ncbi.nlm.nih.gov/genbank/ |
| Nanoarchaeum equitans Kin4_M | Archaea | Euryarchaeota | genome | 536 | NCBI | https://www.ncbi.nlm.nih.gov/genbank/ |
| Natrialba magadii ATCC 43099 | Archaea | Euryarchaeota | genome | 4212 | NCBI | https://www.ncbi.nlm.nih.gov/genbank/ |
| Natronomonas pharaonis DSM_2160 | Archaea | Euryarchaeota | genome | 2822 | NCBI | https://www.ncbi.nlm.nih.gov/genbank/ |
| Picrophilus torridus DSM_9790 | Archaea | Euryarchaeota | genome | 1535 | NCBI | https://www.ncbi.nlm.nih.gov/genbank/ |
| Pyrococcus abyssi GE5 | Archaea | Euryarchaeota | genome | 1782 | NCBI | https://www.ncbi.nlm.nih.gov/genbank/ |
| Pyrococcus furiosus DSM 3638 | Archaea | Euryarchaeota | genome | 2125 | NCBI | https://www.ncbi.nlm.nih.gov/genbank/ |
| Pyrococcus horikoshii OT3 | Archaea | Euryarchaeota | genome | 1955 | NCBI | https://www.ncbi.nlm.nih.gov/genbank/ |
| Pyrococcus sp. NA2 | Archaea | Euryarchaeota | genome | 1980 | NCBI | https://www.ncbi.nlm.nih.gov/genbank/ |
| Pyrococcus yayanosii CH1 | Archaea | Euryarchaeota | genome | 1865 | NCBI | https://www.ncbi.nlm.nih.gov/genbank/ |
| Thermococcus barophilus MP | Archaea | Euryarchaeota | genome | 2207 | NCBI | https://www.ncbi.nlm.nih.gov/genbank/ |
| Thermococcus gammatolerans EJ3 | Archaea | Euryarchaeota | genome | 2156 | NCBI | https://www.ncbi.nlm.nih.gov/genbank/ |
| Thermococcus kodakarensis KOD1 | Archaea | Euryarchaeota | genome | 2306 | NCBI | https://www.ncbi.nlm.nih.gov/genbank/ |
| Thermococcus onnurineus NA1 | Archaea | Euryarchaeota | genome | 1976 | NCBI | https://www.ncbi.nlm.nih.gov/genbank/ |
| Thermococcus sibiricus MM 739 | Archaea | Euryarchaeota | genome | 2035 | NCBI | https://www.ncbi.nlm.nih.gov/genbank/ |
| Thermococcus sp. AM4 | Archaea | Euryarchaeota | genome | 2168 | NCBI | https://www.ncbi.nlm.nih.gov/genbank/ |
| Thermoplasma acidophilum DSM_1728 | Archaea | Euryarchaeota | genome | 1482 | NCBI | https://www.ncbi.nlm.nih.gov/genbank/ |
| Thermoplasma volcanium GSS1 | Archaea | Euryarchaeota | genome | 1499 | NCBI | https://www.ncbi.nlm.nih.gov/genbank/ |
| Uncultured Methanogenic archaeon RC-I | Archaea | Euryarchaeota | genome | 3085 | NCBI | https://www.ncbi.nlm.nih.gov/genbank/ |
| Korarchaeum cryptofilum OPF8 | Archaea | Korarchaeota | genome | 1602 | NCBI | https://www.ncbi.nlm.nih.gov/genbank/ |
| Caldiarchaeum subterraneum | Archaea | Thaumarchaeota | genome | 3235 | NCBI | https://www.ncbi.nlm.nih.gov/genbank/ |
| Candidatus Nitrososphaera gargensis Ga9.2 | Archaea | Thaumarchaeota | genome | 3565 | NCBI | https://www.ncbi.nlm.nih.gov/genbank/ |
| Candidatus Nitrosotenuis uzonensis | Archaea | Thaumarchaeota | genome | 1944 | NCBI | https://www.ncbi.nlm.nih.gov/genbank/ |
| Cenarchaeum symbiosum A | Archaea | Thaumarchaeota | genome | 2017 | NCBI | https://www.ncbi.nlm.nih.gov/genbank/ |
| Nitrosopumilus maritimus SCM1 | Archaea | Thaumarchaeota | genome | 1795 | NCBI | https://www.ncbi.nlm.nih.gov/genbank/ |
| Acidobacteria bacterium Ellin345 | Bacteria | Acidobacteria | genome | 4777 | NCBI | https://www.ncbi.nlm.nih.gov/genbank/ |
| Solibacter usitatus Ellin6076 | Bacteria | Acidobacteria | genome | 7826 | NCBI | https://www.ncbi.nlm.nih.gov/genbank/ |
| Acidothermus cellulolyticus 11B | Bacteria | Actinobacteria | genome | 2157 | NCBI | https://www.ncbi.nlm.nih.gov/genbank/ |
| Actinobacillus pleuropneumoniae L20 | Bacteria | Actinobacteria | genome | 2012 | NCBI | https://www.ncbi.nlm.nih.gov/genbank/ |
| Arthrobacter aurescens TC1 | Bacteria | Actinobacteria | genome | 4590 | NCBI | https://www.ncbi.nlm.nih.gov/genbank/ |
| Bifidobacterium adolescentis ATCC_15703 | Bacteria | Actinobacteria | genome | 1631 | NCBI | https://www.ncbi.nlm.nih.gov/genbank/ |
| Clavibacter michiganensis NCPPB_382 | Bacteria | Actinobacteria | genome | 3079 | NCBI | https://www.ncbi.nlm.nih.gov/genbank/ |
| Corynebacterium diphtheriae NCTC_13129 | Bacteria | Actinobacteria | genome | 2272 | NCBI | https://www.ncbi.nlm.nih.gov/genbank/ |
| Frankia alni ACN14a | Bacteria | Actinobacteria | genome | 6711 | NCBI | https://www.ncbi.nlm.nih.gov/genbank/ |
| Kineococcus radiotolerans SRS30216 | Bacteria | Actinobacteria | genome | 4681 | NCBI | https://www.ncbi.nlm.nih.gov/genbank/ |
| Kocuria rhizophila DC2201 | Bacteria | Actinobacteria | genome | 2357 | NCBI | https://www.ncbi.nlm.nih.gov/genbank/ |
| Leifsonia xyli CTCB07 | Bacteria | Actinobacteria | genome | 2030 | NCBI | https://www.ncbi.nlm.nih.gov/genbank/ |
| Mycobacterium abscessus | Bacteria | Actinobacteria | genome | 4941 | NCBI | https://www.ncbi.nlm.nih.gov/genbank/ |
| Nocardia farcinica IFM_10152 | Bacteria | Actinobacteria | genome | 5936 | NCBI | https://www.ncbi.nlm.nih.gov/genbank/ |
| Nocardioides sp. JS614 | Bacteria | Actinobacteria | genome | 4909 | NCBI | https://www.ncbi.nlm.nih.gov/genbank/ |
| Propionibacterium acnes KPA171202 | Bacteria | Actinobacteria | genome | 2297 | NCBI | https://www.ncbi.nlm.nih.gov/genbank/ |
| Renibacterium salmoninarum ATCC_33209 | Bacteria | Actinobacteria | genome | 3507 | NCBI | https://www.ncbi.nlm.nih.gov/genbank/ |
| Rhodococcus sp. RHA1 | Bacteria | Actinobacteria | genome | 9145 | NCBI | https://www.ncbi.nlm.nih.gov/genbank/ |
| Rubrobacter xylanophilus DSM_9941 | Bacteria | Actinobacteria | genome | 3140 | NCBI | https://www.ncbi.nlm.nih.gov/genbank/ |
| Saccharopolyspora erythraea NRRL_2338 | Bacteria | Actinobacteria | genome | 7197 | NCBI | https://www.ncbi.nlm.nih.gov/genbank/ |
| Salinispora arenicola CNS_205 | Bacteria | Actinobacteria | genome | 4917 | NCBI | https://www.ncbi.nlm.nih.gov/genbank/ |
| Streptococcus agalactiae 2603V_R | Bacteria | Actinobacteria | genome | 2124 | NCBI | https://www.ncbi.nlm.nih.gov/genbank/ |
| Streptomyces avermitilis MA_4680 | Bacteria | Actinobacteria | genome | 7676 | NCBI | https://www.ncbi.nlm.nih.gov/genbank/ |
| Thermobifida fusca YX | Bacteria | Actinobacteria | genome | 3110 | NCBI | https://www.ncbi.nlm.nih.gov/genbank/ |
| Tropheryma whipplei TW08_27 | Bacteria | Actinobacteria | genome | 783 | NCBI | https://www.ncbi.nlm.nih.gov/genbank/ |
| Aquifex aeolicus VF5 | Bacteria | Aquificales | genome | 1560 | NCBI | https://www.ncbi.nlm.nih.gov/genbank/ |
| Hydrogenobaculum sp. Y04AAS1 | Bacteria | Aquificales | genome | 1629 | NCBI | https://www.ncbi.nlm.nih.gov/genbank/ |
| Sulfurihydrogenibium sp. YO3AOP1 | Bacteria | Aquificales | genome | 1721 | NCBI | https://www.ncbi.nlm.nih.gov/genbank/ |
| Bacteroides fragilis NCTC_9343 | Bacteria | CFB | genome | 4231 | NCBI | https://www.ncbi.nlm.nih.gov/genbank/ |

| Species | Domain | Taxonomy | Asembly type | N. proteins | Assembly source | Source link |
|---|---|---|---|---|---|---|
| Bacteroides fragilis YCH46 | Bacteria | CFB | genome | 4625 | NCBI | https://www.ncbi.nlm.nih.gov/genbank/ |
| Cytophaga hutchinsonii ATCC_33406 | Bacteria | CFB | genome | 3785 | NCBI | https://www.ncbi.nlm.nih.gov/genbank/ |
| Flavobacterium johnsoniae UW101 | Bacteria | CFB | genome | 5017 | NCBI | https://www.ncbi.nlm.nih.gov/genbank/ |
| Flavobacterium psychrophilum JIP02_86 | Bacteria | CFB | genome | 2412 | NCBI | https://www.ncbi.nlm.nih.gov/genbank/ |
| Gramella forsetii KT0803 | Bacteria | CFB | genome | 3584 | NCBI | https://www.ncbi.nlm.nih.gov/genbank/ |
| Parabacteroides distasonis ATCC_8503 | Bacteria | CFB | genome | 3850 | NCBI | https://www.ncbi.nlm.nih.gov/genbank/ |
| Porphyromonas gingivalis W83 | Bacteria | CFB | genome | 1909 | NCBI | https://www.ncbi.nlm.nih.gov/genbank/ |
| Salinibacter ruber DSM_13855 | Bacteria | CFB | genome | 2833 | NCBI | https://www.ncbi.nlm.nih.gov/genbank/ |
| Amoebophilus asiaticus 5a2 | Bacteria | Chlorobi | genome | 1283 | NCBI | https://www.ncbi.nlm.nih.gov/genbank/ |
| Chlorobaculum parvum NCIB 8327 | Bacteria | Chlorobi | genome | 2043 | NCBI | https://www.ncbi.nlm.nih.gov/genbank/ |
| Chlorobium chlorochromatii CaD3 | Bacteria | Chlorobi | genome | 2002 | NCBI | https://www.ncbi.nlm.nih.gov/genbank/ |
| Chloroherpeton thalassium ATCC 35110 | Bacteria | Chlorobi | genome | 2710 | NCBI | https://www.ncbi.nlm.nih.gov/genbank/ |
| Pelodictyon luteolum DSM_273 | Bacteria | Chlorobi | genome | 2083 | NCBI | https://www.ncbi.nlm.nih.gov/genbank/ |
| Prosthecochloris vibrioformis DSM_265 | Bacteria | Chlorobi | genome | 1753 | NCBI | https://www.ncbi.nlm.nih.gov/genbank/ |
| Sulcia muelleri GWSS | Bacteria | Chlorobi | genome | 227 | NCBI | https://www.ncbi.nlm.nih.gov/genbank/ |
| Chloroflexus aurantiacus J_10_fl | Bacteria | Chloroflexi | genome | 3853 | NCBI | https://www.ncbi.nlm.nih.gov/genbank/ |
| Dehalococcoides sp. BAV1 | Bacteria | Chloroflexi | genome | 1371 | NCBI | https://www.ncbi.nlm.nih.gov/genbank/ |
| Herpetosiphon aurantiacus ATCC 23779 | Bacteria | Chloroflexi | genome | 5278 | NCBI | https://www.ncbi.nlm.nih.gov/genbank/ |
| Roseiflexus castenholzii DSM_13941 | Bacteria | Chloroflexi | genome | 4330 | NCBI | https://www.ncbi.nlm.nih.gov/genbank/ |
| Thermomicrobium roseum DSM 5159 | Bacteria | Chloroflexi | genome | 2854 | NCBI | https://www.ncbi.nlm.nih.gov/genbank/ |
| Acaryochloris marina MBIC11017 | Bacteria | Cyanobacteria | genome | 8383 | NCBI | https://www.ncbi.nlm.nih.gov/genbank/ |
| Acaryochloris sp. CCMEE 5410 | Bacteria | Cyanobacteria | genome | 6776 | NCBI | https://www.ncbi.nlm.nih.gov/genbank/ |
| Anabaena azollae 0708 | Bacteria | Cyanobacteria | genome | 3646 | NCBI | https://www.ncbi.nlm.nih.gov/genbank/ |
| Anabaena cylindrica PCC 7122 | Bacteria | Cyanobacteria | genome | 6129 | JGI | https://mycocosm.jgi.doe.gov/ |
| Anabaena sp. PCC 7108 | Bacteria | Cyanobacteria | genome | 5148 | JGI | https://mycocosm.jgi.doe.gov/ |
| Arthrospira maxima CS-328 | Bacteria | Cyanobacteria | genome | 5431 | NCBI | https://www.ncbi.nlm.nih.gov/genbank/ |
| Arthrospira platensis NIES-39 | Bacteria | Cyanobacteria | genome | 5967 | NCBI | https://www.ncbi.nlm.nih.gov/genbank/ |
| Arthrospira platensis str. Paraca | Bacteria | Cyanobacteria | genome | 4630 | NCBI | https://www.ncbi.nlm.nih.gov/genbank/ |
| Calothrix sp. PCC 6303 | Bacteria | Cyanobacteria | genome | 5719 | JGI | https://mycocosm.jgi.doe.gov/ |
| Calothrix sp. PCC 7103 | Bacteria | Cyanobacteria | genome | 10042 | JGI | https://mycocosm.jgi.doe.gov/ |
| Calothrix sp. PCC 7507 | Bacteria | Cyanobacteria | genome | 6132 | JGI | https://mycocosm.jgi.doe.gov/ |
| Candidatus Synechococcus calcipolaris | Bacteria | Cyanobacteria | genome | 3257 | NCBI | https://www.ncbi.nlm.nih.gov/genbank/ |
| Chamaesiphon minutus PCC 6605 | Bacteria | Cyanobacteria | genome | 5813 | NCBI | https://www.ncbi.nlm.nih.gov/genbank/ |
| Chlorogloeopsis fritschii PCC 6912 | Bacteria | Cyanobacteria | genome | 6491 | NCBI | https://www.ncbi.nlm.nih.gov/genbank/ |
| Chlorogloeopsis fritschii PCC 9212 | Bacteria | Cyanobacteria | genome | 6351 | NCBI | https://www.ncbi.nlm.nih.gov/genbank/ |
| Chroococcidiopsis sp. PCC 6712 | Bacteria | Cyanobacteria | genome | 5031 | JGI | https://mycocosm.jgi.doe.gov/ |
| Chroococcidiopsis thermalis PCC 7203 | Bacteria | Cyanobacteria | genome | 5961 | JGI | https://mycocosm.jgi.doe.gov/ |
| Coleofasciculus chthonoplastes PCC 7420 | Bacteria | Cyanobacteria | genome | 8187 | NCBI | https://www.ncbi.nlm.nih.gov/genbank/ |
| Crinalium epipsammum PCC 9333 | Bacteria | Cyanobacteria | genome | 4935 | JGI | https://mycocosm.jgi.doe.gov/ |
| Crocosphaera chwakensis CCY0110 | Bacteria | Cyanobacteria | genome | 6475 | NCBI | https://www.ncbi.nlm.nih.gov/genbank/ |
| Crocosphaera subtropica ATCC 51142 | Bacteria | Cyanobacteria | genome | 5304 | NCBI | https://www.ncbi.nlm.nih.gov/genbank/ |
| Crocosphaera subtropica ATCC 51472 | Bacteria | Cyanobacteria | genome | 5048 | NCBI | https://www.ncbi.nlm.nih.gov/genbank/ |
| Crocosphaera watsonii WH 0003 | Bacteria | Cyanobacteria | genome | 6120 | NCBI | https://www.ncbi.nlm.nih.gov/genbank/ |
| Crocosphaera watsonii WH 8501 | Bacteria | Cyanobacteria | genome | 5958 | NCBI | https://www.ncbi.nlm.nih.gov/genbank/ |
| Cyanobacterium aponinum PCC 10605 | Bacteria | Cyanobacteria | genome | 3540 | JGI | https://mycocosm.jgi.doe.gov/ |
| cyanobacterium sp. PCC 7702 | Bacteria | Cyanobacteria | genome | 4375 | NCBI | https://www.ncbi.nlm.nih.gov/genbank/ |
| Cyanobacterium stanieri PCC 7202 | Bacteria | Cyanobacteria | genome | 2880 | JGI | https://mycocosm.jgi.doe.gov/ |
| Cyanothece sp. PCC 7424 | Bacteria | Cyanobacteria | genome | 5653 | NCBI | https://www.ncbi.nlm.nih.gov/genbank/ |
| Cyanothece sp. PCC 7425 | Bacteria | Cyanobacteria | genome | 5232 | NCBI | https://www.ncbi.nlm.nih.gov/genbank/ |
| Cylindrospermopsis raciborskii CS-505 | Bacteria | Cyanobacteria | genome | 3406 | NCBI | https://www.ncbi.nlm.nih.gov/genbank/ |
| Cylindrospermum stagnale PCC 7417 | Bacteria | Cyanobacteria | genome | 6595 | JGI | https://mycocosm.jgi.doe.gov/ |
| Dactylococcopsis salina PCC 8305 | Bacteria | Cyanobacteria | genome | 3557 | JGI | https://mycocosm.jgi.doe.gov/ |
| Fischerella muscicola PCC 7414 | Bacteria | Cyanobacteria | genome | 5462 | NCBI | https://www.ncbi.nlm.nih.gov/genbank/ |
| Fischerella muscicola SAG 1427-1 | Bacteria | Cyanobacteria | genome | 5798 | NCBI | https://www.ncbi.nlm.nih.gov/genbank/ |
| Fischerella sp. JSC-11 | Bacteria | Cyanobacteria | genome | 4553 | NCBI | https://www.ncbi.nlm.nih.gov/genbank/ |
| Fischerella sp. PCC 9339 | Bacteria | Cyanobacteria | genome | 6563 | NCBI | https://www.ncbi.nlm.nih.gov/genbank/ |
| Fischerella sp. PCC 9431 | Bacteria | Cyanobacteria | genome | 6071 | JGI | https://mycocosm.jgi.doe.gov/ |
| Fischerella sp. PCC 9605 | Bacteria | Cyanobacteria | genome | 6994 | JGI | https://mycocosm.jgi.doe.gov/ |
| Fischerella thermalis PCC 7521 | Bacteria | Cyanobacteria | genome | 4342 | NCBI | https://www.ncbi.nlm.nih.gov/genbank/ |
| Fortiea contorta PCC 7126 | Bacteria | Cyanobacteria | genome | 5130 | JGI | https://mycocosm.jgi.doe.gov/ |
| Geminocystis herdmanii PCC 6308 | Bacteria | Cyanobacteria | genome | 4114 | JGI | https://mycocosm.jgi.doe.gov/ |
| Gloeocapsa sp. PCC 73106 | Bacteria | Cyanobacteria | genome | 4031 | JGI | https://mycocosm.jgi.doe.gov/ |
| Gloeocapsa sp. PCC 7428 | Bacteria | Cyanobacteria | genome | 5241 | JGI | https://mycocosm.jgi.doe.gov/ |
| Gloeothece verrucosa PCC 7822 | Bacteria | Cyanobacteria | genome | 6557 | NCBI | https://www.ncbi.nlm.nih.gov/genbank/ |
| Halothece sp. PCC 7418 | Bacteria | Cyanobacteria | genome | 3849 | JGI | https://mycocosm.jgi.doe.gov/ |
| Kamptonema formosum PCC 6407 | Bacteria | Cyanobacteria | genome | 5642 | JGI | https://mycocosm.jgi.doe.gov/ |
| Kamptonema sp. PCC 6506 | Bacteria | Cyanobacteria | genome | 5826 | NCBI | https://www.ncbi.nlm.nih.gov/genbank/ |
| Leptolyngbya sp. PCC 7376 | Bacteria | Cyanobacteria | genome | 4510 | JGI | https://mycocosm.jgi.doe.gov/ |
| Limnospira indica PCC 8005 | Bacteria | Cyanobacteria | genome | 5653 | NCBI | https://www.ncbi.nlm.nih.gov/genbank/ |
| Lyngbya sp. PCC 8106 | Bacteria | Cyanobacteria | genome | 6142 | NCBI | https://www.ncbi.nlm.nih.gov/genbank/ |
| Mastigocladopsis repens PCC 10914 | Bacteria | Cyanobacteria | genome | 5761 | JGI | https://mycocosm.jgi.doe.gov/ |
| Microcoleus sp. PCC 7113 | Bacteria | Cyanobacteria | genome | 6658 | JGI | https://mycocosm.jgi.doe.gov/ |
| Microcoleus vaginatus FGP-2 | Bacteria | Cyanobacteria | genome | 5102 | NCBI | https://www.ncbi.nlm.nih.gov/genbank/ |

| Species | Domain | Taxonomy | Asembly type | N. proteins | Assembly source | Source link |
|---|---|---|---|---|---|---|
| Microcystis aeruginosa NIES-843 | Bacteria | Cyanobacteria | genome | 6312 | NCBI | https://www.ncbi.nlm.nih.gov/genbank/ |
| Microcystis aeruginosa PCC 7806 | Bacteria | Cyanobacteria | genome | 4891 | NCBI | https://www.ncbi.nlm.nih.gov/genbank/ |
| Moorena producens 3L | Bacteria | Cyanobacteria | genome | 7355 | NCBI | https://www.ncbi.nlm.nih.gov/genbank/ |
| Myxosarcina sp. GI1 | Bacteria | Cyanobacteria | genome | 6110 | NCBI | https://www.ncbi.nlm.nih.gov/genbank/ |
| Nodularia spumigena CCY9414 | Bacteria | Cyanobacteria | genome | 4860 | NCBI | https://www.ncbi.nlm.nih.gov/genbank/ |
| Nostoc punctiforme 11 | Bacteria | Cyanobacteria | genome | 6690 | NCBI | https://www.ncbi.nlm.nih.gov/genbank/ |
| Nostoc sp. PCC 7107 | Bacteria | Cyanobacteria | genome | 5372 | JGI | https://mycocosm.jgi.doe.gov/ |
| Nostoc sp. PCC 7524 | Bacteria | Cyanobacteria | genome | 5534 | JGI | https://mycocosm.jgi.doe.gov/ |
| Nostoc sp. PCC7120 | Bacteria | Cyanobacteria | genome | 6130 | NCBI | https://www.ncbi.nlm.nih.gov/genbank/ |
| Oscillatoria acuminata PCC 6304 | Bacteria | Cyanobacteria | genome | 5953 | JGI | https://mycocosm.jgi.doe.gov/ |
| Oscillatoria nigro-viridis PCC 7112 | Bacteria | Cyanobacteria | genome | 6805 | JGI | https://mycocosm.jgi.doe.gov/ |
| Oscillatoria sp. PCC 10802 | Bacteria | Cyanobacteria | genome | 6950 | JGI | https://mycocosm.jgi.doe.gov/ |
| Pleurocapsa sp. PCC 7319 | Bacteria | Cyanobacteria | genome | 6566 | JGI | https://mycocosm.jgi.doe.gov/ |
| Pleurocapsa sp. PCC 7327 | Bacteria | Cyanobacteria | genome | 4573 | JGI | https://mycocosm.jgi.doe.gov/ |
| Raphidiopsis brookii D9 | Bacteria | Cyanobacteria | genome | 3007 | NCBI | https://www.ncbi.nlm.nih.gov/genbank/ |
| Richelia intracellularis HH01 | Bacteria | Cyanobacteria | genome | 3841 | NCBI | https://www.ncbi.nlm.nih.gov/genbank/ |
| Rippkaea orientalis PCC 8801 | Bacteria | Cyanobacteria | genome | 4349 | NCBI | https://www.ncbi.nlm.nih.gov/genbank/ |
| Rippkaea orientalis PCC 8802 | Bacteria | Cyanobacteria | genome | 4403 | NCBI | https://www.ncbi.nlm.nih.gov/genbank/ |
| Rivularia sp. PCC 7116 | Bacteria | Cyanobacteria | genome | 6848 | JGI | https://mycocosm.jgi.doe.gov/ |
| Rubidibacter lacunae KORDI 51-2 | Bacteria | Cyanobacteria | genome | 3450 | NCBI | https://www.ncbi.nlm.nih.gov/genbank/ |
| Scytonema hofmanni PCC 7110 | Bacteria | Cyanobacteria | genome | 9591 | NCBI | https://www.ncbi.nlm.nih.gov/genbank/ |
| Spirulina major PCC 6313 | Bacteria | Cyanobacteria | genome | 4356 | JGI | https://mycocosm.jgi.doe.gov/ |
| Spirulina subsalsa PCC 9445 | Bacteria | Cyanobacteria | genome | 4565 | JGI | https://mycocosm.jgi.doe.gov/ |
| Stanieria cyanosphaera PCC 7437 | Bacteria | Cyanobacteria | genome | 4955 | JGI | https://mycocosm.jgi.doe.gov/ |
| Synechococcus sp. BDU 130192 | Bacteria | Cyanobacteria | genome | 3014 | NCBI | https://www.ncbi.nlm.nih.gov/genbank/ |
| Synechococcus sp. NIES-970 | Bacteria | Cyanobacteria | genome | 2844 | NCBI | https://www.ncbi.nlm.nih.gov/genbank/ |
| Synechococcus sp. NKBG042902 | Bacteria | Cyanobacteria | genome | 3066 | NCBI | https://www.ncbi.nlm.nih.gov/genbank/ |
| Synechococcus sp. NKBG15041c | Bacteria | Cyanobacteria | genome | 2865 | NCBI | https://www.ncbi.nlm.nih.gov/genbank/ |
| Synechococcus sp. PCC 11901 | Bacteria | Cyanobacteria | genome | 3162 | NCBI | https://www.ncbi.nlm.nih.gov/genbank/ |
| Synechococcus sp. PCC 6312 | Bacteria | Cyanobacteria | genome | 3740 | JGI | https://mycocosm.jgi.doe.gov/ |
| Synechococcus sp. PCC 7002 | Bacteria | Cyanobacteria | genome | 3252 | NCBI | https://www.ncbi.nlm.nih.gov/genbank/ |
| Synechococcus sp. PCC 7117 | Bacteria | Cyanobacteria | genome | 3152 | NCBI | https://www.ncbi.nlm.nih.gov/genbank/ |
| Synechococcus sp. PCC 73109 | Bacteria | Cyanobacteria | genome | 3012 | NCBI | https://www.ncbi.nlm.nih.gov/genbank/ |
| Synechococcus sp. PCC 8807 | Bacteria | Cyanobacteria | genome | 3072 | NCBI | https://www.ncbi.nlm.nih.gov/genbank/ |
| Synechocystis sp. PCC 6803 | Bacteria | Cyanobacteria | genome | 3572 | NCBI | https://www.ncbi.nlm.nih.gov/genbank/ |
| Synechocystis sp. PCC 7509 | Bacteria | Cyanobacteria | genome | 4787 | JGI | https://mycocosm.jgi.doe.gov/ |
| Thermosynechococcus elongatus BP-1 | Bacteria | Cyanobacteria | genome | 2476 | NCBI | https://www.ncbi.nlm.nih.gov/genbank/ |
| Tolypothrix sp. PCC 9009 | Bacteria | Cyanobacteria | genome | 7225 | JGI | https://mycocosm.jgi.doe.gov/ |
| Trichodesmium erythraeum | Bacteria | Cyanobacteria | genome | 4451 | NCBI | https://www.ncbi.nlm.nih.gov/genbank/ |
| Trichormus variabilis | Bacteria | Cyanobacteria | genome | 5661 | NCBI | https://www.ncbi.nlm.nih.gov/genbank/ |
| Xenococcus sp. PCC 7305 | Bacteria | Cyanobacteria | genome | 5329 | JGI | https://mycocosm.jgi.doe.gov/ |
| Gloeobacter kilaueensis JS1 | Bacteria | Cyanobacteria | genome | 4462 | NCBI | https://www.ncbi.nlm.nih.gov/genbank/ |
| Gloeobacter violaceus PCC 7421 | Bacteria | Cyanobacteria | genome | 4430 | NCBI | https://www.ncbi.nlm.nih.gov/genbank/ |
| Geitlerinema sp. PCC 7407 | Bacteria | Cyanobacteria | genome | 3812 | NCBI | https://www.ncbi.nlm.nih.gov/genbank/ |
| Leptolyngbya boryana PCC 6306 | Bacteria | Cyanobacteria | genome | 6778 | JGI | https://mycocosm.jgi.doe.gov/ |
| Leptolyngbya sp. PCC 6406 | Bacteria | Cyanobacteria | genome | 5124 | JGI | https://mycocosm.jgi.doe.gov/ |
| Leptolyngbya sp. PCC 7375 | Bacteria | Cyanobacteria | genome | 8065 | JGI | https://mycocosm.jgi.doe.gov/ |
| Nodosilinea nodulosa PCC 7104 | Bacteria | Cyanobacteria | genome | 6278 | JGI | https://mycocosm.jgi.doe.gov/ |
| Synechococcus elongatus PCC 6301 | Bacteria | Cyanobacteria | genome | 2527 | NCBI | https://www.ncbi.nlm.nih.gov/genbank/ |
| Synechococcus elongatus PCC 7942 | Bacteria | Cyanobacteria | genome | 2728 | NCBI | https://www.ncbi.nlm.nih.gov/genbank/ |
| Synechococcus sp. PCC 7335 | Bacteria | Cyanobacteria | genome | 5469 | NCBI | https://www.ncbi.nlm.nih.gov/genbank/ |
| Cyanobium gracile PCC 6307 | Bacteria | Cyanobacteria | genome | 3360 | JGI | https://mycocosm.jgi.doe.gov/ |
| Prochlorococcus marinus str. AS9601 | Bacteria | Cyanobacteria | genome | 1921 | NCBI | https://www.ncbi.nlm.nih.gov/genbank/ |
| Prochlorococcus marinus str. CCMP1375 | Bacteria | Cyanobacteria | genome | 1883 | NCBI | https://www.ncbi.nlm.nih.gov/genbank/ |
| Prochlorococcus marinus str. MIT 9215 | Bacteria | Cyanobacteria | genome | 1983 | NCBI | https://www.ncbi.nlm.nih.gov/genbank/ |
| Prochlorococcus marinus str. MIT 9301 | Bacteria | Cyanobacteria | genome | 1907 | NCBI | https://www.ncbi.nlm.nih.gov/genbank/ |
| Prochlorococcus marinus str. MIT 9303 | Bacteria | Cyanobacteria | genome | 2997 | NCBI | https://www.ncbi.nlm.nih.gov/genbank/ |
| Prochlorococcus marinus str. MIT 9312 | Bacteria | Cyanobacteria | genome | 1810 | NCBI | https://www.ncbi.nlm.nih.gov/genbank/ |
| Prochlorococcus marinus str. MIT 9313 | Bacteria | Cyanobacteria | genome | 2269 | NCBI | https://www.ncbi.nlm.nih.gov/genbank/ |
| Prochlorococcus marinus str. MIT 9515 | Bacteria | Cyanobacteria | genome | 1906 | NCBI | https://www.ncbi.nlm.nih.gov/genbank/ |
| Prochlorococcus marinus str. NATL1A | Bacteria | Cyanobacteria | genome | 2193 | NCBI | https://www.ncbi.nlm.nih.gov/genbank/ |
| Prochlorococcus marinus str. NATL2A | Bacteria | Cyanobacteria | genome | 2163 | NCBI | https://www.ncbi.nlm.nih.gov/genbank/ |
| Prochlorococcus marinus subsp. pastoris str. CCMP1986 | Bacteria | Cyanobacteria | genome | 1717 | NCBI | https://www.ncbi.nlm.nih.gov/genbank/ |
| Synechococcus sp. CC9311 | Bacteria | Cyanobacteria | genome | 2892 | NCBI | https://www.ncbi.nlm.nih.gov/genbank/ |
| Synechococcus sp. CC9605 | Bacteria | Cyanobacteria | genome | 2645 | NCBI | https://www.ncbi.nlm.nih.gov/genbank/ |
| Synechococcus sp. CC9902 | Bacteria | Cyanobacteria | genome | 2307 | NCBI | https://www.ncbi.nlm.nih.gov/genbank/ |
| Synechococcus sp. RCC307 | Bacteria | Cyanobacteria | genome | 2535 | NCBI | https://www.ncbi.nlm.nih.gov/genbank/ |
| Synechococcus sp. WH 5701 | Bacteria | Cyanobacteria | genome | 3346 | NCBI | https://www.ncbi.nlm.nih.gov/genbank/ |
| Synechococcus sp. WH 7803 | Bacteria | Cyanobacteria | genome | 2533 | NCBI | https://www.ncbi.nlm.nih.gov/genbank/ |
| Synechococcus sp. WH 8102 | Bacteria | Cyanobacteria | genome | 2519 | NCBI | https://www.ncbi.nlm.nih.gov/genbank/ |
| Pseudanabaena biceps PCC 7429 | Bacteria | Cyanobacteria | genome | 4727 | JGI | https://mycocosm.jgi.doe.gov/ |
| Pseudanabaena sp. PCC 6802 | Bacteria | Cyanobacteria | genome | 5212 | JGI | https://mycocosm.jgi.doe.gov/ |
| Pseudanabaena sp. PCC 7367 | Bacteria | Cyanobacteria | genome | 3908 | JGI | https://mycocosm.jgi.doe.gov/ |

| Species | Domain | Taxonomy | Asembly type | N. proteins | Assembly source | Source link |
|---|---|---|---|---|---|---|
| Synechococcus sp. PCC 7502 | Bacteria | Cyanobacteria | genome | 3574 | JGI | https://mycocosm.jgi.doe.gov/ |
| Synechococcus sp. JA-2-3Ba | Bacteria | Cyanobacteria | genome | 2862 | NCBI | https://www.ncbi.nlm.nih.gov/genbank/ |
| Synechococcus sp. JA-3-3Ab | Bacteria | Cyanobacteria | genome | 2760 | NCBI | https://www.ncbi.nlm.nih.gov/genbank/ |
| Synechococcus sp. PCC 7336 | Bacteria | Cyanobacteria | genome | 4682 | JGI | https://mycocosm.jgi.doe.gov/ |
| Deinococcus geothermalis DSM_11300 | Bacteria | Deinococcales | genome | 3054 | NCBI | https://www.ncbi.nlm.nih.gov/genbank/ |
| Thermus thermophilus HB8 | Bacteria | Deinococcales | genome | 2238 | NCBI | https://www.ncbi.nlm.nih.gov/genbank/ |
| Dictyoglomus thermophilum H-6-12 | Bacteria | Dictyoglomi | genome | 1912 | NCBI | https://www.ncbi.nlm.nih.gov/genbank/ |
| Acholeplasma laidlawii PG_8A | Bacteria | Firmicutes | genome | 1380 | NCBI | https://www.ncbi.nlm.nih.gov/genbank/ |
| Alkaliphilus metalliredigens QYMF | Bacteria | Firmicutes | genome | 4625 | NCBI | https://www.ncbi.nlm.nih.gov/genbank/ |
| Anaerocellum thermophilum DSM 6725 | Bacteria | Firmicutes | genome | 2666 | NCBI | https://www.ncbi.nlm.nih.gov/genbank/ |
| Anoxybacillus flavithermus WK1 | Bacteria | Firmicutes | genome | 2832 | NCBI | https://www.ncbi.nlm.nih.gov/genbank/ |
| Bacillus amyloliquefaciens FZB42 | Bacteria | Firmicutes | genome | 3693 | NCBI | https://www.ncbi.nlm.nih.gov/genbank/ |
| Caldicellulosiruptor saccharolyticus DSM_8903 | Bacteria | Firmicutes | genome | 2679 | NCBI | https://www.ncbi.nlm.nih.gov/genbank/ |
| Candidatus Phytoplasma asteris AYWB | Bacteria | Firmicutes | genome | 693 | NCBI | https://www.ncbi.nlm.nih.gov/genbank/ |
| Carboxydothermus hydrogenoformans Z_2901 | Bacteria | Firmicutes | genome | 2620 | NCBI | https://www.ncbi.nlm.nih.gov/genbank/ |
| Clostridium acetobutylicum ATCC_824 | Bacteria | Firmicutes | genome | 3848 | NCBI | https://www.ncbi.nlm.nih.gov/genbank/ |
| Coprothermobacter proteolyticus DSM 5265 | Bacteria | Firmicutes | genome | 1482 | NCBI | https://www.ncbi.nlm.nih.gov/genbank/ |
| Desulfitobacterium hafniense Y51 | Bacteria | Firmicutes | genome | 5060 | NCBI | https://www.ncbi.nlm.nih.gov/genbank/ |
| Desulfotomaculum reducens MI_1 | Bacteria | Firmicutes | genome | 3276 | NCBI | https://www.ncbi.nlm.nih.gov/genbank/ |
| Enterococcus faecalis V583 | Bacteria | Firmicutes | genome | 3265 | NCBI | https://www.ncbi.nlm.nih.gov/genbank/ |
| Exiguobacterium sibiricum 255-15 | Bacteria | Firmicutes | genome | 3015 | NCBI | https://www.ncbi.nlm.nih.gov/genbank/ |
| Finegoldia magna ATCC_29328 | Bacteria | Firmicutes | genome | 1813 | NCBI | https://www.ncbi.nlm.nih.gov/genbank/ |
| Fusobacterium nucleatum ATCC_25586 | Bacteria | Firmicutes | genome | 2067 | NCBI | https://www.ncbi.nlm.nih.gov/genbank/ |
| Geobacillus kaustophilus HTA426 | Bacteria | Firmicutes | genome | 3540 | NCBI | https://www.ncbi.nlm.nih.gov/genbank/ |
| Halothermothrix orenii H 168 | Bacteria | Firmicutes | genome | 2342 | NCBI | https://www.ncbi.nlm.nih.gov/genbank/ |
| Heliobacterium modesticaldum Ice1 | Bacteria | Firmicutes | genome | 3000 | NCBI | https://www.ncbi.nlm.nih.gov/genbank/ |
| Lactobacillus acidophilus NCFM | Bacteria | Firmicutes | genome | 1862 | NCBI | https://www.ncbi.nlm.nih.gov/genbank/ |
| Lactococcus lactis Il1403 | Bacteria | Firmicutes | genome | 2321 | NCBI | https://www.ncbi.nlm.nih.gov/genbank/ |
| Leuconostoc citreum KM20 | Bacteria | Firmicutes | genome | 1820 | NCBI | https://www.ncbi.nlm.nih.gov/genbank/ |
| Listeria innocua Clip11262 | Bacteria | Firmicutes | genome | 3043 | NCBI | https://www.ncbi.nlm.nih.gov/genbank/ |
| Lysinibacillus sphaericus C3_41 | Bacteria | Firmicutes | genome | 4771 | NCBI | https://www.ncbi.nlm.nih.gov/genbank/ |
| Macrococcus caseolyticus JCSC5402 | Bacteria | Firmicutes | genome | 2052 | NCBI | https://www.ncbi.nlm.nih.gov/genbank/ |
| Mesoplasma florum L1 | Bacteria | Firmicutes | genome | 682 | NCBI | https://www.ncbi.nlm.nih.gov/genbank/ |
| Moorella thermoacetica ATCC 39073 | Bacteria | Firmicutes | genome | 2463 | NCBI | https://www.ncbi.nlm.nih.gov/genbank/ |
| Mycoplasma agalactiae PG2 | Bacteria | Firmicutes | genome | 742 | NCBI | https://www.ncbi.nlm.nih.gov/genbank/ |
| Natranaerobius thermophilus JWNM-WN-LF | Bacteria | Firmicutes | genome | 2906 | NCBI | https://www.ncbi.nlm.nih.gov/genbank/ |
| Oceanobacillus iheyensis HTE831 | Bacteria | Firmicutes | genome | 3500 | NCBI | https://www.ncbi.nlm.nih.gov/genbank/ |
| Oenococcus oeni PSU_1 | Bacteria | Firmicutes | genome | 1691 | NCBI | https://www.ncbi.nlm.nih.gov/genbank/ |
| Onion yellows Phytoplasma sp. OY-M | Bacteria | Firmicutes | genome | 754 | NCBI | https://www.ncbi.nlm.nih.gov/genbank/ |
| Pediococcus pentosaceus ATCC_25745 | Bacteria | Firmicutes | genome | 1755 | NCBI | https://www.ncbi.nlm.nih.gov/genbank/ |
| Pelotomaculum thermopropionicum SI | Bacteria | Firmicutes | genome | 2918 | NCBI | https://www.ncbi.nlm.nih.gov/genbank/ |
| Phytoplasma australiense | Bacteria | Firmicutes | genome | 696 | NCBI | https://www.ncbi.nlm.nih.gov/genbank/ |
| Staphylococcus aureus MRSA252 | Bacteria | Firmicutes | genome | 2656 | NCBI | https://www.ncbi.nlm.nih.gov/genbank/ |
| Symbiobacterium thermophilum IAM_14863 | Bacteria | Firmicutes | genome | 3338 | NCBI | https://www.ncbi.nlm.nih.gov/genbank/ |
| Syntrophomonas wolfei subsp. Wolfei str. Goettingen | Bacteria | Firmicutes | genome | 2504 | NCBI | https://www.ncbi.nlm.nih.gov/genbank/ |
| Thermoanaerobacter pseudethanolicus ATCC 33223 | Bacteria | Firmicutes | genome | 2243 | NCBI | https://www.ncbi.nlm.nih.gov/genbank/ |
| Ureaplasma parvum ATCC 700970 | Bacteria | Firmicutes | genome | 614 | NCBI | https://www.ncbi.nlm.nih.gov/genbank/ |
| Anaeromyxobacter sp. Fw109_5 | Bacteria | Myxobacteria | genome | 4466 | NCBI | https://www.ncbi.nlm.nih.gov/genbank/ |
| Haliangium ochraceum DSM 14365 | Bacteria | Myxobacteria | genome | 6719 | NCBI | https://www.ncbi.nlm.nih.gov/genbank/ |
| Myxococcus xanthus DK_1622 | Bacteria | Myxobacteria | genome | 7331 | NCBI | https://www.ncbi.nlm.nih.gov/genbank/ |
| Plesiocystis pacifica SIR-1 | Bacteria | Myxobacteria | genome | 8450 | NCBI | https://www.ncbi.nlm.nih.gov/genbank/ |
| Sorangium cellulosum | Bacteria | Myxobacteria | genome | 9381 | NCBI | https://www.ncbi.nlm.nih.gov/genbank/ |
| Thermodesulfovibrio yellowstonii DSM 11347 | Bacteria | Nitrospirae | genome | 2033 | NCBI | https://www.ncbi.nlm.nih.gov/genbank/ |
| Elusimicrobium minutum Pei191 | Bacteria | Elusimicrobiota | genome | 1529 | NCBI | https://www.ncbi.nlm.nih.gov/genbank/ |
| Acidithiobacillus ferrooxidans ATCC 2327 | Bacteria | Proteobacteria | genome | 3147 | NCBI | https://www.ncbi.nlm.nih.gov/genbank/ |
| Acinetobacter baumannii AB0057 | Bacteria | Proteobacteria | genome | 3801 | NCBI | https://www.ncbi.nlm.nih.gov/genbank/ |
| Aeromonas hydrophila ATCC_7966 | Bacteria | Proteobacteria | genome | 4122 | NCBI | https://www.ncbi.nlm.nih.gov/genbank/ |
| Alcanivorax borkumensis SK2 | Bacteria | Proteobacteria | genome | 2755 | NCBI | https://www.ncbi.nlm.nih.gov/genbank/ |
| Aliivibrio salmonicida LFI1238 | Bacteria | Proteobacteria | genome | 3911 | NCBI | https://www.ncbi.nlm.nih.gov/genbank/ |
| Alkalilimnicola ehrlichei MLHE_1 | Bacteria | Proteobacteria | genome | 2865 | NCBI | https://www.ncbi.nlm.nih.gov/genbank/ |
| Alteromonas macleodii Deep ecotype | Bacteria | Proteobacteria | genome | 4072 | NCBI | https://www.ncbi.nlm.nih.gov/genbank/ |
| Baumannia cicadellinicola Hc | Bacteria | Proteobacteria | genome | 595 | NCBI | https://www.ncbi.nlm.nih.gov/genbank/ |
| Blochmannia floridanus | Bacteria | Proteobacteria | genome | 583 | NCBI | https://www.ncbi.nlm.nih.gov/genbank/ |
| Buchnera aphidicola APS | Bacteria | Proteobacteria | genome | 555 | NCBI | https://www.ncbi.nlm.nih.gov/genbank/ |
| Carsonella ruddii PV | Bacteria | Proteobacteria | genome | 182 | NCBI | https://www.ncbi.nlm.nih.gov/genbank/ |
| Cellvibrio japonicus Ueda107 | Bacteria | Proteobacteria | genome | 3754 | NCBI | https://www.ncbi.nlm.nih.gov/genbank/ |
| Chromohalobacter salexigens DSM 3043 | Bacteria | Proteobacteria | genome | 3298 | NCBI | https://www.ncbi.nlm.nih.gov/genbank/ |
| Citrobacter koseri ATCC_BAA895 | Bacteria | Proteobacteria | genome | 5008 | NCBI | https://www.ncbi.nlm.nih.gov/genbank/ |
| Colwellia psychrerythraea 34H | Bacteria | Proteobacteria | genome | 4910 | NCBI | https://www.ncbi.nlm.nih.gov/genbank/ |
| Coxiella burnetii RSA_493 | Bacteria | Proteobacteria | genome | 1848 | NCBI | https://www.ncbi.nlm.nih.gov/genbank/ |
| Cronobacter sakazakii ATCC BAA-894 | Bacteria | Proteobacteria | genome | 4420 | NCBI | https://www.ncbi.nlm.nih.gov/genbank/ |
| Dichelobacter nodosus VCS1703A | Bacteria | Proteobacteria | genome | 1280 | NCBI | https://www.ncbi.nlm.nih.gov/genbank/ |
| Enterobacter sp. 638 | Bacteria | Proteobacteria | genome | 4240 | NCBI | https://www.ncbi.nlm.nih.gov/genbank/ |

| Species | Domain | Taxonomy | Asembly type | N. proteins | Assembly source | Source link |
|---|---|---|---|---|---|---|
| Escherichia coli 536 | Bacteria | Proteobacteria | genome | 4620 | NCBI | https://www.ncbi.nlm.nih.gov/genbank/ |
| Francisella novicida U112 | Bacteria | Proteobacteria | genome | 1719 | NCBI | https://www.ncbi.nlm.nih.gov/genbank/ |
| Haemophilus ducreyi 35000HP | Bacteria | Proteobacteria | genome | 1717 | NCBI | https://www.ncbi.nlm.nih.gov/genbank/ |
| Hahella chejuensis KCTC_2396 | Bacteria | Proteobacteria | genome | 6778 | NCBI | https://www.ncbi.nlm.nih.gov/genbank/ |
| Halorhodospira halophila SL1 | Bacteria | Proteobacteria | genome | 2407 | NCBI | https://www.ncbi.nlm.nih.gov/genbank/ |
| Idiomarina loihiensis L2TR | Bacteria | Proteobacteria | genome | 2628 | NCBI | https://www.ncbi.nlm.nih.gov/genbank/ |
| Klebsiella pneumoniae MGH_78578 | Bacteria | Proteobacteria | genome | 5185 | NCBI | https://www.ncbi.nlm.nih.gov/genbank/ |
| Legionella pneumophila Corby | Bacteria | Proteobacteria | genome | 3206 | NCBI | https://www.ncbi.nlm.nih.gov/genbank/ |
| Mannheimia succiniciproducens MBEL55E | Bacteria | Proteobacteria | genome | 2369 | NCBI | https://www.ncbi.nlm.nih.gov/genbank/ |
| Marinobacter aquaeolei VT8 | Bacteria | Proteobacteria | genome | 4272 | NCBI | https://www.ncbi.nlm.nih.gov/genbank/ |
| Marinomonas sp. MWYL1 | Bacteria | Proteobacteria | genome | 4439 | NCBI | https://www.ncbi.nlm.nih.gov/genbank/ |
| Methylococcus capsulatus Bath | Bacteria | Proteobacteria | genome | 2956 | NCBI | https://www.ncbi.nlm.nih.gov/genbank/ |
| Nitrosococcus oceani ATCC_19707 | Bacteria | Proteobacteria | genome | 3017 | NCBI | https://www.ncbi.nlm.nih.gov/genbank/ |
| Pasteurella multocida Pm70 | Bacteria | Proteobacteria | genome | 2015 | NCBI | https://www.ncbi.nlm.nih.gov/genbank/ |
| Pectobacterium atrosepticum SCRI1043 | Bacteria | Proteobacteria | genome | 4472 | NCBI | https://www.ncbi.nlm.nih.gov/genbank/ |
| Photobacterium profundum SS9 | Bacteria | Proteobacteria | genome | 5489 | NCBI | https://www.ncbi.nlm.nih.gov/genbank/ |
| Photorhabdus luminescens TTO1 | Bacteria | Proteobacteria | genome | 4683 | NCBI | https://www.ncbi.nlm.nih.gov/genbank/ |
| Proteus mirabilis HI4320 | Bacteria | Proteobacteria | genome | 3662 | NCBI | https://www.ncbi.nlm.nih.gov/genbank/ |
| Pseudoalteromonas atlantica T6c | Bacteria | Proteobacteria | genome | 4281 | NCBI | https://www.ncbi.nlm.nih.gov/genbank/ |
| Pseudomonas aeruginosa PAO1 | Bacteria | Proteobacteria | genome | 5566 | NCBI | https://www.ncbi.nlm.nih.gov/genbank/ |
| Psychrobacter arcticus 273_4 | Bacteria | Proteobacteria | genome | 2120 | NCBI | https://www.ncbi.nlm.nih.gov/genbank/ |
| Psychromonas ingrahamii 37 | Bacteria | Proteobacteria | genome | 3545 | NCBI | https://www.ncbi.nlm.nih.gov/genbank/ |
| Saccharophagus degradans 2_40 | Bacteria | Proteobacteria | genome | 4007 | NCBI | https://www.ncbi.nlm.nih.gov/genbank/ |
| Salmonella enterica arizonae | Bacteria | Proteobacteria | genome | 4498 | NCBI | https://www.ncbi.nlm.nih.gov/genbank/ |
| Serratia proteamaculans 568 | Bacteria | Proteobacteria | genome | 4942 | NCBI | https://www.ncbi.nlm.nih.gov/genbank/ |
| Shewanella amazonensis SB2B | Bacteria | Proteobacteria | genome | 3645 | NCBI | https://www.ncbi.nlm.nih.gov/genbank/ |
| Shigella boydii Sb227 | Bacteria | Proteobacteria | genome | 4282 | NCBI | https://www.ncbi.nlm.nih.gov/genbank/ |
| Sodalis glossinidius | Bacteria | Proteobacteria | genome | 2715 | NCBI | https://www.ncbi.nlm.nih.gov/genbank/ |
| Stenotrophomonas maltophilia K279a | Bacteria | Proteobacteria | genome | 4386 | NCBI | https://www.ncbi.nlm.nih.gov/genbank/ |
| Thioalkalivibrio sp. HL-EbGR7 | Bacteria | Proteobacteria | genome | 3283 | NCBI | https://www.ncbi.nlm.nih.gov/genbank/ |
| Thiomicrospira crunogena XCL_2 | Bacteria | Proteobacteria | genome | 2196 | NCBI | https://www.ncbi.nlm.nih.gov/genbank/ |
| Vibrio cholerae N16961 | Bacteria | Proteobacteria | genome | 3835 | NCBI | https://www.ncbi.nlm.nih.gov/genbank/ |
| Wigglesworthia glossinidia | Bacteria | Proteobacteria | genome | 617 | NCBI | https://www.ncbi.nlm.nih.gov/genbank/ |
| Xanthomonas campestris ATCC_33913 | Bacteria | Proteobacteria | genome | 4181 | NCBI | https://www.ncbi.nlm.nih.gov/genbank/ |
| Xylella fastidiosa 9a5c | Bacteria | Proteobacteria | genome | 2832 | NCBI | https://www.ncbi.nlm.nih.gov/genbank/ |
| Yersinia enterocolitica 8081 | Bacteria | Proteobacteria | genome | 4051 | NCBI | https://www.ncbi.nlm.nih.gov/genbank/ |
| Caulobacter crescentus CB15 | Bacteria | Proteobacteria | genome | 3737 | NCBI | https://www.ncbi.nlm.nih.gov/genbank/ |
| Phenylobacterium zucineum HLK1 | Bacteria | Proteobacteria | genome | 3854 | NCBI | https://www.ncbi.nlm.nih.gov/genbank/ |
| Agrobacterium tumefaciens C58 | Bacteria | Proteobacteria | genome | 5360 | NCBI | https://www.ncbi.nlm.nih.gov/genbank/ |
| Azorhizobium caulinodans ORS_571 | Bacteria | Proteobacteria | genome | 4717 | NCBI | https://www.ncbi.nlm.nih.gov/genbank/ |
| Bartonella bacilliformis KC583 | Bacteria | Proteobacteria | genome | 1283 | NCBI | https://www.ncbi.nlm.nih.gov/genbank/ |
| Beijerinckia indica subsp. Indica ATCC 9039 | Bacteria | Proteobacteria | genome | 3784 | NCBI | https://www.ncbi.nlm.nih.gov/genbank/ |
| Bradyrhizobium sp. BTAi1 | Bacteria | Proteobacteria | genome | 7622 | NCBI | https://www.ncbi.nlm.nih.gov/genbank/ |
| Brucella abortus 9_941 | Bacteria | Proteobacteria | genome | 3085 | NCBI | https://www.ncbi.nlm.nih.gov/genbank/ |
| Mesorhizobium sp. BNC1 | Bacteria | Proteobacteria | genome | 4543 | NCBI | https://www.ncbi.nlm.nih.gov/genbank/ |
| Methylobacterium extorquens PA1 | Bacteria | Proteobacteria | genome | 4829 | NCBI | https://www.ncbi.nlm.nih.gov/genbank/ |
| Methylocella silvestris BL2 | Bacteria | Proteobacteria | genome | 3818 | NCBI | https://www.ncbi.nlm.nih.gov/genbank/ |
| Ochrobactrum anthropi ATCC_49188 | Bacteria | Proteobacteria | genome | 4799 | NCBI | https://www.ncbi.nlm.nih.gov/genbank/ |
| Oligotropha carboxidovorans OM5 | Bacteria | Proteobacteria | genome | 3847 | NCBI | https://www.ncbi.nlm.nih.gov/genbank/ |
| Parvibaculum lavamentivorans DS1 | Bacteria | Proteobacteria | genome | 3636 | NCBI | https://www.ncbi.nlm.nih.gov/genbank/ |
| Rhizobium etli CFN_42 | Bacteria | Proteobacteria | genome | 5963 | NCBI | https://www.ncbi.nlm.nih.gov/genbank/ |
| Rhodopseudomonas palustris BisA53 | Bacteria | Proteobacteria | genome | 4878 | NCBI | https://www.ncbi.nlm.nih.gov/genbank/ |
| Sinorhizobium medicae WSM419 | Bacteria | Proteobacteria | genome | 6213 | NCBI | https://www.ncbi.nlm.nih.gov/genbank/ |
| Xanthobacter autotrophicus Py2 | Bacteria | Proteobacteria | genome | 5035 | NCBI | https://www.ncbi.nlm.nih.gov/genbank/ |
| Hyphomonas neptunium ATCC_15444 | Bacteria | Proteobacteria | genome | 3505 | NCBI | https://www.ncbi.nlm.nih.gov/genbank/ |
| Nitrobacter hamburgensis X14 | Bacteria | Proteobacteria | genome | 4326 | NCBI | https://www.ncbi.nlm.nih.gov/genbank/ |
| Magnetococcus sp. MC_1 | Bacteria | Proteobacteria | genome | 3716 | NCBI | https://www.ncbi.nlm.nih.gov/genbank/ |
| Maricaulis maris MCS10 | Bacteria | Proteobacteria | genome | 3063 | NCBI | https://www.ncbi.nlm.nih.gov/genbank/ |
| Pelagibacter ubique HTCC1062 | Bacteria | Proteobacteria | genome | 1354 | NCBI | https://www.ncbi.nlm.nih.gov/genbank/ |
| Dinoroseobacter shibae DFL_12 | Bacteria | Proteobacteria | genome | 4187 | NCBI | https://www.ncbi.nlm.nih.gov/genbank/ |
| Jannaschia sp. CCS1 | Bacteria | Proteobacteria | genome | 4283 | NCBI | https://www.ncbi.nlm.nih.gov/genbank/ |
| Paracoccus denitrificans PD1222 | Bacteria | Proteobacteria | genome | 5077 | NCBI | https://www.ncbi.nlm.nih.gov/genbank/ |
| Rhodobacter sphaeroides 2_4_1 | Bacteria | Proteobacteria | genome | 4242 | NCBI | https://www.ncbi.nlm.nih.gov/genbank/ |
| Roseobacter denitrificans OCh_114 | Bacteria | Proteobacteria | genome | 4129 | NCBI | https://www.ncbi.nlm.nih.gov/genbank/ |
| Ruegeria sp. TM1040 | Bacteria | Proteobacteria | genome | 3864 | NCBI | https://www.ncbi.nlm.nih.gov/genbank/ |
| Silicibacter pomeroyi DSS_3 | Bacteria | Proteobacteria | genome | 4252 | NCBI | https://www.ncbi.nlm.nih.gov/genbank/ |
| Acidiphilium cryptum JF_5 | Bacteria | Proteobacteria | genome | 3559 | NCBI | https://www.ncbi.nlm.nih.gov/genbank/ |
| Gluconacetobacter diazotrophicus PAI_5 | Bacteria | Proteobacteria | genome | 7353 | NCBI | https://www.ncbi.nlm.nih.gov/genbank/ |
| Granulibacter bethesdensis CGDNIH1 | Bacteria | Proteobacteria | genome | 2437 | NCBI | https://www.ncbi.nlm.nih.gov/genbank/ |
| Magnetospirillum magneticum AMB_1 | Bacteria | Proteobacteria | genome | 4559 | NCBI | https://www.ncbi.nlm.nih.gov/genbank/ |
| Rhodospirillum rubrum ATCC 11170 | Bacteria | Proteobacteria | genome | 3841 | NCBI | https://www.ncbi.nlm.nih.gov/genbank/ |
| Anaplasma marginale StMaries | Bacteria | Proteobacteria | genome | 948 | NCBI | https://www.ncbi.nlm.nih.gov/genbank/ |
| Ehrlichia canis Jake | Bacteria | Proteobacteria | genome | 925 | NCBI | https://www.ncbi.nlm.nih.gov/genbank/ |

| Species | Domain | Taxonomy | Asembly type | N. proteins | Assembly source | Source link |
|---|---|---|---|---|---|---|
| Neorickettsia sennetsu Miyayama | Bacteria | Proteobacteria | genome | 932 | NCBI | https://www.ncbi.nlm.nih.gov/genbank/ |
| Wolbachia endosymbiont TRS | Bacteria | Proteobacteria | genome | 805 | NCBI | https://www.ncbi.nlm.nih.gov/genbank/ |
| Rickettsia akari Hartford | Bacteria | Proteobacteria | genome | 1259 | NCBI | https://www.ncbi.nlm.nih.gov/genbank/ |
| Erythrobacter litoralis HTCC2594 | Bacteria | Proteobacteria | genome | 3011 | NCBI | https://www.ncbi.nlm.nih.gov/genbank/ |
| Novosphingobium aromaticivorans DSM_12444 | Bacteria | Proteobacteria | genome | 3937 | NCBI | https://www.ncbi.nlm.nih.gov/genbank/ |
| Sphingomonas wittichii RW1 | Bacteria | Proteobacteria | genome | 5345 | NCBI | https://www.ncbi.nlm.nih.gov/genbank/ |
| Sphingopyxis alaskensis RB2256 | Bacteria | Proteobacteria | genome | 3195 | NCBI | https://www.ncbi.nlm.nih.gov/genbank/ |
| Zymomonas mobilis ZM4 | Bacteria | Proteobacteria | genome | 2041 | NCBI | https://www.ncbi.nlm.nih.gov/genbank/ |
| Aromatoleum aromaticum EbN1 | Bacteria | Proteobacteria | genome | 4590 | NCBI | https://www.ncbi.nlm.nih.gov/genbank/ |
| Azoarcus sp. BH72 | Bacteria | Proteobacteria | genome | 3989 | NCBI | https://www.ncbi.nlm.nih.gov/genbank/ |
| Bordetella bronchiseptica RB50 | Bacteria | Proteobacteria | genome | 4994 | NCBI | https://www.ncbi.nlm.nih.gov/genbank/ |
| Burkholderia sp. 383 | Bacteria | Proteobacteria | genome | 7717 | NCBI | https://www.ncbi.nlm.nih.gov/genbank/ |
| Chromobacterium violaceum ATCC_12472 | Bacteria | Proteobacteria | genome | 4407 | NCBI | https://www.ncbi.nlm.nih.gov/genbank/ |
| Cupriavidus taiwanensis | Bacteria | Proteobacteria | genome | 5899 | NCBI | https://www.ncbi.nlm.nih.gov/genbank/ |
| Dechloromonas aromatica RCB | Bacteria | Proteobacteria | genome | 4171 | NCBI | https://www.ncbi.nlm.nih.gov/genbank/ |
| Delftia acidovorans SPH_1 | Bacteria | Proteobacteria | genome | 6040 | NCBI | https://www.ncbi.nlm.nih.gov/genbank/ |
| Diaphorobacter sp. TPSY | Bacteria | Proteobacteria | genome | 3479 | NCBI | https://www.ncbi.nlm.nih.gov/genbank/ |
| Herminiimonas arsenicoxydans | Bacteria | Proteobacteria | genome | 3295 | NCBI | https://www.ncbi.nlm.nih.gov/genbank/ |
| Janthinobacterium sp. Marseille | Bacteria | Proteobacteria | genome | 3697 | NCBI | https://www.ncbi.nlm.nih.gov/genbank/ |
| Leptothrix cholodnii SP-6 | Bacteria | Proteobacteria | genome | 4363 | NCBI | https://www.ncbi.nlm.nih.gov/genbank/ |
| Methylibium petroleiphilum PM1 | Bacteria | Proteobacteria | genome | 4449 | NCBI | https://www.ncbi.nlm.nih.gov/genbank/ |
| Methylobacillus flagellatus KT | Bacteria | Proteobacteria | genome | 2753 | NCBI | https://www.ncbi.nlm.nih.gov/genbank/ |
| Neisseria gonorrhoeae FA_1090 | Bacteria | Proteobacteria | genome | 2002 | NCBI | https://www.ncbi.nlm.nih.gov/genbank/ |
| Nitrosomonas europaea ATCC_19718 | Bacteria | Proteobacteria | genome | 2461 | NCBI | https://www.ncbi.nlm.nih.gov/genbank/ |
| Nitrosospira multiformis ATCC_25196 | Bacteria | Proteobacteria | genome | 2805 | NCBI | https://www.ncbi.nlm.nih.gov/genbank/ |
| Polaromonas sp. JS666 | Bacteria | Proteobacteria | genome | 5453 | NCBI | https://www.ncbi.nlm.nih.gov/genbank/ |
| Polynucleobacter sp. QLW P1DMWA 1 | Bacteria | Proteobacteria | genome | 2077 | NCBI | https://www.ncbi.nlm.nih.gov/genbank/ |
| Ralstonia eutropha H16 | Bacteria | Proteobacteria | genome | 6626 | NCBI | https://www.ncbi.nlm.nih.gov/genbank/ |
| Rhodoferax ferrireducens T118 | Bacteria | Proteobacteria | genome | 4418 | NCBI | https://www.ncbi.nlm.nih.gov/genbank/ |
| Thiobacillus denitrificans ATCC_25259 | Bacteria | Proteobacteria | genome | 2827 | NCBI | https://www.ncbi.nlm.nih.gov/genbank/ |
| Verminephrobacter eiseniae EF01_2 | Bacteria | Proteobacteria | genome | 4947 | NCBI | https://www.ncbi.nlm.nih.gov/genbank/ |
| Arcobacter butzleri RM4018 | Bacteria | Proteobacteria | genome | 2259 | NCBI | https://www.ncbi.nlm.nih.gov/genbank/ |
| Bdellovibrio bacteriovorus HD100 | Bacteria | Proteobacteria | genome | 3587 | NCBI | https://www.ncbi.nlm.nih.gov/genbank/ |
| Campylobacter concisus 13826 | Bacteria | Proteobacteria | genome | 1985 | NCBI | https://www.ncbi.nlm.nih.gov/genbank/ |
| Desulfococcus oleovorans Hxd3 | Bacteria | Proteobacteria | genome | 3265 | NCBI | https://www.ncbi.nlm.nih.gov/genbank/ |
| Desulfotalea psychrophila LSv54 | Bacteria | Proteobacteria | genome | 3234 | NCBI | https://www.ncbi.nlm.nih.gov/genbank/ |
| Desulfovibrio desulfuricans G20 | Bacteria | Proteobacteria | genome | 3775 | NCBI | https://www.ncbi.nlm.nih.gov/genbank/ |
| Desulfovibrio vulgaris DP4 | Bacteria | Proteobacteria | genome | 3091 | NCBI | https://www.ncbi.nlm.nih.gov/genbank/ |
| Desulfovibrio vulgaris Hildenborough | Bacteria | Proteobacteria | genome | 3531 | NCBI | https://www.ncbi.nlm.nih.gov/genbank/ |
| Geobacter metallireducens GS_15 | Bacteria | Proteobacteria | genome | 3532 | NCBI | https://www.ncbi.nlm.nih.gov/genbank/ |
| Geobacter sulfurreducens PCA | Bacteria | Proteobacteria | genome | 3445 | NCBI | https://www.ncbi.nlm.nih.gov/genbank/ |
| Geobacter uraniumreducens Rf4 | Bacteria | Proteobacteria | genome | 4357 | NCBI | https://www.ncbi.nlm.nih.gov/genbank/ |
| Helicobacter acinonychis Sheeba | Bacteria | Proteobacteria | genome | 1618 | NCBI | https://www.ncbi.nlm.nih.gov/genbank/ |
| Lawsonia intracellularis PHE_MN1_00 | Bacteria | Proteobacteria | genome | 1337 | NCBI | https://www.ncbi.nlm.nih.gov/genbank/ |
| Nautilia profundicola AmH | Bacteria | Proteobacteria | genome | 1730 | NCBI | https://www.ncbi.nlm.nih.gov/genbank/ |
| Nitratiruptor sp. SB155_2 | Bacteria | Proteobacteria | genome | 1843 | NCBI | https://www.ncbi.nlm.nih.gov/genbank/ |
| Pelobacter carbinolicus DSM_2380 | Bacteria | Proteobacteria | genome | 3352 | NCBI | https://www.ncbi.nlm.nih.gov/genbank/ |
| Pelobacter propionicus DSM_2379 | Bacteria | Proteobacteria | genome | 3804 | NCBI | https://www.ncbi.nlm.nih.gov/genbank/ |
| Sulfurimonas denitrificans DSM 1251 | Bacteria | Proteobacteria | genome | 2096 | NCBI | https://www.ncbi.nlm.nih.gov/genbank/ |
| Sulfurovum sp. NBC37_1 | Bacteria | Proteobacteria | genome | 2438 | NCBI | https://www.ncbi.nlm.nih.gov/genbank/ |
| Syntrophobacter fumaroxidans MPOB | Bacteria | Proteobacteria | genome | 4064 | NCBI | https://www.ncbi.nlm.nih.gov/genbank/ |
| Syntrophus aciditrophicus SB | Bacteria | Proteobacteria | genome | 3168 | NCBI | https://www.ncbi.nlm.nih.gov/genbank/ |
| Wolinella succinogenes DSM_1740 | Bacteria | Proteobacteria | genome | 2042 | NCBI | https://www.ncbi.nlm.nih.gov/genbank/ |
| Chlamydia muridarum Nigg | Bacteria | Chlamydiae | genome | 911 | NCBI | https://www.ncbi.nlm.nih.gov/genbank/ |
| Chlamydia trachomatis 70 | Bacteria | Chlamydiae | genome | 923 | NCBI | https://www.ncbi.nlm.nih.gov/genbank/ |
| Chlamydophila abortus S26_3 | Bacteria | Chlamydiae | genome | 932 | NCBI | https://www.ncbi.nlm.nih.gov/genbank/ |
| Chlamydophila pneumoniae CWL029 | Bacteria | Chlamydiae | genome | 1052 | NCBI | https://www.ncbi.nlm.nih.gov/genbank/ |
| Waddlia chondrophila WSU 86-1044 | Bacteria | Chlamydiae | genome | 1956 | NCBI | https://www.ncbi.nlm.nih.gov/genbank/ |
| Criblamydia sequanensis CRIB-18 | Bacteria | Chlamydiae | genome | 2418 | NCBI | https://www.ncbi.nlm.nih.gov/genbank/ |
| Parachlamydia acanthamoebae | Bacteria | Chlamydiae | genome | 2809 | NCBI | https://www.ncbi.nlm.nih.gov/genbank/ |
| Protochlamydia amoebophila UWE25 | Bacteria | Chlamydiae | genome | 2031 | NCBI | https://www.ncbi.nlm.nih.gov/genbank/ |
| Simkania negevensis Z | Bacteria | Chlamydiae | genome | 2512 | NCBI | https://www.ncbi.nlm.nih.gov/genbank/ |
| Rhodopirellula baltica SH_1 | Bacteria | Planctomycetes | genome | 7325 | NCBI | https://www.ncbi.nlm.nih.gov/genbank/ |
| Akkermansia muciniphila ATCC BAA-835 | Bacteria | Verrucomicrobia | genome | 2138 | NCBI | https://www.ncbi.nlm.nih.gov/genbank/ |
| Opitutus terrae PB90-1 | Bacteria | Verrucomicrobia | genome | 4612 | NCBI | https://www.ncbi.nlm.nih.gov/genbank/ |
| Borrelia afzelii PKo | Bacteria | Spirochaeta | genome | 1214 | NCBI | https://www.ncbi.nlm.nih.gov/genbank/ |
| Brachyspira hyodysenteriae WA1 | Bacteria | Spirochaeta | genome | 2642 | NCBI | https://www.ncbi.nlm.nih.gov/genbank/ |
| Leptospira borgpetersenii JB197 | Bacteria | Spirochaeta | genome | 2880 | NCBI | https://www.ncbi.nlm.nih.gov/genbank/ |
| Treponema denticola ATCC_35405 | Bacteria | Spirochaeta | genome | 2767 | NCBI | https://www.ncbi.nlm.nih.gov/genbank/ |
| Fervidobacterium nodosum Rt17_B1 | Bacteria | Thermotogales | genome | 1750 | NCBI | https://www.ncbi.nlm.nih.gov/genbank/ |
| Petrotoga mobilis SJ95 | Bacteria | Thermotogales | genome | 1898 | NCBI | https://www.ncbi.nlm.nih.gov/genbank/ |
| Thermosipho melanesiensis BI429 | Bacteria | Thermotogales | genome | 1879 | NCBI | https://www.ncbi.nlm.nih.gov/genbank/ |
| Thermotoga lettingae TMO | Bacteria | Thermotogales | genome | 2040 | NCBI | https://www.ncbi.nlm.nih.gov/genbank/ |

| Species | Domain | Taxonomy | Asembly type | N. proteins | Assembly source | Source link |
|---|---|---|---|---|---|---|
| Babesia bigemina | Eukaryota | Alveolata\|Apicomplexa | genome | 5079 | NCBI | https://www.ncbi.nlm.nih.gov/genbank/ |
| Babesia microti | Eukaryota | Alveolata\|Apicomplexa | genome | 3492 | NCBI | https://www.ncbi.nlm.nih.gov/genbank/ |
| Cryptosporidium hominis TU502 | Eukaryota | Alveolata\|Apicomplexa | genome | 3885 | NCBI | https://www.ncbi.nlm.nih.gov/genbank/ |
| Cryptosporidium parvum | Eukaryota | Alveolata\|Apicomplexa | genome | 3805 | NCBI | https://www.ncbi.nlm.nih.gov/genbank/ |
| Eimeria mitis | Eukaryota | Alveolata\|Apicomplexa | genome | 8748 | NCBI | https://www.ncbi.nlm.nih.gov/genbank/ |
| Gregarina niphandrodes | Eukaryota | Alveolata\|Apicomplexa | genome | 6375 | NCBI | https://www.ncbi.nlm.nih.gov/genbank/ |
| Neospora caninum | Eukaryota | Alveolata\|Apicomplexa | genome | 6936 | NCBI | https://www.ncbi.nlm.nih.gov/genbank/ |
| Plasmodium falciparum 3D7 | Eukaryota | Alveolata\|Apicomplexa | genome | 5262 | NCBI | https://www.ncbi.nlm.nih.gov/genbank/ |
| Plasmodium vivax North Korea | Eukaryota | Alveolata\|Apicomplexa | genome | 6664 | NCBI | https://www.ncbi.nlm.nih.gov/genbank/ |
| Plasmodium yoelii 17XNL | Eukaryota | Alveolata\|Apicomplexa | genome | 7353 | NCBI | https://www.ncbi.nlm.nih.gov/genbank/ |
| Theileria orientalis Shintoku | Eukaryota | Alveolata\|Apicomplexa | genome | 3995 | NCBI | https://www.ncbi.nlm.nih.gov/genbank/ |
| Theileria parva Muguga | Eukaryota | Alveolata\|Apicomplexa | genome | 4061 | NCBI | https://www.ncbi.nlm.nih.gov/genbank/ |
| Toxoplasma gondii GT1 | Eukaryota | Alveolata\|Apicomplexa | genome | 8102 | Autre | https://toxodb.org/toxo/app/ |
| Toxoplasma gondii GT1 | Eukaryota | Alveolata\|Apicomplexa | plastid genome | 26 | NCBI | https://www.ncbi.nlm.nih.gov/genbank/ |
| Anophryoides haemophila AH6 | Eukaryota | Alveolata\|Ciliophora | transcriptome | 423 | MMETSP | https://www.imicrobe.us/#/projects/104 |
| Condylostoma magnum COL2 | Eukaryota | Alveolata\|Ciliophora | transcriptome | 2247 | MMETSP | https://www.imicrobe.us/#/projects/104 |
| Euplotes focardii TN1 | Eukaryota | Alveolata\|Ciliophora | transcriptome | 12634 | MMETSP | https://www.imicrobe.us/#/projects/104 |
| Litonotus pictus P1 | Eukaryota | Alveolata\|Ciliophora | transcriptome | 13248 | MMETSP | https://www.imicrobe.us/#/projects/104 |
| Paramecium tetraurelia | Eukaryota | Alveolata\|Ciliophora | genome | 39571 | Autre | https://www.genoscope.cns.fr/paramecium/ |
| Tetrahymena thermophila | Eukaryota | Alveolata\|Ciliophora | genome | 24725 | Autre | https://tet.ciliate.org/ |
| Chromera velia | Eukaryota | Alveolata\|Colpodellida | plastid genome | 80 | NCBI | https://www.ncbi.nlm.nih.gov/genbank/ |
| Chromera velia | Eukaryota | Alveolata\|Colpodellida | transcriptome | 19244 | MMETSP | https://www.imicrobe.us/#/projects/104 |
| Vitrella brassicaformis CCMP3155 | Eukaryota | Alveolata\|Colpodellida | plastid genome | 81 | NCBI | https://www.ncbi.nlm.nih.gov/genbank/ |
| Vitrella brassicaformis CCMP3346 | Eukaryota | Alveolata\|Colpodellida | transcriptome | 23743 | MMETSP | https://www.imicrobe.us/#/projects/104 |
| Perkinsus marinus | Eukaryota | Alveolata\|Perkinsozoa | genome | 23654 | NCBI | https://www.ncbi.nlm.nih.gov/genbank/ |
| Acanthamoeba castellanii str Neff | Eukaryota | Amoebozoa | genome | 15014 | NCBI | https://www.ncbi.nlm.nih.gov/genbank/ |
| Dictyostelium discoideum | Eukaryota | Amoebozoa | genome | 13425 | Dyctibase | https://www.ncbi.nlm.nih.gov/genbank/ |
| Dictyostelium purpureum QSDP1 | Eukaryota | Amoebozoa | genome | 12410 | JGI | https://mycocosm.jgi.doe.gov/ |
| Entamoeba dispar SAW760 | Eukaryota | Amoebozoa | genome | 8811 | NCBI | https://www.ncbi.nlm.nih.gov/genbank/ |
| Entamoeba histolytica | Eukaryota | Amoebozoa | genome | 9772 | NCBI | https://www.ncbi.nlm.nih.gov/genbank/ |
| Entamoeba invadens IP1 | Eukaryota | Amoebozoa | genome | 11997 | NCBI | https://www.ncbi.nlm.nih.gov/genbank/ |
| Entamoeba nuttalli P19 | Eukaryota | Amoebozoa | genome | 6187 | NCBI | https://www.ncbi.nlm.nih.gov/genbank/ |
| Filamoeba nolandi | Eukaryota | Amoebozoa | transcriptome | 19578 | MMETSP | https://www.imicrobe.us/#/projects/104 |
| Mayorella sp. | Eukaryota | Amoebozoa | transcriptome | 10394 | MMETSP | https://www.imicrobe.us/#/projects/104 |
| Neoparamoeba aestuarina | Eukaryota | Amoebozoa | transcriptome | 18898 | MMETSP | https://www.imicrobe.us/#/projects/104 |
| Paramoeba atlantica | Eukaryota | Amoebozoa | transcriptome | 15740 | MMETSP | https://www.imicrobe.us/#/projects/104 |
| Pessonella sp. | Eukaryota | Amoebozoa | transcriptome | 14148 | MMETSP | https://www.imicrobe.us/#/projects/104 |
| Polysphondylium pallidum | Eukaryota | Amoebozoa | genome | 12409 | NCBI | https://www.ncbi.nlm.nih.gov/genbank/ |
| Stereomyxa ramosa | Eukaryota | Amoebozoa | transcriptome | 15014 | MMETSP | https://www.imicrobe.us/#/projects/104 |
| Trichosphaerium sp. | Eukaryota | Amoebozoa | transcriptome | 15840 | MMETSP | https://www.imicrobe.us/#/projects/104 |
| Vannella robusta | Eukaryota | Amoebozoa | transcriptome | 14576 | MMETSP | https://www.imicrobe.us/#/projects/104 |
| Vannella sp. | Eukaryota | Amoebozoa | transcriptome | 16925 | MMETSP | https://www.imicrobe.us/#/projects/104 |
| Vexillifera sp. | Eukaryota | Amoebozoa | transcriptome | 9684 | MMETSP | https://www.imicrobe.us/#/projects/104 |
| Bigelowiella natans | Eukaryota | Chlorarachniophyta | genome | 21708 | JGI | https://mycocosm.jgi.doe.gov/ |
| Bigelowiella natans | Eukaryota | Chlorarachniophyta | nucleomorph genome | 283 | NCBI | https://www.ncbi.nlm.nih.gov/genbank/ |
| Bigelowiella natans | Eukaryota | Chlorarachniophyta | plastid genome | 61 | NCBI | https://www.ncbi.nlm.nih.gov/genbank/ |
| Chlorarachnion reptans CCCM449 | Eukaryota | Chlorarachniophyta | transcriptome | 21543 | MMETSP | https://www.imicrobe.us/#/projects/104 |
| Gymnochlora sp. CCMP2014 | Eukaryota | Chlorarachniophyta | transcriptome | 15507 | MMETSP | https://www.imicrobe.us/#/projects/104 |
| Gymnochlora stellata | Eukaryota | Chlorarachniophyta | plastid genome | 60 | NCBI | https://www.ncbi.nlm.nih.gov/genbank/ |
| Lotharella amoebiformis | Eukaryota | Chlorarachniophyta | transcriptome | 18612 | MMETSP | https://www.imicrobe.us/#/projects/104 |
| Lotharella globosa | Eukaryota | Chlorarachniophyta | transcriptome | 21225 | MMETSP | https://www.imicrobe.us/#/projects/104 |
| Lotharella oceanica | Eukaryota | Chlorarachniophyta | nucleomorph genome | 608 | NCBI | https://www.ncbi.nlm.nih.gov/genbank/ |
| Lotharella oceanica | Eukaryota | Chlorarachniophyta | plastid genome | 60 | NCBI | https://www.ncbi.nlm.nih.gov/genbank/ |
| Lotharella vacuolata | Eukaryota | Chlorarachniophyta | plastid genome | 59 | NCBI | https://www.ncbi.nlm.nih.gov/genbank/ |
| Norrisiella sphaerica BC52 | Eukaryota | Chlorarachniophyta | transcriptome | 14550 | MMETSP | https://www.imicrobe.us/#/projects/104 |
| Partenskyella glossopodia RCC365 | Eukaryota | Chlorarachniophyta | transcriptome | 15025 | MMETSP | https://www.imicrobe.us/#/projects/104 |
| Partenskyella glossopodia RCC365 | Eukaryota | Chlorarachniophyta | plastid genome | 61 | NCBI | https://www.ncbi.nlm.nih.gov/genbank/ |
| Capsaspora owczarzaki | Eukaryota | Choanoflagellata | genome | 8381 | JGI | https://mycocosm.jgi.doe.gov/ |
| Monosiga brevicollis | Eukaryota | Choanoflagellata | genome | 9203 | NCBI | https://www.ncbi.nlm.nih.gov/genbank/ |
| Thecamonas trahens | Eukaryota | Choanoflagellata | genome | 11582 | JGI | https://mycocosm.jgi.doe.gov/ |
| Palpitomonas bilix NIES-2562 | Eukaryota | Cryptista | transcriptome | 17940 | MMETSP | https://www.imicrobe.us/#/projects/104 |
| Chroomonas mesostigmatica CCMP1168 | Eukaryota | Cryptophyta | transcriptome | 36883 | MMETSP | https://www.imicrobe.us/#/projects/104 |
| Chroomonas mesostigmatica CCMP1168 | Eukaryota | Cryptophyta | nucleomorph genome | 505 | NCBI | https://www.ncbi.nlm.nih.gov/genbank/ |
| Chroomonas placoidea | Eukaryota | Cryptophyta | plastid genome | 147 | NCBI | https://www.ncbi.nlm.nih.gov/genbank/ |
| Cryptomonas curvata | Eukaryota | Cryptophyta | plastid genome | 146 | NCBI | https://www.ncbi.nlm.nih.gov/genbank/ |
| Cryptomonas curvata | Eukaryota | Cryptophyta | transcriptome | 21670 | NCBI | https://www.ncbi.nlm.nih.gov/genbank/ |
| Cryptomonas paramecium CCAP977 | Eukaryota | Cryptophyta | transcriptome | 34554 | MMETSP | https://www.imicrobe.us/#/projects/104 |
| Cryptomonas paramecium CCAP977 | Eukaryota | Cryptophyta | plastid genome | 82 | NCBI | https://www.ncbi.nlm.nih.gov/genbank/ |
| Cryptomonas paramecium CCAP977 | Eukaryota | Cryptophyta | nucleomorph genome | 466 | NCBI | https://www.ncbi.nlm.nih.gov/genbank/ |
| Geminigera cryophila CCMP2564 | Eukaryota | Cryptophyta | transcriptome | 45232 | MMETSP | https://www.imicrobe.us/#/projects/104 |
| Goniomonas sp. | Eukaryota | Cryptophyta | transcriptome | 15726 | MMETSP | https://www.imicrobe.us/#/projects/104 |
| Guillardia theta | Eukaryota | Cryptophyta | genome | 24867 | JGI | https://mycocosm.jgi.doe.gov/ |
| Guillardia theta | Eukaryota | Cryptophyta | nucleomorph genome | 485 | NCBI | https://www.ncbi.nlm.nih.gov/genbank/ |

| Species | Domain | Taxonomy | Asembly type | N. proteins | Assembly source | Source link |
|---|---|---|---|---|---|---|
| Guillardia theta | Eukaryota | Cryptophyta | plastid genome | 147 | NCBI | https://www.ncbi.nlm.nih.gov/genbank/ |
| Hanusia phi CCMP325 | Eukaryota | Cryptophyta | transcriptome | 20820 | MMETSP | https://www.imicrobe.us/#/projects/104 |
| Hemiselmis andersenii CCMP439 | Eukaryota | Cryptophyta | transcriptome | 24741 | MMETSP | https://www.imicrobe.us/#/projects/104 |
| Hemiselmis andersenii CCMP439 | Eukaryota | Cryptophyta | nucleomorph genome | 471 | NCBI | https://www.ncbi.nlm.nih.gov/genbank/ |
| Hemiselmis rufescens | Eukaryota | Cryptophyta | transcriptome | 9800 | NCBI | https://www.ncbi.nlm.nih.gov/genbank/ |
| Hemiselmis tepida | Eukaryota | Cryptophyta | transcriptome | 16614 | NCBI | https://www.ncbi.nlm.nih.gov/genbank/ |
| Proteomonas sulcata CCMP704 | Eukaryota | Cryptophyta | transcriptome | 21553 | MMETSP | https://www.imicrobe.us/#/projects/104 |
| Rhodomonas abbreviata | Eukaryota | Cryptophyta | transcriptome | 30145 | MMETSP | https://www.imicrobe.us/#/projects/104 |
| Rhodomonas salina | Eukaryota | Cryptophyta | plastid genome | 146 | NCBI | https://www.ncbi.nlm.nih.gov/genbank/ |
| Teleaulax amphioxeia | Eukaryota | Cryptophyta | plastid genome | 143 | NCBI | https://www.ncbi.nlm.nih.gov/genbank/ |
| Bodo saltans | Eukaryota | Euglenozoa | EST | 78 | NCBI | https://www.ncbi.nlm.nih.gov/genbank/ |
| Cryptoglena skujai | Eukaryota | Euglenozoa | plastid genome | 66 | NCBI | https://www.ncbi.nlm.nih.gov/genbank/ |
| Euglena gracilis | Eukaryota | Euglenozoa | EST | 215 | NCBI | https://www.ncbi.nlm.nih.gov/genbank/ |
| Euglena gracilis | Eukaryota | Euglenozoa | plastid genome | 67 | NCBI | https://www.ncbi.nlm.nih.gov/genbank/ |
| Euglena gracilis | Eukaryota | Euglenozoa | transcriptome | 14856 | Home Made | |
| Euglena longa | Eukaryota | Euglenozoa | plastid genome | 46 | NCBI | https://www.ncbi.nlm.nih.gov/genbank/ |
| Euglena sp. | Eukaryota | Euglenozoa | transcriptome | 10407 | 1KP | https://db.cngb.org/onekp/ |
| Euglenaformis proxima | Eukaryota | Euglenozoa | plastid genome | 63 | NCBI | https://www.ncbi.nlm.nih.gov/genbank/ |
| Euglenaria anabaena | Eukaryota | Euglenozoa | plastid genome | 67 | NCBI | https://www.ncbi.nlm.nih.gov/genbank/ |
| Eutreptiella gymnastica | Eukaryota | Euglenozoa | transcriptome | 19125 | MMETSP | https://www.imicrobe.us/#/projects/104 |
| Eutreptiella gymnastica | Eukaryota | Euglenozoa | plastid genome | 62 | NCBI | https://www.ncbi.nlm.nih.gov/genbank/ |
| Leishmania infantum | Eukaryota | Euglenozoa | genome | 7992 | NCBI | https://www.ncbi.nlm.nih.gov/genbank/ |
| Leishmania major strain Friedlin | Eukaryota | Euglenozoa | genome | 8266 | NCBI | https://www.ncbi.nlm.nih.gov/genbank/ |
| Monomorphina aenigmatica | Eukaryota | Euglenozoa | plastid genome | 62 | NCBI | https://www.ncbi.nlm.nih.gov/genbank/ |
| Neobodo designis CCAP1951 | Eukaryota | Euglenozoa | transcriptome | 29005 | MMETSP | https://www.imicrobe.us/#/projects/104 |
| Trachelomonas volvocina | Eukaryota | Euglenozoa | plastid genome | 64 | NCBI | https://www.ncbi.nlm.nih.gov/genbank/ |
| Trypanosoma brucei TREU927 | Eukaryota | Euglenozoa | genome | 8712 | NCBI | https://www.ncbi.nlm.nih.gov/genbank/ |
| Trypanosoma cruzi strain CL Brener | Eukaryota | Euglenozoa | genome | 19608 | NCBI | https://www.ncbi.nlm.nih.gov/genbank/ |
| Giardia lamblia | Eukaryota | Excavata | genome | 6499 | giardiadb | https://giardiadb.org/giardiadb/app |
| Naegleria gruberi | Eukaryota | Excavata | genome | 15753 | JGI | https://mycocosm.jgi.doe.gov/ |
| Percolomonas cosmopolitus AE-1 | Eukaryota | Excavata | transcriptome | 8191 | MMETSP | https://www.imicrobe.us/#/projects/104 |
| Stygamoeba regulata BSH-02190019 | Eukaryota | Excavata | transcriptome | 30471 | MMETSP | https://www.imicrobe.us/#/projects/104 |
| Allomyces macrogynus | Eukaryota | Fungi | genome | 17600 | JGI | https://mycocosm.jgi.doe.gov/ |
| Aspergillus niger | Eukaryota | Fungi | genome | 11197 | JGI | https://mycocosm.jgi.doe.gov/ |
| Batrachochytrium dendrobatidis JAM81 | Eukaryota | Fungi | genome | 8732 | JGI | https://mycocosm.jgi.doe.gov/ |
| Botrytis cinerea | Eukaryota | Fungi | genome | 16448 | JGI | https://mycocosm.jgi.doe.gov/ |
| Chaetomium globosum | Eukaryota | Fungi | genome | 11124 | JGI | https://mycocosm.jgi.doe.gov/ |
| Coprinopsis cinerea | Eukaryota | Fungi | genome | 13342 | JGI | https://mycocosm.jgi.doe.gov/ |
| Cryptococcus neoformans JEC21 | Eukaryota | Fungi | genome | 6594 | NCBI | https://www.ncbi.nlm.nih.gov/genbank/ |
| Debaryomyces hansenii CBS767 | Eukaryota | Fungi | genome | 6317 | NCBI | https://www.ncbi.nlm.nih.gov/genbank/ |
| Fusarium graminearum | Eukaryota | Fungi | genome | 13321 | NCBI | https://www.ncbi.nlm.nih.gov/genbank/ |
| Histoplasma capsulatum | Eukaryota | Fungi | genome | 9532 | NCBI | https://www.ncbi.nlm.nih.gov/genbank/ |
| Laccaria bicolor | Eukaryota | Fungi | genome | 20614 | JGI | https://mycocosm.jgi.doe.gov/ |
| Melampsora larici-populina | Eukaryota | Fungi | genome | 16831 | NCBI | https://www.ncbi.nlm.nih.gov/genbank/ |
| Mucor circinelloides | Eukaryota | Fungi | genome | 10930 | JGI | https://mycocosm.jgi.doe.gov/ |
| Mycosphaerella graminicola | Eukaryota | Fungi | genome | 11395 | NCBI | https://www.ncbi.nlm.nih.gov/genbank/ |
| Neurospora crassa OR74A | Eukaryota | Fungi | genome | 9841 | NCBI | https://www.ncbi.nlm.nih.gov/genbank/ |
| Phanerochaete chrisosporium | Eukaryota | Fungi | genome | 10048 | JGI | https://mycocosm.jgi.doe.gov/ |
| Phycomyces blakesleeanus | Eukaryota | Fungi | genome | 14792 | NCBI | https://www.ncbi.nlm.nih.gov/genbank/ |
| Postia placenta | Eukaryota | Fungi | genome | 9113 | JGI | https://mycocosm.jgi.doe.gov/ |
| Puccinia graminis | Eukaryota | Fungi | genome | 20566 | JGI | https://mycocosm.jgi.doe.gov/ |
| Rhizopus oryzae | Eukaryota | Fungi | genome | 17459 | NCBI | https://www.ncbi.nlm.nih.gov/genbank/ |
| Saccharomyces cerevisiae RM11-1a | Eukaryota | Fungi | genome | 5880 | NCBI | https://www.ncbi.nlm.nih.gov/genbank/ |
| Saccharomyces cerevisiae S288C | Eukaryota | Fungi | genome | 5894 | SGD | https://www.yeastgenome.org/ |
| Schizophylum commune | Eukaryota | Fungi | genome | 13181 | JGI | https://mycocosm.jgi.doe.gov/ |
| Schizosaccharomyces pombe | Eukaryota | Fungi | genome | 5003 | NCBI | https://www.ncbi.nlm.nih.gov/genbank/ |
| Spizellomyces punctatus | Eukaryota | Fungi | genome | 8804 | JGI | https://mycocosm.jgi.doe.gov/ |
| Sporobolomyces roseus | Eukaryota | Fungi | genome | 5536 | JGI | https://mycocosm.jgi.doe.gov/ |
| Stagonospora nodorum | Eukaryota | Fungi | genome | 16597 | JGI | https://mycocosm.jgi.doe.gov/ |
| Trichoderma reesei | Eukaryota | Fungi | genome | 9143 | JGI | https://mycocosm.jgi.doe.gov/ |
| Ustilago maydis | Eukaryota | Fungi | genome | 6548 | NCBI | https://www.ncbi.nlm.nih.gov/genbank/ |
| Yarrowia lipolytica | Eukaryota | Fungi | genome | 6472 | NCBI | https://www.ncbi.nlm.nih.gov/genbank/ |
| Cyanophora paradoxa | Eukaryota | Glaucophyta | genome | 32167 | Rutgers | http://cyanophora.rutgers.edu/ |
| Cyanophora paradoxa | Eukaryota | Glaucophyta | EST | 20255 | NCBI | https://www.ncbi.nlm.nih.gov/genbank/ |
| Cyanophora paradoxa | Eukaryota | Glaucophyta | plastid genome | 149 | NCBI | https://www.ncbi.nlm.nih.gov/genbank/ |
| Cyanophora paradoxa | Eukaryota | Glaucophyta | genome | 28 | Rutgers | http://cyanophora.rutgers.edu/ |
| Cyanoptyche gloeocystis | Eukaryota | Glaucophyta | transcriptome | 4903 | 1KP | https://db.cngb.org/onekp/ |
| Glaucocystis nostochinearum | Eukaryota | Glaucophyta | transcriptome | 15415 | 1KP | https://db.cngb.org/onekp/ |
| Glaucocystis nostochinearum | Eukaryota | Glaucophyta | transcriptome | 3611 | NCBI | https://www.ncbi.nlm.nih.gov/genbank/ |
| Gloeochaete wittrockiana | Eukaryota | Glaucophyta | transcriptome | 17062 | 1KP | https://db.cngb.org/onekp/ |
| Gloeochaete wittrockiana | Eukaryota | Glaucophyta | transcriptome | 11758 | 1KP | https://db.cngb.org/onekp/ |
| Calcidiscus leptoporus RCC1130 | Eukaryota | Haptophyta | transcriptome | 27349 | MMETSP | https://www.imicrobe.us/#/projects/104 |
| Chrysochromulina polylepis CCMP1757 | Eukaryota | Haptophyta | transcriptome | 50016 | MMETSP | https://www.imicrobe.us/#/projects/104 |

| Species | Domain | Taxonomy | Asembly type | N. proteins | Assembly source | Source link |
|---|---|---|---|---|---|---|
| Emiliania huxleyi CCMP1516 | Eukaryota | Haptophyta | genome | 33340 | JGI | https://mycocosm.jgi.doe.gov/ |
| Emiliania huxleyi CCMP1516 | Eukaryota | Haptophyta | plastid genome | 119 | NCBI | https://www.ncbi.nlm.nih.gov/genbank/ |
| Exanthemachrysis gayraliae RCC1523 | Eukaryota | Haptophyta | transcriptome | 17259 | MMETSP | https://www.imicrobe.us/#/projects/104 |
| Florenciella sp. RCC1587 | Eukaryota | Haptophyta | transcriptome | 24779 | MMETSP | https://www.imicrobe.us/#/projects/104 |
| Gephyrocapsa oceanica RCC1303 | Eukaryota | Haptophyta | transcriptome | 35096 | MMETSP | https://www.imicrobe.us/#/projects/104 |
| Isochrysis sp. CCMP1324 | Eukaryota | Haptophyta | transcriptome | 17820 | MMETSP | https://www.imicrobe.us/#/projects/104 |
| Pavlova lutheri | Eukaryota | Haptophyta | plastid genome | 111 | NCBI | https://www.ncbi.nlm.nih.gov/genbank/ |
| Pavlova sp. CCMP459 | Eukaryota | Haptophyta | transcriptome | 17723 | MMETSP | https://www.imicrobe.us/#/projects/104 |
| Pleurochrysis carterae CCMP645 | Eukaryota | Haptophyta | transcriptome | 30387 | MMETSP | https://www.imicrobe.us/#/projects/104 |
| Prymnesium parvum Texoma1 | Eukaryota | Haptophyta | transcriptome | 36946 | MMETSP | https://www.imicrobe.us/#/projects/104 |
| Amphimedon queenslandica | Eukaryota | Metazoa | genome | 43615 | JGI | https://mycocosm.jgi.doe.gov/ |
| Caenorhabditis elegans | Eukaryota | Metazoa | genome | 23906 | NCBI | https://www.ncbi.nlm.nih.gov/genbank/ |
| Daphnia pulex | Eukaryota | Metazoa | genome | 30940 | JGI | https://mycocosm.jgi.doe.gov/ |
| Helobdella robusta | Eukaryota | Metazoa | genome | 23432 | JGI | https://mycocosm.jgi.doe.gov/ |
| Homo sapiens | Eukaryota | Metazoa | genome | 45366 | NCBI | https://www.ncbi.nlm.nih.gov/genbank/ |
| Nematostella vectensis | Eukaryota | Metazoa | genome | 27273 | JGI | https://mycocosm.jgi.doe.gov/ |
| Trichoplax adhaerens Grell-BS-1999 | Eukaryota | Metazoa | genome | 11520 | JGI | https://mycocosm.jgi.doe.gov/ |
| Mitosporidium daphniae | Eukaryota | Microsporidia | genome | 3330 | NCBI | https://www.ncbi.nlm.nih.gov/genbank/ |
| Rozella allomycis CSF55 | Eukaryota | Microsporidia | genome | 6350 | JGI | https://mycocosm.jgi.doe.gov/ |
| Ahnfeltiopsis flabelliformis | Eukaryota | Rhodophyceae | transcriptome | 4120 | 1KP | https://db.cngb.org/onekp/ |
| Bangia atropurpurea | Eukaryota | Rhodophyceae | plastid genome | 206 | NCBI | https://www.ncbi.nlm.nih.gov/genbank/ |
| Bangiopsis subsimplex | Eukaryota | Rhodophyceae | plastid genome | 198 | NCBI | https://www.ncbi.nlm.nih.gov/genbank/ |
| Betaphycus philippinensis | Eukaryota | Rhodophyceae | transcriptome | 4063 | 1KP | https://db.cngb.org/onekp/ |
| Boldia erythrosiphon | Eukaryota | Rhodophyceae | plastid genome | 187 | NCBI | https://www.ncbi.nlm.nih.gov/genbank/ |
| Bulboplastis apyrenoidosa | Eukaryota | Rhodophyceae | plastid genome | 171 | NCBI | https://www.ncbi.nlm.nih.gov/genbank/ |
| Calliarthron tuberculosum | Eukaryota | Rhodophyceae | genome | 23966 | Rutgers | http://cyanophora.rutgers.edu/ |
| Calliarthron tuberculosum | Eukaryota | Rhodophyceae | plastid genome | 201 | NCBI | https://www.ncbi.nlm.nih.gov/genbank/ |
| Ceramium kondoi | Eukaryota | Rhodophyceae | transcriptome | 3782 | 1KP | https://db.cngb.org/onekp/ |
| Chondrus crispus | Eukaryota | Rhodophyceae | genome | 10406 | NCBI | https://www.ncbi.nlm.nih.gov/genbank/ |
| Chondrus crispus | Eukaryota | Rhodophyceae | EST | 3227 | NCBI | https://www.ncbi.nlm.nih.gov/genbank/ |
| Chondrus crispus | Eukaryota | Rhodophyceae | plastid genome | 204 | NCBI | https://www.ncbi.nlm.nih.gov/genbank/ |
| Chroodactylon ornatum | Eukaryota | Rhodophyceae | transcriptome | 6836 | 1KP | https://db.cngb.org/onekp/ |
| Compsopogon coeruleus | Eukaryota | Rhodophyceae | transcriptome | 6844 | MMETSP | https://www.imicrobe.us/#/projects/104 |
| Compsopogon coeruleus | Eukaryota | Rhodophyceae | plastid genome | 195 | NCBI | https://www.ncbi.nlm.nih.gov/genbank/ |
| Corynoplastis japonica NIES-2662 | Eukaryota | Rhodophyceae | plastid genome | 169 | NCBI | https://www.ncbi.nlm.nih.gov/genbank/ |
| Cyanidioschyzon merolae | Eukaryota | Rhodophyceae | genome | 4764 | NCBI | https://www.ncbi.nlm.nih.gov/genbank/ |
| Cyanidioschyzon merolae | Eukaryota | Rhodophyceae | plastid genome | 207 | NCBI | https://www.ncbi.nlm.nih.gov/genbank/ |
| Cyanidium caldarium | Eukaryota | Rhodophyceae | plastid genome | 197 | NCBI | https://www.ncbi.nlm.nih.gov/genbank/ |
| Dumontia simplex | Eukaryota | Rhodophyceae | transcriptome | 4141 | 1KP | https://db.cngb.org/onekp/ |
| Erythrolobus australicus CCMP3124 | Eukaryota | Rhodophyceae | transcriptome | 7264 | MMETSP | https://www.imicrobe.us/#/projects/104 |
| Erythrolobus madagascarensis CCMP3276 | Eukaryota | Rhodophyceae | transcriptome | 7364 | MMETSP | https://www.imicrobe.us/#/projects/104 |
| Eucheuma denticulatum | Eukaryota | Rhodophyceae | EST | 3826 | Rutgers | http://cyanophora.rutgers.edu/ |
| Flintiella sanguinaria | Eukaryota | Rhodophyceae | plastid genome | 186 | NCBI | https://www.ncbi.nlm.nih.gov/genbank/ |
| Furcellaria lumbricalis | Eukaryota | Rhodophyceae | EST | 1131 | NCBI | https://www.ncbi.nlm.nih.gov/genbank/ |
| Galdieria sulphuraria | Eukaryota | Rhodophyceae | genome | 6623 | MSU | http://genomics.msu.edu/galdieria |
| Gelidium elegans | Eukaryota | Rhodophyceae | plastid genome | 200 | NCBI | https://www.ncbi.nlm.nih.gov/genbank/ |
| Gelidium vagum | Eukaryota | Rhodophyceae | plastid genome | 200 | NCBI | https://www.ncbi.nlm.nih.gov/genbank/ |
| Glaucosphaera vacuolata | Eukaryota | Rhodophyceae | transcriptome | 11499 | 1KP | https://db.cngb.org/onekp/ |
| Gloiopeltis furcata | Eukaryota | Rhodophyceae | transcriptome | 3947 | 1KP | https://db.cngb.org/onekp/ |
| Gracilaria changii | Eukaryota | Rhodophyceae | EST | 3800 | NCBI | https://www.ncbi.nlm.nih.gov/genbank/ |
| Gracilaria chilensis | Eukaryota | Rhodophyceae | plastid genome | 202 | NCBI | https://www.ncbi.nlm.nih.gov/genbank/ |
| Gracilaria chouae | Eukaryota | Rhodophyceae | transcriptome | 3572 | 1KP | https://db.cngb.org/onekp/ |
| Gracilaria salicornia | Eukaryota | Rhodophyceae | plastid genome | 202 | NCBI | https://www.ncbi.nlm.nih.gov/genbank/ |
| Gracilaria tenuistipitata | Eukaryota | Rhodophyceae | plastid genome | 203 | NCBI | https://www.ncbi.nlm.nih.gov/genbank/ |
| Gracilariopsis lemaneiformis | Eukaryota | Rhodophyceae | plastid genome | 203 | NCBI | https://www.ncbi.nlm.nih.gov/genbank/ |
| Grateloupia filicina | Eukaryota | Rhodophyceae | transcriptome | 19456 | 1KP | https://db.cngb.org/onekp/ |
| Grateloupia taiwanensis | Eukaryota | Rhodophyceae | plastid genome | 233 | NCBI | https://www.ncbi.nlm.nih.gov/genbank/ |
| Griffithsia okiensis | Eukaryota | Rhodophyceae | EST | 717 | NCBI | https://www.ncbi.nlm.nih.gov/genbank/ |
| Heterosiphonia pulchra | Eukaryota | Rhodophyceae | transcriptome | 4518 | 1KP | https://db.cngb.org/onekp/ |
| Madagascaria erythrocladiodes CCMP3234 | Eukaryota | Rhodophyceae | transcriptome | 32290 | MMETSP | https://www.imicrobe.us/#/projects/104 |
| Mazzaella japonica | Eukaryota | Rhodophyceae | transcriptome | 4449 | 1KP | https://db.cngb.org/onekp/ |
| Neosiphonia japonica | Eukaryota | Rhodophyceae | transcriptome | 3905 | 1KP | https://db.cngb.org/onekp/ |
| Porphyra pulchra | Eukaryota | Rhodophyceae | plastid genome | 207 | NCBI | https://www.ncbi.nlm.nih.gov/genbank/ |
| Porphyra purpurea | Eukaryota | Rhodophyceae | plastid genome | 209 | NCBI | https://www.ncbi.nlm.nih.gov/genbank/ |
| Porphyra sp. | Eukaryota | Rhodophyceae | EST | 13080 | NCBI | https://www.ncbi.nlm.nih.gov/genbank/ |
| Porphyra yezoensis | Eukaryota | Rhodophyceae | plastid genome | 209 | NCBI | https://www.ncbi.nlm.nih.gov/genbank/ |
| Porphyridium cruentum | Eukaryota | Rhodophyceae | EST | 17827 | Rutgers | http://cyanophora.rutgers.edu/ |
| Porphyridium purpureum | Eukaryota | Rhodophyceae | plastid genome | 224 | NCBI | https://www.ncbi.nlm.nih.gov/genbank/ |
| Pyropia haitanensis | Eukaryota | Rhodophyceae | plastid genome | 211 | NCBI | https://www.ncbi.nlm.nih.gov/genbank/ |
| Pyropia perforata | Eukaryota | Rhodophyceae | plastid genome | 209 | NCBI | https://www.ncbi.nlm.nih.gov/genbank/ |
| Rhodella maculata CCMP736 | Eukaryota | Rhodophyceae | transcriptome | 6080 | MMTESP | https://www.imicrobe.us/#/projects/104 |
| Rhodella violacea | Eukaryota | Rhodophyceae | transcriptome | 4475 | 1KP | https://db.cngb.org/onekp/ |
| Rhodochaete parvula | Eukaryota | Rhodophyceae | transcriptome | 3816 | 1KP | https://db.cngb.org/onekp/ |

| Species | Domain | Taxonomy | Asembly type | N. proteins | Assembly source | Source link |
|---|---|---|---|---|---|---|
| Rhodochaete parvula | Eukaryota | Rhodophyceae | plastid genome | 192 | NCBI | https://www.ncbi.nlm.nih.gov/genbank/ |
| Rhodosorus marinus CCMP769 | Eukaryota | Rhodophyceae | transcriptome | 8169 | MMETSP | https://www.imicrobe.us/#/projects/104 |
| Sheathia arcuata | Eukaryota | Rhodophyceae | plastid genome | 185 | NCBI | https://www.ncbi.nlm.nih.gov/genbank/ |
| Sporolithon durum | Eukaryota | Rhodophyceae | plastid genome | 203 | NCBI | https://www.ncbi.nlm.nih.gov/genbank/ |
| Thorea hispida | Eukaryota | Rhodophyceae | plastid genome | 193 | NCBI | https://www.ncbi.nlm.nih.gov/genbank/ |
| Timspurckia oligopyrenoides CCMP3278 | Eukaryota | Rhodophyceae | transcriptome | 5797 | MMETSP | https://www.imicrobe.us/#/projects/104 |
| Vertebrata lanosa | Eukaryota | Rhodophyceae | plastid genome | 194 | NCBI | https://www.ncbi.nlm.nih.gov/genbank/ |
| Aplanochytrium sp. PBS07 | Eukaryota | Stramenopiles | transcriptome | 9294 | MMETSP | https://www.imicrobe.us/#/projects/104 |
| Aplanochytrium stocchinoi GSBS06 | Eukaryota | Stramenopiles | transcriptome | 10896 | MMETSP | https://www.imicrobe.us/#/projects/104 |
| Asterionella formosa | Eukaryota | Stramenopiles | plastid genome | 134 | NCBI | https://www.ncbi.nlm.nih.gov/genbank/ |
| Asterionellopsis glacialis | Eukaryota | Stramenopiles | plastid genome | 145 | NCBI | https://www.ncbi.nlm.nih.gov/genbank/ |
| Aurantiochytrium limacinum ATCCMYA1381 | Eukaryota | Stramenopiles | transcriptome | 12583 | MMETSP | https://www.imicrobe.us/#/projects/104 |
| Aureococcus anophagefferens | Eukaryota | Stramenopiles | genome | 11501 | JGI | https://mycocosm.jgi.doe.gov/ |
| Aureococcus anophagefferens | Eukaryota | Stramenopiles | plastid genome | 105 | NCBI | https://www.ncbi.nlm.nih.gov/genbank/ |
| Aureoumbra lagunensis CCMP1507 | Eukaryota | Stramenopiles | plastid genome | 110 | NCBI | https://www.ncbi.nlm.nih.gov/genbank/ |
| Aureoumbra lagunensis CCMP1510 | Eukaryota | Stramenopiles | transcriptome | 10284 | MMETSP | https://www.imicrobe.us/#/projects/104 |
| Bicosoecid sp. ms1 | Eukaryota | Stramenopiles | transcriptome | 15066 | MMETSP | https://www.imicrobe.us/#/projects/104 |
| Blastocystis hominis | Eukaryota | Stramenopiles | genome | 6020 | NCBI | https://www.ncbi.nlm.nih.gov/genbank/ |
| Bolidomonas pacifica CCMP1866 | Eukaryota | Stramenopiles | transcriptome | 17780 | MMETSP | https://www.imicrobe.us/#/projects/104 |
| Cafeteria roebergensis E4-10 | Eukaryota | Stramenopiles | transcriptome | 13783 | MMETSP | https://www.imicrobe.us/#/projects/104 |
| Cerataulina daemon | Eukaryota | Stramenopiles | plastid genome | 132 | NCBI | https://www.ncbi.nlm.nih.gov/genbank/ |
| Chaetoceros curvisetus | Eukaryota | Stramenopiles | transcriptome | 18190 | MMETSP | https://www.imicrobe.us/#/projects/104 |
| Chaetoceros neogracile CCMP1317 | Eukaryota | Stramenopiles | transcriptome | 18670 | MMETSP | https://www.imicrobe.us/#/projects/104 |
| Chaetoceros simplex | Eukaryota | Stramenopiles | plastid genome | 131 | NCBI | https://www.ncbi.nlm.nih.gov/genbank/ |
| Chattonella subsalsa CCMP2191 | Eukaryota | Stramenopiles | transcriptome | 18256 | MMETSP | https://www.imicrobe.us/#/projects/104 |
| Corethron pennatum L29A3 | Eukaryota | Stramenopiles | transcriptome | 39296 | MMETSP | https://www.imicrobe.us/#/projects/104 |
| Cylindrotheca closterium | Eukaryota | Stramenopiles | plastid genome | 161 | NCBI | https://www.ncbi.nlm.nih.gov/genbank/ |
| Dictyocha speculum CCMP1381 | Eukaryota | Stramenopiles | transcriptome | 28139 | MMETSP | https://www.imicrobe.us/#/projects/104 |
| Didymosphenia geminata | Eukaryota | Stramenopiles | plastid genome | 130 | NCBI | https://www.ncbi.nlm.nih.gov/genbank/ |
| Dinobryon sp. UTEXLB2267 | Eukaryota | Stramenopiles | transcriptome | 18284 | MMETSP | https://www.imicrobe.us/#/projects/104 |
| Ectocarpus siliculosus | Eukaryota | Stramenopiles | genome | 16254 | JGI | https://mycocosm.jgi.doe.gov/ |
| Ectocarpus siliculosus | Eukaryota | Stramenopiles | plastid genome | 148 | NCBI | https://www.ncbi.nlm.nih.gov/genbank/ |
| Eucampia antarctica CCMP1452 | Eukaryota | Stramenopiles | transcriptome | 10209 | MMETSP | https://www.imicrobe.us/#/projects/104 |
| Eunotia naegelii | Eukaryota | Stramenopiles | plastid genome | 160 | NCBI | https://www.ncbi.nlm.nih.gov/genbank/ |
| Fistulifera solaris | Eukaryota | Stramenopiles | plastid genome | 135 | NCBI | https://www.ncbi.nlm.nih.gov/genbank/ |
| Florenciella parvula RCC1693 | Eukaryota | Stramenopiles | transcriptome | 16685 | MMETSP | https://www.imicrobe.us/#/projects/104 |
| Fragilariopsis cylindrus | Eukaryota | Stramenopiles | genome | 27137 | JGI | https://mycocosm.jgi.doe.gov/ |
| Fragilariopsis kerguelensis L26-C5 | Eukaryota | Stramenopiles | transcriptome | 48137 | MMETSP | https://www.imicrobe.us/#/projects/104 |
| Fucus vesiculosus | Eukaryota | Stramenopiles | plastid genome | 139 | NCBI | https://www.ncbi.nlm.nih.gov/genbank/ |
| Heterosigma akashiwo CCMP2393 | Eukaryota | Stramenopiles | transcriptome | 27903 | MMETSP | https://www.imicrobe.us/#/projects/104 |
| Heterosigma akashiwo NIES293 | Eukaryota | Stramenopiles | plastid genome | 156 | NCBI | https://www.ncbi.nlm.nih.gov/genbank/ |
| Leptocylindrus danicus CCMP1856 | Eukaryota | Stramenopiles | transcriptome | 13968 | MMETSP | https://www.imicrobe.us/#/projects/104 |
| Lithodesmium undulatum | Eukaryota | Stramenopiles | plastid genome | 138 | NCBI | https://www.ncbi.nlm.nih.gov/genbank/ |
| Nannochloropsis gaditana | Eukaryota | Stramenopiles | plastid genome | 119 | NCBI | https://www.ncbi.nlm.nih.gov/genbank/ |
| Nannochloropsis granulata | Eukaryota | Stramenopiles | plastid genome | 125 | NCBI | https://www.ncbi.nlm.nih.gov/genbank/ |
| Ochromonas sp. CCMP1393 | Eukaryota | Stramenopiles | transcriptome | 16703 | MMETSP | https://www.imicrobe.us/#/projects/104 |
| Odontella aurita isolate 1302-5 | Eukaryota | Stramenopiles | transcriptome | 37918 | MMETSP | https://www.imicrobe.us/#/projects/104 |
| Odontella sinensis | Eukaryota | Stramenopiles | plastid genome | 140 | NCBI | https://www.ncbi.nlm.nih.gov/genbank/ |
| Paraphysomonas imperforata PA2 | Eukaryota | Stramenopiles | transcriptome | 15182 | MMETSP | https://www.imicrobe.us/#/projects/104 |
| Pelagococcus subviridis CCMP1429 | Eukaryota | Stramenopiles | transcriptome | 20101 | MMETSP | https://www.imicrobe.us/#/projects/104 |
| Pelagomonas calceolata CCMP1756 | Eukaryota | Stramenopiles | transcriptome | 18094 | MMETSP | https://www.imicrobe.us/#/projects/104 |
| Phaeodactylum tricornutum | Eukaryota | Stramenopiles | genome | 10402 | JGI | https://mycocosm.jgi.doe.gov/ |
| Phaeodactylum tricornutum | Eukaryota | Stramenopiles | plastid genome | 132 | NCBI | https://www.ncbi.nlm.nih.gov/genbank/ |
| Phaeomonas parva CCMP2877 | Eukaryota | Stramenopiles | transcriptome | 19435 | MMETSP | https://www.imicrobe.us/#/projects/104 |
| Phytophthora capsici | Eukaryota | Stramenopiles | genome | 17414 | JGI | https://mycocosm.jgi.doe.gov/ |
| Phytophthora sojae | Eukaryota | Stramenopiles | genome | 19027 | JGI | https://mycocosm.jgi.doe.gov/ |
| Pinguiococcus pyrenoidosus CCMP2078 | Eukaryota | Stramenopiles | transcriptome | 10811 | MMETSP | https://www.imicrobe.us/#/projects/104 |
| Proboscia alata PI-D3 | Eukaryota | Stramenopiles | transcriptome | 25076 | MMETSP | https://www.imicrobe.us/#/projects/104 |
| Proboscia inermis CCAP1064-1 | Eukaryota | Stramenopiles | transcriptome | 17446 | MMETSP | https://www.imicrobe.us/#/projects/104 |
| Pseudopedinella elastica CCMP716 | Eukaryota | Stramenopiles | transcriptome | 39045 | MMETSP | https://www.imicrobe.us/#/projects/104 |
| Pteridomonas danica PT | Eukaryota | Stramenopiles | transcriptome | 20124 | MMETSP | https://www.imicrobe.us/#/projects/104 |
| Rhizosolenia imbricata | Eukaryota | Stramenopiles | plastid genome | 135 | NCBI | https://www.ncbi.nlm.nih.gov/genbank/ |
| Rhizosolenia setigera CCMP 1694 | Eukaryota | Stramenopiles | transcriptome | 21238 | MMETSP | https://www.imicrobe.us/#/projects/104 |
| Roundia cardiophora | Eukaryota | Stramenopiles | plastid genome | 140 | NCBI | https://www.ncbi.nlm.nih.gov/genbank/ |
| Saccharina japonica | Eukaryota | Stramenopiles | plastid genome | 139 | NCBI | https://www.ncbi.nlm.nih.gov/genbank/ |
| Sargassum horneri | Eukaryota | Stramenopiles | plastid genome | 139 | NCBI | https://www.ncbi.nlm.nih.gov/genbank/ |
| Schizochytrium aggregatum ATCC28209 | Eukaryota | Stramenopiles | transcriptome | 13373 | MMETSP | https://www.imicrobe.us/#/projects/104 |
| Striatella unipunctata CCMP2910 | Eukaryota | Stramenopiles | transcriptome | 26607 | MMETSP | https://www.imicrobe.us/#/projects/104 |
| Synchroma pusillum CCMP3072 | Eukaryota | Stramenopiles | transcriptome | 12897 | MMETSP | https://www.imicrobe.us/#/projects/104 |
| Thalassiosira pseudonana | Eukaryota | Stramenopiles | genome | 11776 | JGI | https://mycocosm.jgi.doe.gov/ |
| Thalassiosira pseudonana | Eukaryota | Stramenopiles | plastid genome | 141 | NCBI | https://www.ncbi.nlm.nih.gov/genbank/ |
| Thalassiothrix antarctica L6-D1 | Eukaryota | Stramenopiles | transcriptome | 18200 | MMETSP | https://www.imicrobe.us/#/projects/104 |
| Thraustochytrium sp. LLF1b | Eukaryota | Stramenopiles | transcriptome | 11151 | MMETSP | https://www.imicrobe.us/#/projects/104 |

| Species | Domain | Taxonomy | Asembly type | N. proteins | Assembly source | Source link |
|---|---|---|---|---|---|---|
| Triparma laevis | Eukaryota | Stramenopiles | plastid genome | 141 | NCBI | https://www.ncbi.nlm.nih.gov/genbank/ |
| Ulnaria acus | Eukaryota | Stramenopiles | plastid genome | 130 | NCBI | https://www.ncbi.nlm.nih.gov/genbank/ |
| Undaria pinnatifida | Eukaryota | Stramenopiles | plastid genome | 139 | NCBI | https://www.ncbi.nlm.nih.gov/genbank/ |
| Unidentified eukaryote sp. NY0313808BC1 | Eukaryota | Stramenopiles | transcriptome | 11105 | MMETSP | https://www.imicrobe.us/#/projects/104 |
| Vaucheria litorea CCMP2940 | Eukaryota | Stramenopiles | transcriptome | 8258 | MMETSP | https://www.imicrobe.us/#/projects/104 |
| Vaucheria litorea CCMP2940 | Eukaryota | Stramenopiles | plastid genome | 139 | NCBI | https://www.ncbi.nlm.nih.gov/genbank/ |
| Arabidopsis thaliana | Eukaryota | Chloroplastida | genome | 32615 | TAIR | https://arabidopsis.org/ |
| Arabidopsis thaliana | Eukaryota | Chloroplastida | plastid genome | 85 | NCBI | https://www.ncbi.nlm.nih.gov/genbank/ |
| Bathycoccus prasinos | Eukaryota | Chloroplastida | transcriptome | 5479 | 1KP | https://db.cngb.org/onekp/ |
| Brachypodium distachyon | Eukaryota | Chloroplastida | genome | 32255 | JGI | https://mycocosm.jgi.doe.gov/ |
| Brachypodium distachyon | Eukaryota | Chloroplastida | plastid genome | 81 | NCBI | https://www.ncbi.nlm.nih.gov/genbank/ |
| Bryopsis plumosa | Eukaryota | Chloroplastida | transcriptome | 7322 | 1KP | https://db.cngb.org/onekp/ |
| Cephaleuros virescens | Eukaryota | Chloroplastida | transcriptome | 4495 | 1KP | https://db.cngb.org/onekp/ |
| Chaetopeltis orbicularis | Eukaryota | Chloroplastida | transcriptome | 9074 | 1KP | https://db.cngb.org/onekp/ |
| Chlamydomonas reinhardtii | Eukaryota | Chloroplastida | genome | 16709 | JGI | https://mycocosm.jgi.doe.gov/ |
| Chlamydomonas reinhardtii | Eukaryota | Chloroplastida | plastid genome | 69 | NCBI | https://www.ncbi.nlm.nih.gov/genbank/ |
| Chlorella sp. NC64A | Eukaryota | Chloroplastida | genome | 9791 | JGI | https://mycocosm.jgi.doe.gov/ |
| Chlorella vulgaris C-169 | Eukaryota | Chloroplastida | genome | 9994 | JGI | https://mycocosm.jgi.doe.gov/ |
| Chlorella vulgaris C-169 | Eukaryota | Chloroplastida | plastid genome | 174 | NCBI | https://www.ncbi.nlm.nih.gov/genbank/ |
| Codium fragile | Eukaryota | Chloroplastida | transcriptome | 6868 | 1KP | https://db.cngb.org/onekp/ |
| Crustomastix stigmata CCMP3273 | Eukaryota | Chloroplastida | transcriptome | 17228 | MMETSP | https://www.imicrobe.us/#/projects/104 |
| Dolichomastix tenuilepis CCMP3274 | Eukaryota | Chloroplastida | transcriptome | 17000 | MMETSP | https://www.imicrobe.us/#/projects/104 |
| Dunaliella salina | Eukaryota | Chloroplastida | transcriptome | 11646 | 1KP | https://db.cngb.org/onekp/ |
| Gonium pectorale | Eukaryota | Chloroplastida | transcriptome | 9614 | 1KP | https://db.cngb.org/onekp/ |
| Heterochlamydomonas inaequalis | Eukaryota | Chloroplastida | transcriptome | 9117 | 1KP | https://db.cngb.org/onekp/ |
| Ignatius tetrasporus | Eukaryota | Chloroplastida | transcriptome | 7647 | 1KP | https://db.cngb.org/onekp/ |
| Leptosira obovata | Eukaryota | Chloroplastida | transcriptome | 7570 | 1KP | https://db.cngb.org/onekp/ |
| Lobochlamys segnis | Eukaryota | Chloroplastida | transcriptome | 7468 | 1KP | https://db.cngb.org/onekp/ |
| Lobomonas rostrata | Eukaryota | Chloroplastida | transcriptome | 8230 | 1KP | https://db.cngb.org/onekp/ |
| Micromonas pusilla CCMP1545 | Eukaryota | Chloroplastida | genome | 10547 | JGI | https://mycocosm.jgi.doe.gov/ |
| Micromonas pusilla CCMP1545 | Eukaryota | Chloroplastida | plastid genome | 27 | NCBI | https://www.ncbi.nlm.nih.gov/genbank/ |
| Micromonas sp. RCC299 | Eukaryota | Chloroplastida | plastid genome | 57 | NCBI | https://www.ncbi.nlm.nih.gov/genbank/ |
| Micromonas sp. RCC299 | Eukaryota | Chloroplastida | genome | 10109 | JGI | https://mycocosm.jgi.doe.gov/ |
| Mimulus guttatus | Eukaryota | Chloroplastida | genome | 27501 | JGI | https://mycocosm.jgi.doe.gov/ |
| Nephroselmis olivacea | Eukaryota | Chloroplastida | transcriptome | 10728 | 1KP | https://db.cngb.org/onekp/ |
| Oedogonium cardiacum | Eukaryota | Chloroplastida | transcriptome | 7246 | 1KP | https://db.cngb.org/onekp/ |
| Oryza sativa | Eukaryota | Chloroplastida | genome | 26940 | NCBI | https://www.ncbi.nlm.nih.gov/genbank/ |
| Oryza sativa | Eukaryota | Chloroplastida | plastid genome | 108 | NCBI | https://www.ncbi.nlm.nih.gov/genbank/ |
| Ostreococcus lucimarinus | Eukaryota | Chloroplastida | genome | 7651 | JGI | https://mycocosm.jgi.doe.gov/ |
| Ostreococcus tauri | Eukaryota | Chloroplastida | genome | 7725 | JGI | https://mycocosm.jgi.doe.gov/ |
| Ostreococcus tauri | Eukaryota | Chloroplastida | plastid genome | 61 | NCBI | https://www.ncbi.nlm.nih.gov/genbank/ |
| Parachlorella kessleri | Eukaryota | Chloroplastida | transcriptome | 7510 | 1KP | https://db.cngb.org/onekp/ |
| Phacotus lenticularis | Eukaryota | Chloroplastida | transcriptome | 8348 | 1KP | https://db.cngb.org/onekp/ |
| Physcomitrella patens | Eukaryota | Chloroplastida | genome | 35938 | JGI | https://mycocosm.jgi.doe.gov/ |
| Physcomitrella patens | Eukaryota | Chloroplastida | plastid genome | 85 | NCBI | https://www.ncbi.nlm.nih.gov/genbank/ |
| Pirula salina | Eukaryota | Chloroplastida | transcriptome | 5817 | 1KP | https://db.cngb.org/onekp/ |
| Planophila laetevirens | Eukaryota | Chloroplastida | transcriptome | 6158 | 1KP | https://db.cngb.org/onekp/ |
| Populus trichocarpa | Eukaryota | Chloroplastida | genome | 45555 | JGI | https://mycocosm.jgi.doe.gov/ |
| Populus trichocarpa | Eukaryota | Chloroplastida | plastid genome | 98 | NCBI | https://www.ncbi.nlm.nih.gov/genbank/ |
| Prasinococcus capsulatus CCMP1194 | Eukaryota | Chloroplastida | transcriptome | 8610 | MMETSP | https://www.imicrobe.us/#/projects/104 |
| Prasinoderma coloniale CCMP1413 | Eukaryota | Chloroplastida | transcriptome | 10355 | MMETSP | https://www.imicrobe.us/#/projects/104 |
| Pseudoneochloris marina | Eukaryota | Chloroplastida | transcriptome | 6231 | 1KP | https://db.cngb.org/onekp/ |
| Pyramimonas parkeae CCMP726 | Eukaryota | Chloroplastida | transcriptome | 23234 | MMETSP | https://www.imicrobe.us/#/projects/104 |
| Selaginella moellendorffii | Eukaryota | Chloroplastida | genome | 34697 | JGI | https://mycocosm.jgi.doe.gov/ |
| Selaginella moellendorffii | Eukaryota | Chloroplastida | plastid genome | 70 | NCBI | https://www.ncbi.nlm.nih.gov/genbank/ |
| Spermatozopsis exsultans | Eukaryota | Chloroplastida | transcriptome | 6524 | 1KP | https://db.cngb.org/onekp/ |
| Tetraselmis chuii PLY429 | Eukaryota | Chloroplastida | transcriptome | 23036 | MMETSP | https://www.imicrobe.us/#/projects/104 |
| Trebouxia arboricola | Eukaryota | Chloroplastida | transcriptome | 7032 | 1KP | https://db.cngb.org/onekp/ |
| Trentepohlia annulata | Eukaryota | Chloroplastida | transcriptome | 4705 | 1KP | https://db.cngb.org/onekp/ |
| Uronema belkae | Eukaryota | Chloroplastida | transcriptome | 7496 | 1KP | https://db.cngb.org/onekp/ |
| Volvox carteri | Eukaryota | Chloroplastida | genome | 15669 | JGI | https://mycocosm.jgi.doe.gov/ |
